# A Brain-Wide Map of Neural Activity during Complex Behaviour

**DOI:** 10.1101/2023.07.04.547681

**Authors:** International Brain Laboratory, Brandon Benson, Julius Benson, Daniel Birman, Niccolò Bonacchi, Kcénia Bougrova, Sebastian A Bruijns, Matteo Carandini, Joana A Catarino, Gaelle A Chapuis, Anne K Churchland, Yang Dan, Felicia Davatolhagh, Peter Dayan, Eric EJ DeWitt, Tatiana A Engel, Michele Fabbri, Mayo Faulkner, Ila Rani Fiete, Charles Findling, Laura Freitas-Silva, Berk Gerçek, Kenneth D Harris, Michael Häusser, Sonja B Hofer, Fei Hu, Félix Hubert, Julia M Huntenburg, Anup Khanal, Christopher Krasniak, Christopher Langdon, Petrina Y P Lau, Zachary F Mainen, Guido T Meijer, Nathaniel J Miska, Thomas D Mrsic-Flogel, Jean-Paul Noel, Kai Nylund, Alejandro Pan-Vazquez, Liam Paninski, Alexandre Pouget, Cyrille Rossant, Noam Roth, Rylan Schaeffer, Michael Schartner, Yanliang Shi, Karolina Z Socha, Nicholas A Steinmetz, Karel Svoboda, Anne E Urai, Miles J Wells, Steven Jon West, Matthew R Whiteway, Olivier Winter, Ilana B Witten

## Abstract

A key challenge in neuroscience is understanding how neurons in hundreds of interconnected brain regions integrate sensory inputs with prior expectations to initiate movements and thereby make decisions. It is difficult to meet this challenge if different laboratories apply different analyses to different recordings in different regions during different behaviours. Here, we report a comprehensive set of recordings from 621733 neurons across 139 mice in 12 labs performing a decision-making task with sensory, motor, and cognitive components, obtained with 699 Neuropixels probe insertions covering 279 brain areas in the left forebrain and midbrain and the right hindbrain and cerebellum. We provide an initial appraisal of this brain-wide map, assessing how neural activity encoded key task variables. Representations of visual stimuli appeared transiently in classical visual areas after stimulus onset and then spread to ramp-like activity in a collection of mid- and hindbrain regions that also encoded choices. Neural responses correlated with motor action almost everywhere in the brain. Responses to reward delivery and consumption versus negative feedback were also widespread. This publicly available dataset represents an unprecedented resource for understanding how computations distributed across and within brain areas drive behaviour.

## Introduction

How does coherent and effective output emerge from hundreds of interconnected brain areas processing information related to sensation, decisions, action, and behaviour^1–3?^ To answer this question we need to know how the activity of individual neurons and populations of neurons across the brain reflects variables such as stimuli, expectations, choices, actions, rewards, and punishments^4^. Electrophysiological recordings from animals have been instrumental in this exploration. Until recently, however, technical limitations restricted these recordings to a limited number of brain areas, leaving much of the mammalian brain uncharted or described by fragmentary maps. For instance, the mouse brain comprises over 300 identified regions^5^, of which only a minority has been recorded systematically in comparable behavioural settings. The regions studied were typically chosen based on *a priori* hypotheses derived from previous recordings and anatomical connectivity. This approach can suggest a localisation of function, revealing brain regions as being engaged in computations such as the accumulation of sensory evidence in favour of a decision^6^. Nevertheless, studies have shown that such regions can sometimes be silenced without substantial behavioural consequences^7–11^, suggesting the involvement of other regions. Overall, it has proven difficult to obtain a comprehensive picture of neural processing based on different reports from different laboratories recording in different regions during different behaviours and analysing the data with different methods.

A broader search for the neuronal correlates of variables such as sensation and decision-making thus requires systematic recording at a wider scale, using a single task with sufficient behavioural complexity, and using the same analysis methods on all the data. Obtaining such a comprehensive dataset has recently become possible with advances in recording technology. In a species with a small brain such as the mouse, Neuropixels probes^12^ have enabled larger-scale recordings, such as sampling activity across eight visual areas^13^, or tens of brain regions in mice performing behavioural tasks^14–16^, or experiencing changes in physiological state^17^. Modern imaging techniques also provide a wider view over activity across dorsal cortical regions^18–20^. Results from these broad surveys suggest that the encoding of task variables varies greatly: some neural correlates are found only in few brain areas, others in sparse sets of cells, while other variables appear to be distributed much more broadly. Thus, it is critical to record even more fully, because past recordings may have missed essential regions with focused coding of certain variables, and have not fully characterised the nature of distributed coding.

Here we present a publicly available dataset^21^ of recordings from 699 Neuropixels probe insertions spaced across an entire hemisphere of the brain in mice performing a behavioural task that requires sensory, cognitive, and motor processing^22^. This approach enables detection of brain-wide correlates of sensation, choice, action, and reward, as well as internal cognitive states including stimulus expectation or priors (this “block” prior is described in the companion paper^23^). We also describe initial analyses of these data, which indicate that the neural correlates of some variables, such as reward and action, can be found in many neurons across essentially the whole brain, while those of other variables such as the input stimulus can be decoded from a narrower range of regions, and significantly influence the activity of far fewer individual neurons. These data, which can be examined at viz.internationalbrainlab.org and downloaded from int-brain-lab.github.io/iblenv/notebooks_external/data_release_brainwidemap.html are intended to be the starting point for a detailed examination of decision-making processes across the brain, and represent a unique resource enabling the community to perform a broad range of further analyses with single-neuron resolution at a brain-wide scale.

## Results

First, we describe the task and recording strategy; further details of how we ensured reproducible behaviour, electrophysiology, and videography are available separately^22, 24^ and are summarised in the Methods. Next, we describe the analysis methods we used to provide different views of a rich and complex dataset. Finally, we report the neural correlates of the main events and variables in the task: the visual stimulus, choice, feedback, and wheel movement.

### Behavioural task and recording

We trained 139 mice (94 males, 45 females) on the International Brain Laboratory (IBL) decision-making task^22^ (Fig. 1a,b). On each trial, a visual stimulus appeared to the left or right on a screen, and the mouse had to move it to the centre by turning a wheel with its front paws within 60 seconds (Fig. 1c). The prior probability for the stimulus to appear on the left/right side was constant over a block of trials, at 20/80% (right block) or 80/20% (left block). Blocks lasted for between 20 and 100 trials, drawn from a truncated geometric distribution (mean 51 trials). Block changes were not cued. Stimulus contrast was sampled uniformly from five possible values (100, 25, 12.5, 6.25, and 0%). The 0% contrast trials, when no stimulus was presented, were assigned to a left or right side following the probability distribution of the block, allowing mice to perform above chance by incorporating this prior in their choices. Following a wheel turn, mice received positive feedback in the form of a water reward, or negative feedback in the form of a white noise pulse and a 2 s time-out. The next trial began after a delay, followed by a quiescence period during which the mice had to hold the wheel still.

**Figure 1.**
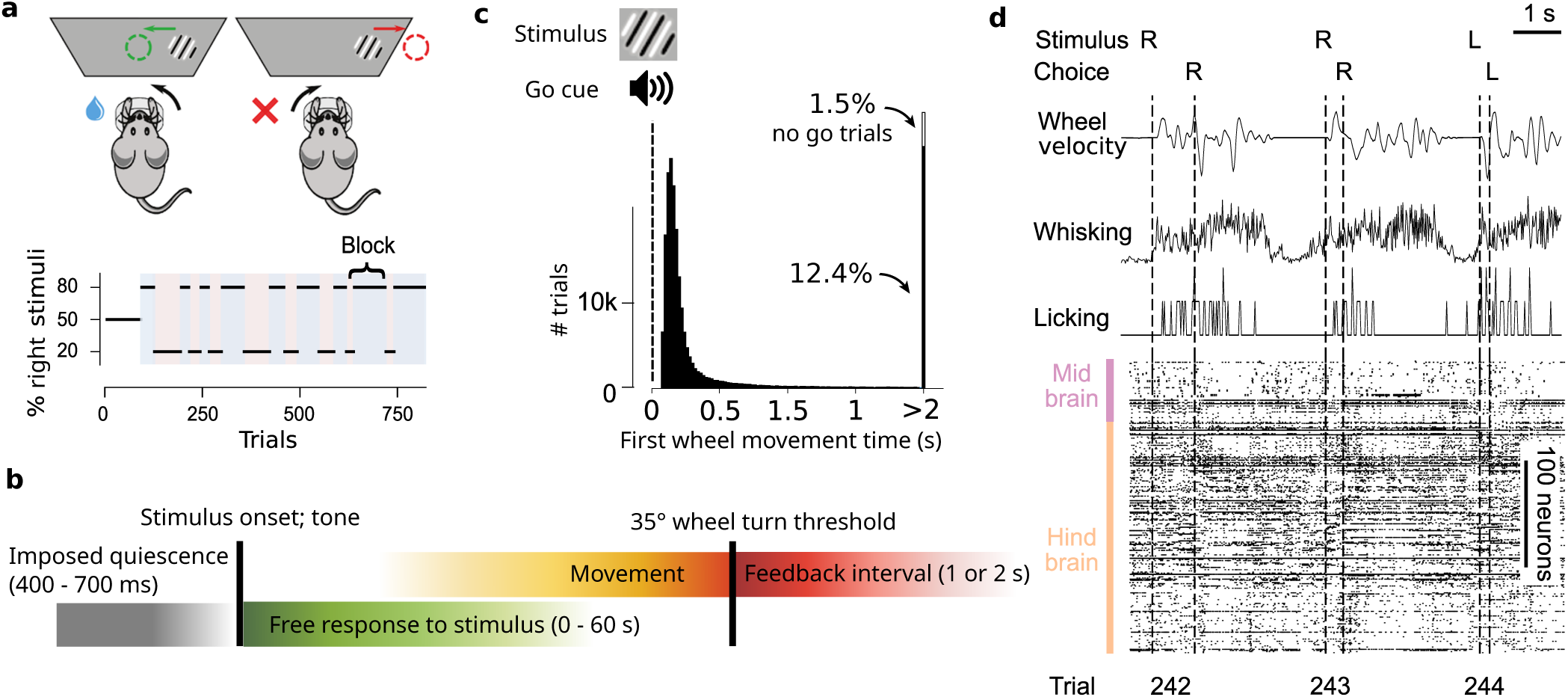
The IBL task and data types. **a)** Schematic of the IBL task and block structure of an example session. After the first 90 trials in a session, the probability of the stimulus being on the right side was varied in blocks of consecutive trials. **b)** Timeline of the IBL task with main events and the variables on which we concentrate. After mice hold the wheel still for an imposed quiescence period, the stimulus appears on the screen, then the first movement is detected by a small threshold of wheel rotation. When the wheel is turned by more than 35*^◦^*, the animal is considered to have made a choice and feedback is provided. Colours are as in later figures. **c)** Distribution of times between stimulus onset and first wheel movement time (which can be interpreted as a reaction time), from 459 sessions. The distribution is truncated at 80 ms and 2 s, because we only analysed trials with first wheel movement times between these bounds. Not depicted in this histogram are 22.8% of first wheel movement times that occur under 80ms. **d)** Example time series and key trial information for three trials, including the wheel’s rotary encoder output, video analysis and rasters showing the spike times of simultaneously recorded neurons in multiple brain regions.

In these mice, we performed 699 Neuropixels probe insertions (see an example of one recording of three trials in Fig. 1d), following a grid covering the left hemisphere of the forebrain and midbrain, which typically represent stimuli or actions on the contralateral side, and the right hemisphere of the cerebellum and hindbrain, which typically represent the ipsilateral side (Fig. 2a). These recordings were collected by 12 labs in Europe and the USA, with most recordings using two simultaneous probe insertions. To ensure reproducibility, one brain location was targeted in every mouse in every laboratory, as described elsewhere^24^. Only sessions with at least 250 trials and behavioural performance of 90% on the 100% contrast trials were retained for further analysis. Data were uploaded to a central server, preprocessed, and shared through a standardised interface^25^. To perform spike-sorting on the recordings, we used a version of Kilosort^26^, with custom additions**^?^**. This yielded 621733 units (including multi-neuron activity), averaging 889 per probe. To separate individual neurons from clusters of multi-neuron activity, we then applied stringent quality control metrics (based on those described in Ref.^13^), which identified 75708 ‘good’ neurons, averaging 108 per probe.

**Figure 2.**
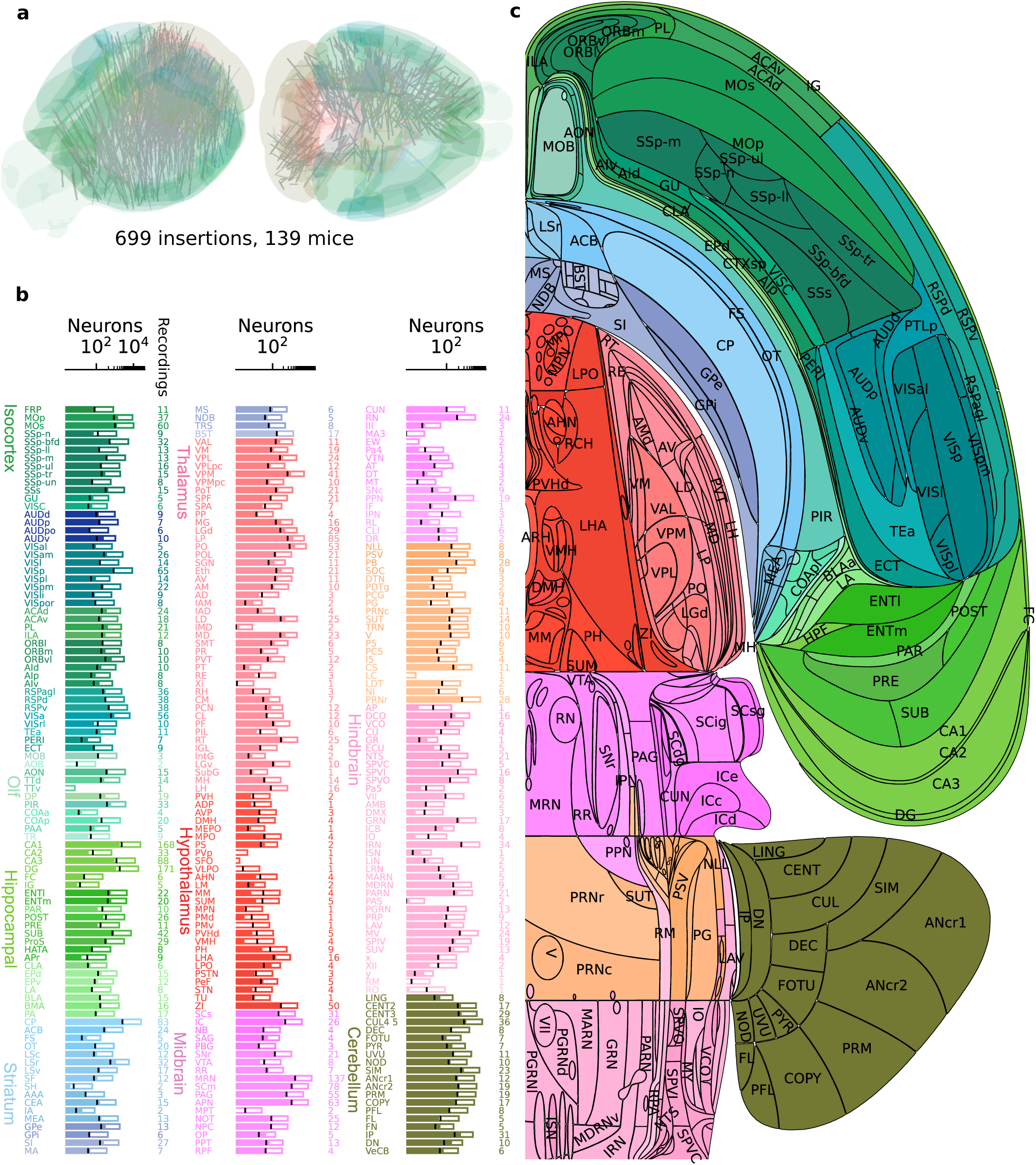
Brain-wide recordings during behaviour. **a)** Neuropixels probe trajectories shown in a 3D brain schematic. **b)** For each region, the number of neurons recorded (full bar length) and the number of good neurons used for analysis (filled portion; for reference, the black line on each bar shows 10% of the number of recorded neurons, which is the average number of neurons that were good). Definitions of the acronyms for brain regions are available in an online table. The same table reports the so-called Cosmos hierarchical grouping of the regions, which distinguishes ‘Isocortex’, ‘OLF’ (olfactory areas), ‘HPF’ (hippocampal formation), ‘CTXsp’ (cortical subplate), ‘CNU’ (cerebral nuclei), ‘TH’ (thalamus), ‘HY’ (hypothalamus), ‘MB’ (midbrain), ‘HB’ (hindbrain), ‘CB’ (cerebellum). **c)** Flatmap of one hemisphere indicating the region acronyms.

Following recordings, probe tracks were reconstructed using serial-section 2-photon microscopy^27^, and each recording site and neuron was assigned a region in the Allen Common Coordinate Framework^5^ (CCF; details in online table; statistics in Fig. 2b). Our main analyses are restricted to regions with 20 or more neurons assigned to them in at least two sessions, with at least 5 neurons per session. Due to the grid-based probe insertion strategy, more recordings were made in larger regions, typically leading to more neurons. Note that it was harder to extract good quality neurons from some regions than others, and so the yield differed substantially. While information about molecular cell types can sometimes be gleaned from spike waveforms, we did not attempt this for the analyses here; for example, while we recorded from some of the main neuromodulatory regions, we do not make specific claims about which neurons release which neurotransmitter.

To illustrate our main results, we plot them into a flatmap of the brain^28^ (Fig. 2c); supplementary figures present some of the results on more conventional 2D sections (Fig. S1). For reference, average activity across all regions aligned to the major task events – stimulus onset, first wheel movement time, and feedback – is shown in Fig. S2. To visualize continuous temporal dynamics across different task epochs, we also computed time-warped average activity, simultaneously aligned to stimulus, movement, and feedback onsets (Fig. S3).

The processed data for each trial consisted of a set of spike trains from multiple brain regions together with continuous behavioural traces and discrete behavioural events (Fig. 1d). These were recorded using a variety of sensors including three video cameras and a rotary encoder on the choice wheel. The data were processed with custom scripts and DeepLabCut^29^ to yield the times of major events in each trial along with wheel velocity, whisker motion energy, lick timing, and the positions of body parts. We only analyzed trials in which the first wheel movement time (which is our operational definition of a reaction time) was between 80 ms and 2 s (Fig. 1c).

Instructions for accessing the data^21^, together with an online browser, are available at https://data.internationalbrainlab.org.

### Behavioural performance

As previously shown^22^, mice learned both to indicate the position of the stimulus and to exploit the block structure of the task. After training, they made correct choices on 81.4% ± 0.4% (mean ± s.d.) of the trials, performing better and faster on trials with high visual stimulus contrast (Fig. 3a). Sessions lasted on average 645 trials (median 602, range [401, 1525]). Towards the end of the sessions, performance decreased and first wheel movement times increased (Fig. 3b,c). On 0% contrast trials, where no visual information was provided, mice gained reward on 58.7% ± 0.4% (mean ± s.d.) of trials, significantly better than chance (t-test *t*_138_ = 20.18*, p* = 5.2*^−^*^43^). After a block switch, mice took around 5 to 10 trials to adjust their behaviour to the new block, as revealed by the fraction of correct choices made on 0% contrast trials after the switch (Fig. 3d). Mice were influenced by their prior estimate also in the presence of visual stimuli: for all contrast values, mice tended to answer right more often on right blocks than left blocks (Fig. 3e).

**Figure 3.**
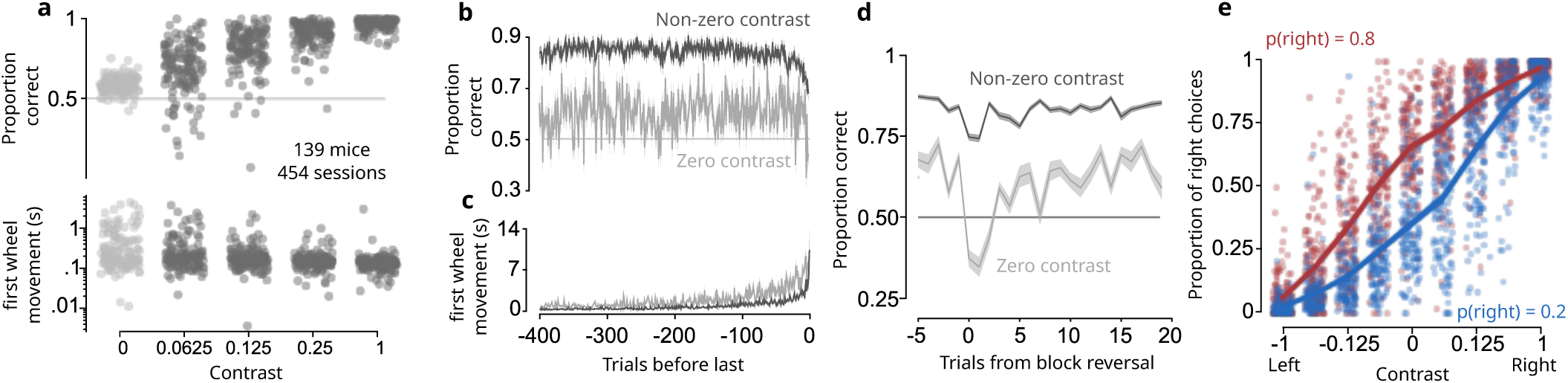
Behaviour during the task. **a)** Proportion of correct choices (top) and first wheel movement time (bottom) as a function of stimulus contrast (one point per mouse per contrast; grey points for 0% contrast; black points for non-zero contrast). Performance on 0% contrast trials (grey) was 58.7% *±* 0.4% (mean *±* s.e. across mice) correct. **b)** Proportion of correct choices as a function of the number of trials before the end of the session for 0% (grey) and non-0% (black) contrast trials (mean *±* s.e. across mice). **c)** The same analysis for first wheel movement times. **d)** Reversal curves: proportion correct around a block change for trials with 0% contrast and *>*0% contrast (mean +/- s.e. across mice). The first 90 trials (when the stimulus was not biased to appear more frequently on a side) were ignored for this analysis. **e)** Psychometric curves showing the fraction of right choices as a function of signed contrast (positive for right stimuli, negative for left stimuli), for all mice (one dot per contrast per mouse). Right choices were more common in right blocks (red), when the stimulus appeared on the right with probability p(right)=0.8, than on left blocks (blue) when p(right)=0.2.

### Neural analyses

To obtain an initial appraisal of the brain-wide map we performed single-cell and population analyses, assessing how neural activity encodes task variables and how it can be analysed to decode these variables. We considered four key task variables (Fig. 4a): visual stimulus, choice (clockwise or counterclockwise turning of the wheel), feedback (reward or time-out), and wheel movement (speed and velocity). The main figures show the results of these analyses in a canonical dataset of 201 regions for which we had at least two sessions with 5 good neurons each and at least 20 good neurons after pooling all sessions (for a total of 62857 neurons; Table 1). Supplementary figures show results for the wider range of neurons and regions appropriate for each analysis.

**Figure 4.**
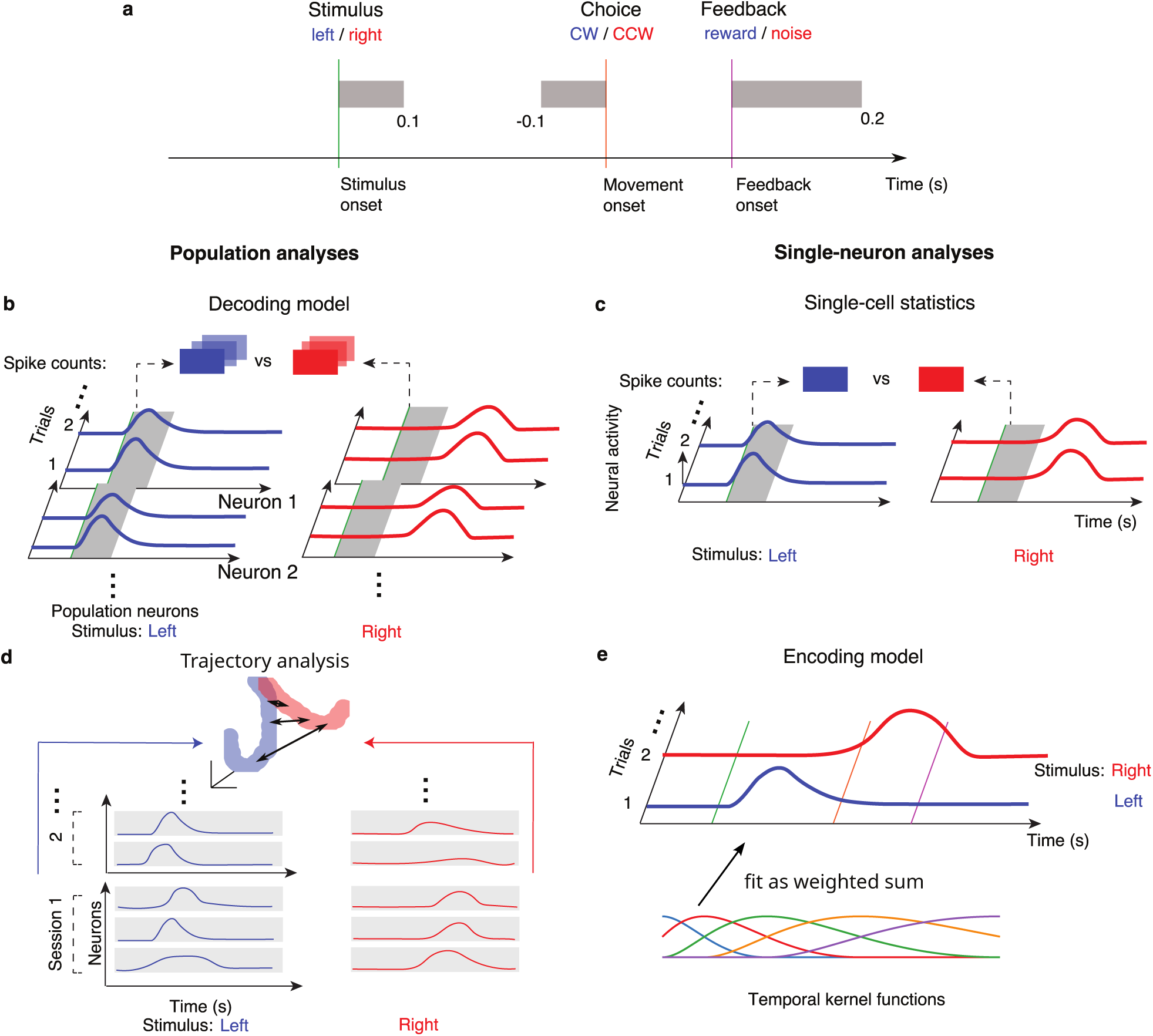
Illustration of neural analyses. See also Fig. S4. **a)** Task structure with time windows used for analysis in grey. **b)** The decoding model quantifies neural population correlates with task variables. **c)** Single-cell analysis quantifies single-cell neural correlates with task variables. **d)** Population trajectory analysis describes the time evolution of the across-session neural population response, pooling cells across all recordings per region. **e)** The encoding model uses multiple linear regression of task- and behaviourally-defined temporal kernels (the multicoloured traces) on the activity of single neurons (shown as the red and blue curves on right and left trials respectively).

**Table 1.** Session and probe filtering. The table indicates the progressive filtering of the sessions and probes based on the various inclusion criteria described in the text. The first row is based on the released dataset, while subsequent rows indicate data constraints associated with the various methods of analysis.

| filter | remaining |  |  |  |
| --- | --- | --- | --- | --- |
|  | sessions | probes | neurons | regions |
| session and insertion QC | 459 | 699 | 621,733 | 281 |
| first wheel movement time & missing events | 459 | 699 | 621,733 | 281 |
| minimum 3 error trials | 459 | 699 | 621,733 | 281 |
| single unit QC | 459 | 698 | 75,708 | 268 |
| gray matter regions | 459 | 698 | 65,336 | 266 |
| minimum 5 units/region | 455 | 691 | 63,357 | 245 |
| minimum 2 sessions/region | 454 | 690 | 62,990 | 210 |
| minimum 20 units/region | 454 | 689 | 62,857 | 201 |

To provide complementary views on how these task variables are represented in each brain region, we used four analysis techniques (Fig. 4b-e; see also Fig. S4 for a fuller picture). The details of each technique are provided in the Methods, along with a discussion of the corresponding null distributions, permutation tests and false discovery rate (FDR*_q_* at a level *q*; using the Benjamini-Hochberg procedure) that we used to limit statistical artefacts.

First, we used a *decoding model* that attempts to predict the value of the task variable on each trial from neural population activity using regularized logistic or linear regression (Fig. 4b; Fig. S4b). This analysis can detect situations in which a variable is encoded robustly, but only in a sparse subset of neurons. We assessed decoding for each variable separately without considering the correlations between the task variables. This quantifies what downstream neurons would be able to determine from the activity, but does not disentangle factors that are related such as stimulus side and choice. We performed decoding within each region, correcting the *R*^2^ of the fit to a variable by the *R*^2^ of the fit to a suitable null distribution and used Fisher’s combined probability test^30, 31^ to combine decoding results across sessions. To correct for the comparisons over multiple regions we chose a level of 0.01 for the false discovery rate (FDR_0.01_).

Second, we computed *single-cell* statistics, testing whether the firing rates of single neurons correlated with three task variables (visual stimulus side, choice side, and feedback) in the appropriate epochs of each trial (Fig. 4c; Fig. S4c). Since the task variables of interest are themselves correlated, we used a condition-combined Mann-Whitney U-test^15^ which compares spike counts between trials differing in just one discrete task variable, with all others held constant. Using a permutation test, we determined the fraction of individual neurons in a region that are significantly selective to a variable, using a threshold specific to each variable. For each session with recordings in a specific region, we computed the significance score of the proportion of significant neurons by using the binomial distribution to estimate false positive events. We then combined significance scores across sessions with Fisher’s combined probability test^30, 31^ to obtain a combined p-value for each region. We report a region as being responsive to a variable if this combined p-value was below the chance level, correcting for multiple comparisons using FDR_0.01_. This method has lower statistical power than decoding since it only examines noisy single neurons, and it may therefore miss areas that have weak but distributed correlates of a variable. Moreover, it controls for correlated variables in a way that the decoding method does not, so it is selectively able to find neurons that do not only represent a variable by virtue of that variable’s correlation with a confound.

Third, we performed a *population trajectory* analysis (Fig. 4d; Fig. S4d), averaging firing rates of single neurons in a session across all trials in 20 ms bins and then aggregating all neurons across sessions and mice per brain region. We examined how trajectories in the high dimensional neural spaces reflected task variables. We did this by measuring the time-varying Euclidean distance between trajectories *d*(*t*) in the interval of interest, normalised by the square root of the number of recorded cells in the given region to obtain a distance in units of spikes/sec. From this time-varying distance, we extracted response amplitude and latency statistics. For significance testing we permuted trials for the variable of interest while keeping the other variables fixed to minimise the effect of correlations among the variables (as in the single-cell statistics), using FDR_0.01_ to control for multiple comparisons. Just for visualisation, we projected the trajectories into a 3-dimensional principal components space. This analysis combines all recordings into one “super-session” before computing a variable’s effect for each brain region, weighting each cell equally, thus combining recordings at the cell level for each region. This approach promises a strong signal-to-noise ratio, but cannot distinguish results obtained in individual sessions. It further stands out by providing the temporal evolution of a region’s sensitivity to a variable during the interval of interest.

Finally, we used an *encoding model*^32^ that fits the activity of each cell on each trial as a linear combination of a set of temporal kernels locked to each task event (Fig. 4e; Fig. S4e). This generalized linear model (GLM) quantifies the dependency at a temporally fine scale, at the cost of a potentially low signal-to-noise ratio. We measure the impact of a variable by removing its temporal kernels and quantifying the reduction in the fit of the activity of a neuron (typically assessed by Δ*R*^2^). This method lacks a convenient null distribution, and so we report effect sizes rather than significance.

It is important to note that we do not expect the different analysis methods to agree perfectly, since they focus on different aspects of the responsivity of individual neurons and populations thereof, and even, in the case of the population trajectory analysis, combining information across multiple sessions, rather than within single sessions (which also allowed us to use this method to test the robustness of our results by comparing findings based on subsets of the data; Fig. S5). The methods thus should be interpreted collectively. For a direct comparison of analysis scores, see flatmaps in Fig. S6 and scatter plots of scores for analysis pairs in Fig. S7.

Besides the four main analyses, we additionally performed a basic, inter-area analysis via Granger interaction for simultaneously recorded brain areas (Fig. S8). We find high Granger interaction scores for region pairs from all major brain regions, weakly correlated with anatomical distance, and mostly bidirectional. A similar analysis was performed with relation to the block variable in the companion paper^23^.

We next describe the results of our four analyses applied to the coding of each of the four task variables.

### Representation of Visual Stimulus

We first consider neural activity related to the visual stimulus. Classical brain regions in which visual responses are expected include superior colliculus^33, 34^, visual thalamus^35–37^, and visual cortical areas^38–41^, with latencies reflecting successive stages in the visual pathway^42, 43^. Correlates of visual stimuli have also been observed in other regions implicated in visual performance, such as parietal^44^ and frontal^45–48^ cortical areas as well as the striatum^49, 50^. Substantial encoding of visual stimuli may also be present beyond these classic pathways, since the retina sends outputs to a large number of brain regions^51^. Indeed, an initial survey^15^ of regions involved in a similar task uncovered visual responses in areas such as the midbrain reticular nucleus (MRN). Thus we hypothesised that visual coding would extend to diverse regions beyond those classically described.

Consistent with this hypothesis, a decoding analysis based on the first 100 ms following stimulus onset revealed correlates of the visual stimulus side in many cortical and subcortical regions, with strong signatures in visual cortex (VISam, VISI, VISa, VISp), prefrontal cortex (MOs), thalamus (LGd, LGv, CL), midbrain (NOT, SNr, MRN, SCm, APN), and hindbrain (GRN, PGRN; Fig. 5a,f). For instance, the activity of neurons in primary visual cortex (VISp) could be used to predict the stimulus side (Fig. 5i). Note that, uniquely amongst our analyses, the decoding analysis does not control for variables correlated with the visual stimulus, such as choice and block, so some of the regions with statistically significant results on this analysis might instead encode these variables.

**Figure 5.**
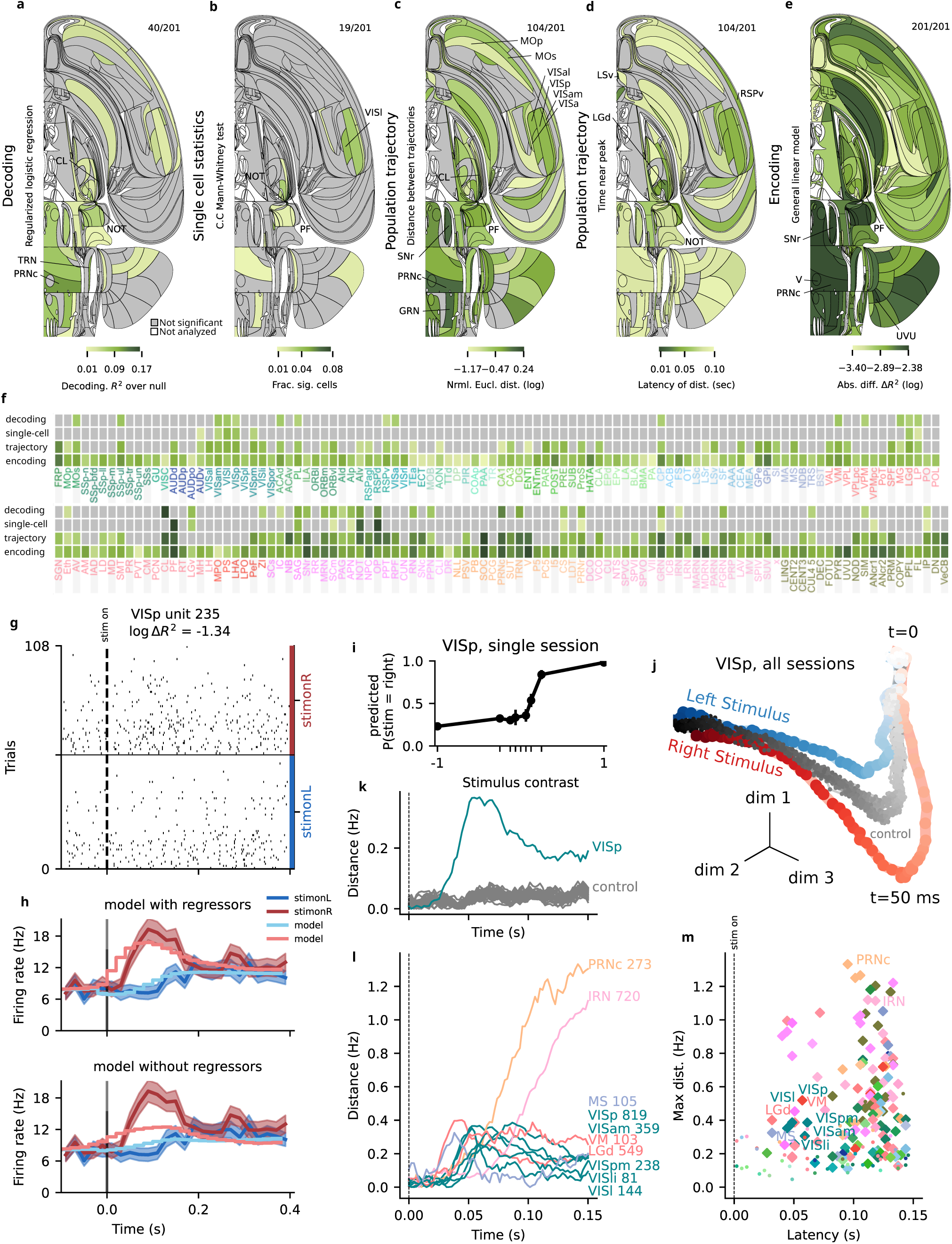
Representation of the Visual Stimulus. See also Fig. S9; S11 S12. An interactive version of this figure can be found on our data website. Analyses based on a period [0, 100] ms aligned to the stimulus onset. **a)** Decoding analysis: Flat brain map of decoding score (balanced accuracy) across sessions. The values have been corrected by subtracting the median of the decoding score in the null distribution. Colour represents effect size. Grey: regions in which decoding was not statistically significant, using FDR_0.01_ to correct for multiple comparisons; upper right ratio indicates significant regions out of all analysed. White: regions that were not analysed due to insufficient data. **b)** Single cell statistics: Flat brain map of the fraction of neurons in a region whose firing rates were significantly modulated by stimulus side (left vs right). The significance score of single cells were obtained by conjoining two tests: a Mann-Whitney test and a condition combined Mann-Whitney test (Methods), using thresholds of *p <* 0.001 and *p <* 0.05, respectively. The significance score of each region was computed by assuming a binomial distribution of false positive events, and FDR_0.01_ was used to correct for multiple comparisons. **c)** Population trajectory distance: Flat brain map of the time-resolved maximum distance between trajectories following stimuli on the left vs. right side, based on Euclidean distance (in spikes/second) in the full-dimensional space (dimension = number of cells) for each brain region. Significance was assessed relative to a shuffle control, using FDR_0.01_ to correct for multiple comparisons. **d)** Population trajectory latency: The earliest time after stimulus onset at which 70% of the maximum of the trajectory distance was reached, for significant regions only. **e)** Encoding analysis: Flat brain map of mean model improvement differences per region, across all neurons in that region computed as the absolute difference between the improvements (*|*Δ*R*^2^*|*) from the right stimulus kernel and left stimulus kernel regressors (400 ms causal kernels aligned to the stimulus onset time). A separation of this plot according to a median split of the first wheel movement time is shown in Fig. S15. **f)** Table of effect significance (grey - not significant; a-c) and effect size (by darkness; a-c; e), grouped by region. Regions are sorted canonically. **g)** Spike raster of example neuron in VISp identified by the encoding model as being sensitive to stimulus side (see Table 3 for session and neuron details). Trials per condition are in temporal order, only every third trial shown. **h)** Upper panel: Comparison of peri-event time histogram (PETH) of spiking activity for left and right stimuli for the example neuron in panel g conditioned on stimulus onset, along with associated predictions of the full encoding model. The width of the PETH traces reflects standard error of the mean. Lower panel: The same PETHs, but with predictions produced by an encoding model in which the stimulus onset-aligned regressors were omitted. Error bars represent 1 SEM about the mean rate at each time point. **i)** Predicted probability from decoding analysis of stimulus side (‘neurometric curve’) as a function of contrast from 40 neurons in VISp in a sample session. Note that the left-side stimulus contrasts are shown with *<* 0 values and the right-side stimulus contrasts are shown with *>* 0 values. Balanced accuracy for this region-session was 0.80; see Table 3 for session and neuron details. **j)** Trajectories obtained from left (right) stimulus side trial-averaged activity of all VISp neurons, visualised by projection via PCA onto 3 dimensions. Each dot corresponds to a single time bin of the trial-averaged population activity, darker colours indicate later times. Grey pseudo-trajectories were obtained by averaging randomly selected trials matched for block and choice but with different visual stimuli presented. (A clear split of pseudo-trajectories indicates correlation of block, choice and stimulus side, as stimulus sides were shuffled within block/choice classes, see Methods.) **k)** Trajectory distance as a function of time for example region VISp, showing an early response; pseudo-trajectory distances in grey (control). **l)** Trajectory distances for more example regions showing early response in most visual areas and ramping stimulus side modulation with time in others. The number of neurons is given alongside the acronym. **m)** Maximal population trajectory distance and modulation latency for all analysed regions (diamonds - significant regions, dots - not significant regions). The same information, but using a longer time window and the activity of all the neurons in the dataset (using only the constraint of 20 neurons in total per region) is shown in Fig. S16a;d;g.

Decoding performance varied across sessions, and therefore, in Fig. S9, we show performance across sessions for all regions, even those that are not significant after the FDR_0.01_ correction for multiple comparisons (also see Fig. S10 for a subset of these results split by sex of the animals; these did not reveal differences between male and female). Given that some regions may well represent visual information in localised sites that were only occasionally covered by our probes, we also report the fraction of sessions in which we were able to decode the stimulus significantly from a region to assess spatial distribution (Fig. S11a).

To distinguish the possible contributions of variables correlated with the visual stimulus, we next analysed responses in the same 100 ms window using a single-cell analysis, which controlled for other variables by holding them constant in each comparison of stimulus side. This analysis yielded a consistent picture but found fewer significant areas, with 0.5% of all neurons correlated with stimulus side (Fig. 5b). The significant regions included visual cortical areas (VISp, VISpm, VISam, VISl) and visual thalamus (LGd, LGv, LP), but also other structures such as auditory cortex (AUDv), dorsal thalamus (PF), parts of the midbrain (SCm, APN, NOT) and hindbrain (CS, PRNr, GRN, IP, ANcr1). However, even in those regions containing the largest fractions of responsive neurons (such as visual cortex), this fraction did not exceed ∼ 10%. Note that this low percentage of neurons could be the result of neurons not having receptive fields overlapping the stimulus position, given our grid-based approach to probe insertions.

To provide an overview of the variability across sessions, Fig. S12 shows the fraction of significant neurons broken down by sessions, without applying the FDR_0.01_ correction.

The results of the population trajectory analysis were consistent with the decoding analysis (Fig. 5c), and provided further information about the time course with which visual signals were encoded (Fig. 5d). For instance, the responses in visual area VISp to right vs. left visual stimuli show early divergence shortly after stimulus onset, followed by rapid convergence (Fig. 5j,k). The shuffled control trajectories (shown in grey) are close to the true trajectories (Fig. 5j) because this analysis controls for choice, which is tightly correlated with the stimulus. Altogether, this analysis indicated that distance was significant in 104 regions (Fig. 5c,f).

The evolution of trajectories over time could be distilled into two numbers (Fig. 5l,m): the maximal response (maximum distance or *d_max_*) and the response latency (first time to reach 70% of *d_max_*; mapped across the brain in Fig. 5d). This characterization of the dynamics of visual representations revealed that some areas had short latencies and early peaks, including classical areas (LGd, LP, VISp, VISam, VIpm), with a spatiotemporally finely resolved view of latency differences such that LGd *<* VISp ≈ LP *<* VISpm *<* VISam (latencies ≈ 34, 42, 42, 57, 78 ms; Fig. 5d,l,m). This early wave of activity was followed substantially later by significant visual encoding in other areas, including MRN, SCm, PRNr, IRN, and GRN (latencies ≈ 100 − 120 ms; Fig. 5d,l,m).

The encoding analysis characterised visual encoding in individual neurons across the brain (Fig. 5e). This analysis asks whether a prediction of single-trial activity can be improved by adding a temporally structured kernel that unfolds over 400 ms after stimulus onset, on top of activity related to feedback, wheel movement speed, and block identity. Since there is no convenient null distribution which could be used to test the significance of this improvement, we only report effect sizes for the encoding analysis. For instance, as expected, an example VISp neuron showed large differences between stimuli on the left and right (Fig. 5g) such that removing the visual kernel resulted in a poor fit of the neuron’s firing rate (Fig. 5h). This analysis indicates that the visual stimulus variable improved fits of encoding models for neurons across a wide range of brain regions (Fig. 5e,f).

These results were broadly consistent with receptive field (RF) measurements. At the end of the decision-making task, we performed RF mapping in most of the recording sessions (504/699 sessions). We thereby computed the visual RFs of neurons across the brain (total number of brain regions covered: 204 regions), including classical visual areas and beyond. We estimated the significance of the receptive field of each neuron by fitting the receptive field to a 2d-Gaussian function, and comparing the variance explained to the fitting of 200 random shuffles of each RF. Overall, we found a relatively small fraction of cells with significant receptive fields (Fig. S13). Those regions with large fractions tend to be classical visual regions (VISp, VISl, SCs). We also observed non-zero fractions in diverse areas beyond classical visual regions, including auditory cortex (AUDv), auditory thalamus (MG), parts of midbrain (MRN, SCm, APN, NOT), and hindbrain (ANcr1) (Fig. S13, S14), which further supports the results of neural analysis on coding of visual stimuli during the task.

We also separated out the effect sizes according to a median split of the first wheel movement time (Fig. S15. Regions with high explained variance from stimulus onset on all trials mostly appear as well in the early-first wheel movement time model (e.g. RSPv, VISl, PAG, and RN) while a handful of new cortical regions in motor areas (namely MOs, ORBm and ILA to a lesser extent) seem to be explained only when fitting early trials. Late response trials show fewer regions with well-explained (greater than an 0.03 change) variance, but there is also explained variance in subiculum and post subiculum which do not appear in the set of all trials.

Taken together, the decoding, single-cell statistics, and population trajectory analyses reveal a largely consistent picture of visual responsiveness that includes large and short-latency responses in classical areas but also extends to diverse other regions, even when controlling for correlated variables, particularly at later times relative to stimulus onset.

### Representation of Choice.

Next, we examined which regions of the brain represented the mouse’s choice, and in which order. Choice-related activity has been observed in parietal, frontal and premotor regions of the primate cortex^6, 52, 53^ where many neurons show ramping activity consistent with evidence accumulation^6, 54^. These choice signals develop across frontoparietal regions and appear later in frontal eye fields^55^. Similar responses were found in rodent parietal^56^, frontal^57, 58^, and premotor^59, 60^ cortical regions. However, in both rodents and primates, choice-related activity has also been found in hippocampal formation^61^ and subcortical areas, in particular in striatum^15, 62^, superior colliculus^15, 63, 64^ and other midbrain structures^15^. Subcortical regions show choice signals with similar timing as cortex^15, 16^ and play a causal role in the choice^64^. This evidence indicates that decision formation engages a distributed network of cortical and subcortical brain regions. Our recordings allowed us to determine in detail where and when choice-related activity emerges across the brain.

The decoding analysis suggested a representation of choice (left versus right upcoming action) in a larger number of brain regions than the representation of the visual stimulus (Fig. 6a,f). The animal’s choice could be decoded from neural population activity during a 100 ms time window prior to first wheel movement time in many analysed regions, with the strongest effect sizes in hindbrain (GRN, VII, PRNr, MARN), thalamus (CL), midbrain (SNr, RPF, MRN), hypothalamus (LPO), and cerebellum (CENT2). For instance, the activity of neurons in the gigantocellular reticular nucleus (GRN) of the medulla could be readily decoded to predict choice in an example session (Fig. 6g,i). Choice was also significantly decodable from somatosensory (SSp-ul), prelimbic (PL), motor (MOp, MOs), orbital (ORBvl), and visual (VISp) cortical areas.

**Figure 6.**
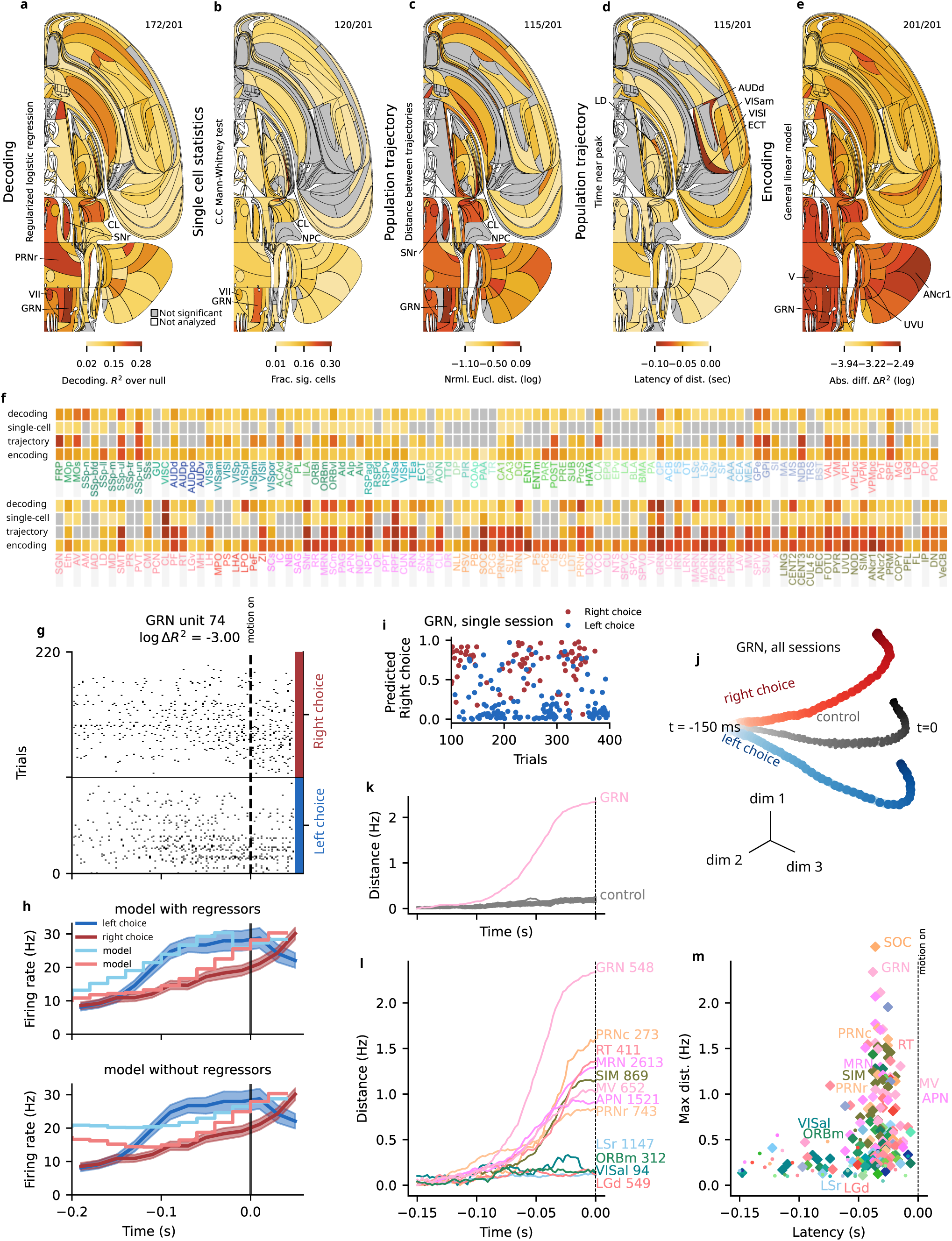
Representation of Choice. See also Fig. S9, S12, S17. An interactive version of this figure can be found on our data website. **a)** Decoding analysis: Flat brain map of median corrected decoding score (balanced accuracy) across sessions. The values were corrected by subtracting the median of the decoding score in the null distribution. Colour represents effect size. Grey: regions in which decoding was found not to be statistically significant, using FDR_0.01_ to correct for multiple comparisons. White: regions that were not analysed due to insufficient data. **b)** Single-cell statistics: Flat brain map of the fraction of neurons in a region whose firing rates were significantly modulated by choice side (left vs. right) in the period [*−*100, 0] ms aligned to the first wheel movement time. The significance scores of the single cells were obtained by conjoining two tests: a Mann-Whitney test and a condition combined Mann-Whitney test (Methods), using thresholds of *p <* 0.001 *and p <* 0.05, respectively. The significance score of each region was computed by assuming a binomial distribution of false positive events, and FDR_0.01_ was used to correct for multiple comparisons. **c)** Population trajectory distance: Flat brain map of the maximum distance between neural trajectories for left and right choice, based on time-resolved Euclidean distance (in spikes/second) in the full-dimensional space (the number of dimensions equals to the number of cells) for each brain region. Significance was assessed relative to a shuffle control, using FDR_0.01_ to correct for multiple comparisons. Canonical window lengths were employed, as shown in Fig. 4a. **d)** Population trajectory latency: The earliest times before first wheel movement time at which 70% of the maximum of the trajectory distance (see part c) was reached, for significant regions only. **e)** Encoding: Flat brain map of mean model improvement differences per region, across all neurons in that region computed as the absolute difference between the improvements (*|*Δ*R*^2^*|*) from the right first movement kernel and left first movement kernel regressors. These regressors for choices in each direction were 200 ms anti-causal kernels aligned to the first wheel movement time. A separation of this plot according to a median split of the first wheel movement time is shown in Fig. S15. **f)** Table of effect significance (grey - not significant; a-c) and effect size (by darkness; a-c; e), grouped by region. Regions are sorted canonically. **g)** Spike raster of an example neuron in GRN identified by the encoding model as sensitive to choice (see Table 3 for session and neuron details). Trials per condition are in temporal order, only every third trial shown. **h)** Upper panel: Comparison of PETHs aligned to first wheel movement time on left (blue) and right (red) choice trials for the example neuron in panel g, along with the encoding model predictions for each condition (dark colours). This neuron was selected for a high difference in 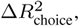 computed as the absolute difference of additional variance explained by the left/right first movement regressors. Lower panel: The same PETHs, but with predictions produced using a model lacking the left and right first movement regressors. Error bars represent 1 SEM about the mean rate at each time point. **i)** Decoding to predict choice for part of an example session from 68 neurons in GRN. Balanced accuracy for this region-session was 0.86; see Table 3 for session and neuron details. Decoder predictions separate in correlation with the mouse’ choices (blue - left choice, red - right choice). **j)** Trajectories obtained from trial-averaged activity of all GRN neurons on left (blue) and right (red) choice trials, visualised by projecting onto the first 3 PCA dimensions. Each dot corresponds to a time bin of the population activity, darker colours indicate times closer to the first wheel movement time. Grey pseudo-trajectories were obtained from trial-averaged activity on trials with randomised choice, controlling for correlations with stimulus and block. **k)** Trajectory distance as a function of time for GRN, showing ramping activity; pseudo-trajectory distances in grey (control). **l)** Trajectory distances for more example regions showing ramping choice-modulation with time. **m)** Maximal population trajectory distance and modulation latency for all analyzed regions (diamonds - significant regions, dots - not significant regions). The same information, but using but using a longer time window and the activity of all the neurons in the dataset (using only the constraint of 20 neurons in total per region) is shown in Fig. S16b;e;h.

Some of the decodable choice information, however, could be due to responses to the visual stimulus or block, which correlate with choice. We thus performed single-cell analyses that control for correlations between all these task variables. More single neurons responded significantly to choice than to the visual stimulus (Fig. 6b,f): firing rates of 4% of all neurons recorded across all brain regions correlated with choice direction during the 100 ms prior to the first wheel movement time, controlling for correlations with the stimulus and block. The largest fractions of neurons responding significantly to choice were in the hindbrain, cerebellar, midbrain, and thalamic regions, consistent with the results of decoding analysis. Neurons with significant responses to choice were highly prevalent in thalamus (CL, SPF), midbrain (SCm, MRN, SNr, RPF, NPC), pons, medulla, and cerebellar nuclei (GRN, IRN, SOC, VII, TRN, FOTU), most of which did not show visual responses. The prevalence of choice selective neurons in these subcortical regions was further confirmed by the single-cell encoding model (Fig. 6e). For instance, an example neuron in GRN showed stronger responses for right choices than left choices (Fig. 6g). The encoding model captured this preference but only in the presence of the kernel associated with choice (Fig. 6h), indicating choice selectivity.

The population trajectory analysis allowed us to compare across brain regions the magnitude of choice representation on the population level, measured as the distance between trajectories in the neural population state space on left versus right choice trials (Fig. 6c,f). The population-level choice representation was evident in many regions across the brain, with the strongest separation of neural trajectories in hind-brain (IRN, GRN, PRNc) and midbrain (APN, MRN, SCm), and similar magnitude of the population-level choice encoding in many other areas (Fig. 6c,f). In our example region GRN, the trajectories for left and right choice trials separated significantly more than in shuffled control (Fig. 6j,k, controlling for correlations with stimulus and block), and the magnitude of this separation was greatest across all brain regions (Fig. 6l). Thus, all our analyses consistently point to a distributed choice representation, with some of the strongest choice signals in midbrain, hindbrain and cerebellum, and relatively weaker choice encoding across many cortical areas.

Next we analysed when the choice signals emerged across the brain by measuring the latency with which neural population trajectories separated on the left versus right choice trials during the time preceding first wheel movement time (Methods). Some of the earliest choice signals developed nearly simultaneously in thalamus (LD), and cortex (VISl, VISam, ECT), and later appeared in a larger distributed set of brain areas (Fig. 6d,l,m). The regions GRN and MRN in the reticular formation showed moderate choice latencies and some of the strongest magnitude of population level choice representations (Fig. 6j,l,m), suggesting a role for these brainstem structures in decision formation or movement preparation.

For the encoding model, we also separated out the differences in variance explained according to a median split of the first wheel movement time (Fig. S15. In regions which show a high degree of variance explained by rightward movement onset RSPv again appears when fitting all trials and early trials, but not late trials. In the case of movement onset, however, secondary visual areas VISam and VISal are consistently involved in all trials along with motor areas. Notably the high variance explained in subiculum extends to hippocampal CA1 and post-subiculum only in late response trials, and does not appear at all in early response trials. Subcortical involvement seems limited to early trials in some regions like PAG which do not appear in the model fit to all trials.

### Representation of Feedback

At the end of each trial, the mouse received feedback for correct or incorrect responses: a liquid reward at the lick port or a noise burst stimulus with a time-out period. These positive or negative reinforcers influence the learning of the task^65–69^. The liquid is consumed through licking, an activity that likely involves prominent neural representations which, in this study, we will not be able to distinguish from the more abstract representation of reward. Feedback also activates neuromodulatory systems such as dopamine^70, 71^, which have widespread connections throughout cortical and subcortical regions. Nevertheless, it is unclear how widespread is the direct encoding of feedback signals in the brain.

The decoding analysis revealed nearly ubiquitous neural responses associated with reward delivery on correct versus incorrect trials, and likely the motor responses associated with its consumption (Fig. 7a,f). Using the neural responses in the 200 ms following feedback onset, we were able to decode whether the trial was correct from nearly all recorded brain regions (Fig. 7a,f). In many regions, decoding was practically perfect, as for instance the activity of the intermediate reticular nucleus in the hindbrain (IRN) in an example session (Fig. 7i).

**Figure 7.**
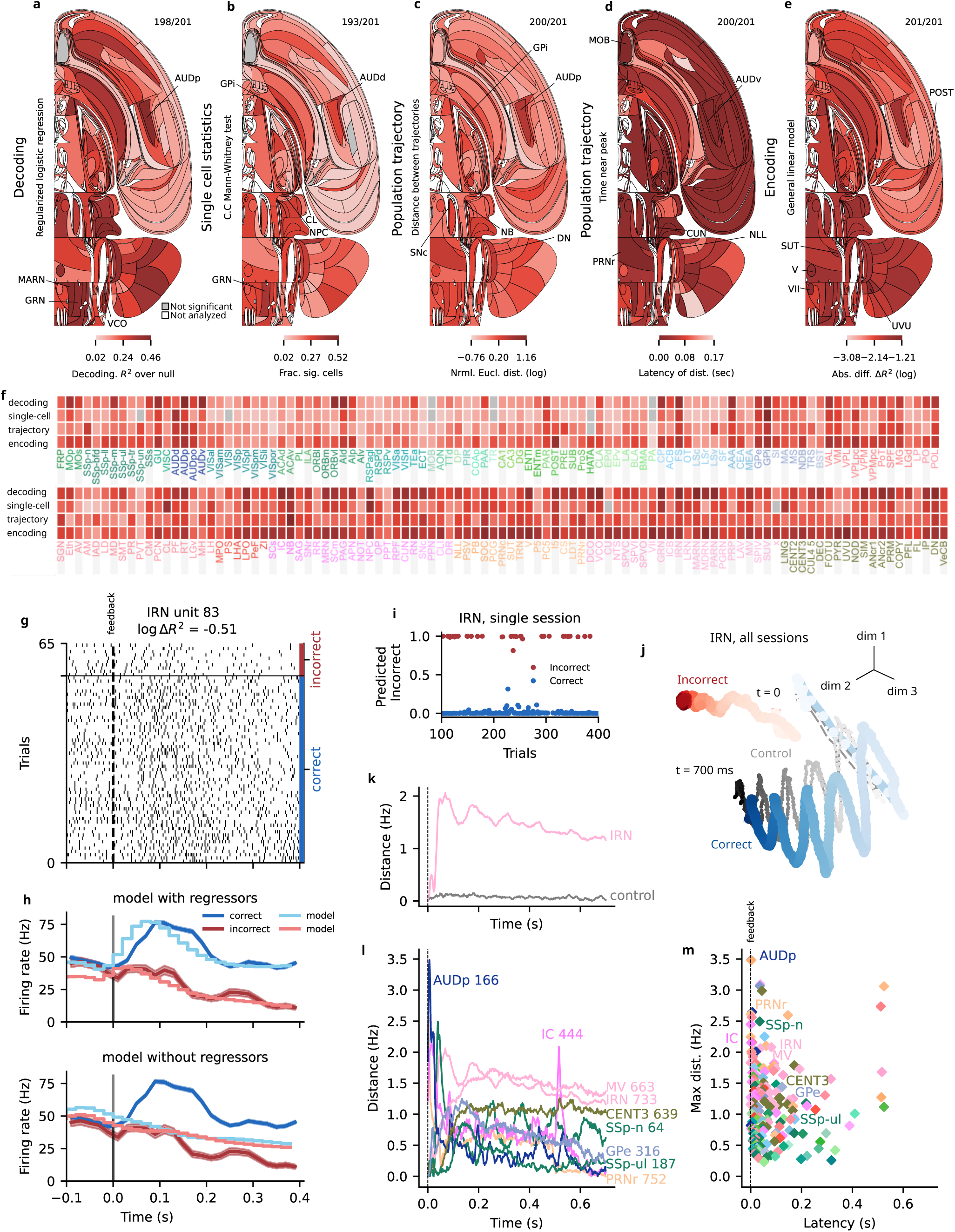
Representation of Feedback. See also Fig. S9, S12, S21. An interactive version of this figure can be found on our data website. **a)** Decoding: Flat brain map of median corrected decoding score (balanced accuracy) across sessions. The values have been corrected by subtracting the median of the decoding score in the null distribution. Colour represents effect size. Grey: regions in which decoding was found not to be statistically significant, using FDR_0.01_ to correct for multiple comparisons. White: regions that were not analysed due to insufficient data. **b)** Single-cell analysis: Flat brain map of the fraction of neurons whose firing rates were significantly modulated by feedback compared to baseline activity ([*−*200, 0] ms aligned to the stimulus onset) in the period following feedback onset ( [0, 200] ms). The significance scores of the single cells were obtained by conjoning two tests: a Mann-Whitney test and a condition combined Mann-Whitney test (Methods), using thresholds of *p <* 0.001 and *p <* 0.05, respectively. The significance score of each region was computed by assuming a binomial distribution of false positive events, and FDR_0.01_ was used to correct for multiple comparisons. **c)** Population trajectory distance: Flat brain map of the time-resolved maximum distance between correct and incorrect choice trajectories, based on Euclidean distance (in spikes/section) in the full-dimensional space (dimension = number of cells across all sessions) for each brain region. Significance was assessed relative to a shuffle control, using FDR_0.01_ to correct for multiple comparisons. Canonical window lengths were employed, as shown in Fig. 4a. **d)** Population trajectory latency: The earliest time after feedback at which 70% of the maximum trajectory distance (see part c) was reached, for significant regions only. **e)** Encoding: Flat brain map of the mean model differences per region, across all neurons in that region, computed as the log of the absolute difference between the improvements (Δ*R*^2^) from the correct feedback kernel and incorrect feedback kernel regressors. These regressors are 400 ms causal kernels aligned to the feedback time. **f)** Tabular form of effect significance (grey - not significant; a-c) and effect size (by darkness; a-c, e). Regions are sorted canonically. Spike raster of example neuron in IRN identified by encoding model as sensitive to feedback (see Table 3 for session and neuron details). Trials per condition are in temporal order, only every third trial shown. **h)** Upper panel: Comparison of PETHs aligned to first wheel movement time on correct (blue) and incorrect (red) trials for the example IRN neuron in panel g, along with the encoding model predictions for each condition (grey). This neuron was selected for a high difference in 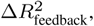 computed as the absolute difference of additional variance explained by the feedback regressors. Lower panel: The same PETHs but with predictions produced using a model lacking the feedback regressors. Error bars represent 1 SEM about the mean rate at each time point. **i)** Predicted probability from decoding analysis about whether reward was received, coloured by true feedback, from 39 neurons in IRN in a sample session. Balanced accuracy for this region-session was 1.00; see Table 3 for session and neuron details. **j)** Trajectories obtained from incorrect (correct) trial-averaged activity of all neurons in IRN vizualised by projection via PCA to 3 dimensions. Each dot corresponds to a time bin of the population activity, darker colours indicate later times. The oscillation of the blue trajectory correlated with licking, and was much stronger in correct trials when the animals received water. Grey pseudo-trajectories were obtained by averaging randomised trials, shuffling choice types within classes of stimulus side and block. **k)** Trajectory distance between correct and incorrect trials, as a function of time for example region IRN, showing oscillatory (licking) activity; pseudo-trajectory distances in grey (control). **l)** Trajectory distances for more example regions showing early response e.g. in auditory areas and prolonged feedback type modulation with time in others. Note IC = inferior colliculus, is known to relay auditory signals, hence first (second) peak at 0.5 sec after feedback, when the noise burst starts (ends) for incorrect trials. **m)** Maximal population trajectory distance and modulation latency for all analyzed regions (diamonds - significant regions, dots - not significant regions). Most regions’ activity was modulated within 100 ms after feedback. The same information, but using but using a longer time window and the activity of all the neurons in the dataset (using only the constraint of 20 neurons in total per region) is shown in Fig. S16c;f;i.

Our single-cell statistics applied to the same trial interval confirmed the decoding results. Neurons with significant response changes to correct versus incorrect feedback or reward consumption were extremely widespread (Fig. 7b,f), with only a small handful of regions not significant for feedback type. The same was true for feedback versus the inter-trial interval baseline (Fig. S18).

Population trajectory analysis also found statistically significant response differences for correct versus incorrect responses across every recorded brain region, predominantly consistent with the other analyses (Fig. 7c,f). It confirmed the relative strength of hindbrain, midbrain, and thalamic responses to feedback seen across analyses. Population trajectory analysis also revealed asymmetries in response to negative versus positive feedback: for positive feedback the response was overall stronger, and multiple brain areas exhibited a coherent ∼10 Hz oscillatory dynamics during reward delivery that was phase-locked across brain areas (Fig. 7j,k,l) and sessions (Fig. S19). Across-session coherence is visible as a large oscillatory signal in example area IRN (Fig. 7j,k). These oscillatory dynamics were missing during negative feedback, and were closely related to licking behaviour (Fig. S19)^72–75^, pointing to consumption-related activity being the dominant factor over more-abstract influences of reward on neural activity.

Assessing response latencies by the divergence of the trajectories over time, we found that the saccade- and gaze-reorienting brainstem region PRNr (rostral pontine reticular nucleus) and the primary auditory region AUDp exhibited the earliest and strongest responses (Fig. 7l,m). Some early responsivity is likely a carry-over from choice-related activity because the latencies are very short and several identified areas exhibit high choice responsivity. The responses from auditory areas likely reflect responses to the error tone and the click from the reward delivery valve. This is especially clear for region IC (inferior colliculus - a region known to relay auditory signals) which has a peak at the start and end of the 0.5 s long error noise burst, Fig. 7l. After these initial responses, latencies across other brain regions appeared roughly similar, suggesting a common signal broadcast across the brain (Fig. 7d). More detailed trajectory distance and latency scatterplots can be found in Fig. S16.

We applied the encoding model to the responses measured in the 400 ms after stimulus onset, and found that the kernel for correct feedback was the largest single contributor to neural response variance across each trial (Mean Δ*R*^2^ of 8.6 × 10*^−^*^3^ averaged across all neurons; Fig. S20), surpassing all other kernels (left or right stimulus, left or right wheel movement, incorrect feedback, block probability, and wheel speed). This high variance-explaining response to reward delivery or consumption held across both wide regions of cortex and subcortical areas. Mid- and hindbrain areas exhibited particularly strong responses to reward delivery, with many additional regions including thalamus and sensory (e.g. AUDp, SSs) and motor (MOp) cortex showing sensitivity as well (Fig. 7e). Removing the regression kernel for correct feedback then refitting the encoding model of an IRN neuron illustrates the large influence of correct feedback on activity (Fig. 7g,h).

In sum, we found feedback signals to be present across nearly all recorded brain regions, with a stronger response to positive feedback (i.e., to reward delivery and consumption) and with particularly strong responses in thalamus, midbrain, and hindbrain. Further research will be needed to distinguish between responses for an internal expectation of feedback or the initiation of choice-related action versus responses to external feedback.

### Representation of Wheel Movement

A consistent finding from previous large-scale recordings in mice has been the macroscopic impact of movement on neural activity, with both task-related and task-unrelated movements influencing activity well outside premotor, motor, and somatosensory cortical areas^18, 19, 76, 77^. Here, we start from the task-dependent component of movement, namely the movement of the wheel to register a response. We observed that different mice (and potentially the same mouse on different sessions) adopted different strategies for moving the wheel – for instance, some used both front paws; others only one. Turning the wheel is also a relatively complex operation, rather than being just a simple, ballistic movement. Thus, one should not expect a simple relationship between these movements and activity in laterally specific motor areas. For simplicity, we restrict our analyses to the activity associated with both wheel velocity (signed to distinguish left from right movements) and its absolute value, wheel speed. Furthermore, unlike the other task variables, movement trajectories change relatively quickly, necessitating different analysis and null control strategies. Accordingly, we only report simple decoding and encoding analyses.

Wheel speed was decodable from 81% of the reportable areas (163/201), with the strongest effect sizes in hypothalamus (LPO), hindbrain (MARN, GRN), midbrain (CLI), thalamus (CL, PF), cortex (ORBvl, VISC, VISrl), and cerebellum (VeCB; Fig. 8a,e). For example, we could readily decode wheel speed from single trials of activity in region GRN (Fig. 8g).

**Figure 8.**
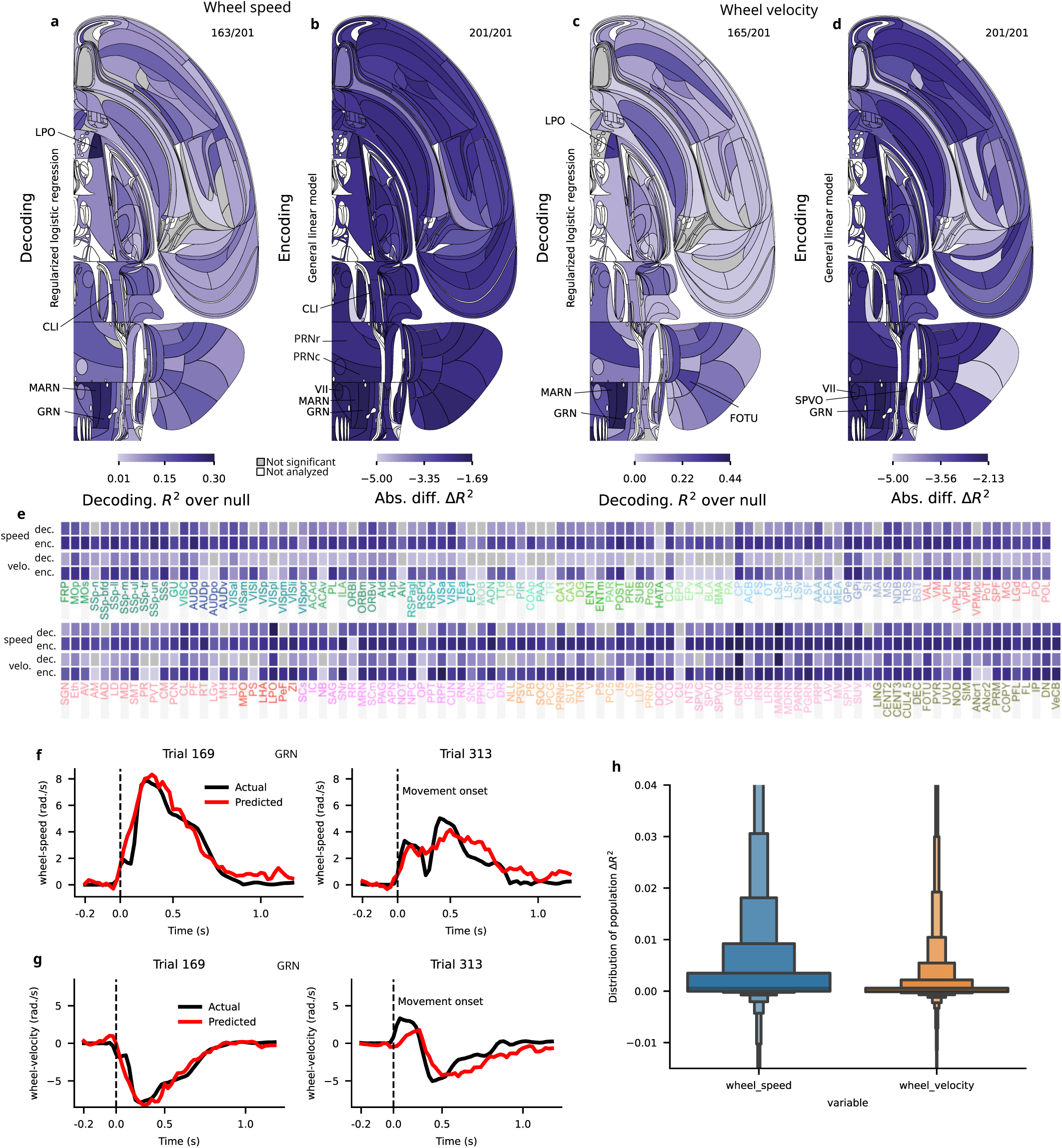
Representation of Wheel Movement. See also Fig. S9, S22. An interactive version of this figure can be found on our data website for speed and velocity. **a)** Decoding: Flat brain map of median corrected decoding score (*R*^2^) across sessions for wheel speed (the absolute value of the wheel velocity). The values have been corrected by subtracting the median of the decoding score in the null distribution. Colour represents effect size. Grey: regions in which decoding was found not to be statistically significant, using FDR_0.01_ to correct for multiple comparisons. White: regions that were not analysed due to insufficient data. **b)** Encoding analysis: Flat brain map of mean model improvement differences per-region, across all neurons in that region computed as the improvements (Δ*R*^2^) from the wheel speed regressor. These regressors are 200 ms anti-causal kernels convolved with the trace of wheel speed. **c,d)** Same as a,b but for wheel velocity rather than wheel speed. Note that the encoding results involve a completely separate model fit using the velocity rather than speed. **e)** Tabular form of effect significance for wheel speed (velocity) (grey - not significant; white - not analyzed) and effect size (by darkness; as in Swansons above), grouped by region. Regions are sorted canonically. **f)** Comparison of actual and predicted wheel speed for an example trial from 68 neurons in region GRN. *R*^2^ for this region-session was 0.73; see Table 3 for session and neuron details. **g)** Same as f, for a target signal of velocity rather than speed (*R*^2^ was 0.72). **h)** A comparison of the distributions of additional variance Δ*R*^2^ explained, across the whole population, when using either wheel speed or velocity as the base signal. Distributions truncated for clarity.

The encoding analysis confirmed that many regions across the brain were sensitive to the wheel speed during the task, with Δ*R*^2^ taking values up to several times larger than for the other variables considered besides feedback (Fig. 8b,e; S20). The pontine reticular nucleus (PRNc) and gigantocellular reticular nucleus (GRN), in particular, stood out in our analysis for the mean Δ*R*^2^ for neurons within those regions (mean Δ*R*^2^ = 9.4 × 10*^−^*^3^ in PRNc, and Δ*R*^2^ = 179 × 10*^−^*^3^ in GRN). Many other cortical (e.g. MOs) and subcortical (e.g. GPe, GPi, CP) regions had less substantial, but still above-average, correlations with the wheel speed relative to other regressors (Fig. 8b,e).

Wheel velocity was also significantly decodable from a rather similar collection of areas as wheel speed (Fig. 8c,e,g; S22a), and was also duly encoded (Fig. 8d,e) albeit with generally smaller values of Δ*R*^2^ (Fig. 8h). The apparently high decodability of velocity was unexpected given the complexities of wheel movement mentioned above, and indeed the uncorrected values of *R*^2^ for decoding speed were substantially larger than those for velocity in most regions (Fig. S22b). However, the null distribution based on imposter sessions (i.e., wheel movements from other sessions, including from other mice; see Methods) can be decoded much more accurately for speed than for velocity (Fig. S22c), reducing the statistical significance of the decoding of speed. We attributed this excess decodability of the null distribution to the more stereotyped, i.e., less variable, trajectory of speed (Fig. S22d).

We also correlated neural activity with behavioural movement traces extracted from video (nose, paw, pupil, and tongue). To test for significance we used the linear-shift method, comparing the correlation of spiking activity with behavioural movement variables against a null distribution in which the movement variables were shifted in time^78^ (Methods, Fig. S23a). More than half of the neurons in most brain regions were significantly correlated with at least one behavioural variable (Fig. S23b).

The widespread relationship between neural activity and motion has various potential sources. These include the specific details of motor planning and execution, efference copy^79^, somatosensory feedback, the suppression of input associated with self-motion^80^, and more subtle effects such as the change in other sensory inputs caused by the movement^18^, or even prediction errors associated with incompetent execution that can fine-tune future performance^81^. Others are more general, including arousal and the calculation and processing of the costs of movement (which would then be balanced against future gain)^82^. More generally, of the components that are indeed specific, only a fraction is likely to be associated with the wheel movement that we monitored, as opposed to other task-related motor actions. This is especially true given the results of previous studies^18, 19^ showing just how important uninstructed movements are in modulating a wide swathe of neural activity.

## Discussion

Building on previous efforts to build large-scale maps of activity in the mouse at neuronal resolution using Neuropixels probes^14–16^ and in other species using imaging (e.g., Refs.^83–85)^, we have developed a new strategy for assembling brain-wide electrophysiological maps by pooling data from numerous laboratories that employed the same standardised and reproducible task^22^. The result is an unprecedented brain-wide consensus map of neural activity of the mouse brain. This map reveals the neural activity that underlies performance in a rich task that requires processing of visual stimuli, rewards, and motor responses.

In addition to releasing the data – which are available to download via an API and to view interactively – we also performed a battery of standard analyses coupled with rigorous statistical methods, which suggest that neural activity throughout the entire brain correlates with some aspects of the task, but with major differences in the ubiquity of representation of different task variables (Figs. 5-8; see also Figs. S6, S9, S12 for side by side comparisons).

The neural representations of feedback (Fig. 7) and movement^15, 18, 19^ (Fig. 8) were particularly widespread. The former may also partly or primarily reflect the licking movements required for reward consumption, rather than the hedonic aspects of reward as such. Distinguishing these possibilities would require experiments recording activity during the presentation of reinforcement with no motor correlates, such as optogenetic stimulation of dopamine systems^86, 87^, or by comparing correlates of rewards with a motor correlate to the same movements when their hedonic reward was devalued, for example by satiation. The brain-wide correlates of movement could potentially reflect a brain-wide change in the state of neural processing during movement periods, along with specific encoding of motor features. The hypothesis of a brain-wide state change is consistent with findings that the neural representation of upcoming movements in cortex is extremely widespread, though not all of this activity is causally related to the performance of the movements^46^.

The subject’s upcoming choice was represented in the activity of neurons across brain systems including cortex, basal ganglia, thalamus, midbrain, hindbrain, and cerebellum (Fig. 6). These representations cannot reflect sensory reafference (i.e. responses related to sensory stimuli that occur as part of the movements, such as pressure on the paws and movements of the visual stimuli across the screen) since we only analysed the time period prior to the earliest detectable first wheel movement time. Moreover, our carefully controlled task design and pseudo-session statistical methods mean that choice coding reported in the single-cell and population trajectory analyses cannot reflect processing of the visual stimulus nor non-specific brain states such as arousal. Instead, these responses reflect aspects of decision formation or motor preparation, potentially including corollary discharge specific to the chosen action^88, 89^. This study is the first report at single-cell resolution of choice coding across these brain systems in a single task, and while many past studies have focused on the role of cortex, basal ganglia, or midbrain in visual perceptual decisions^6, 15, 54, 55, 62–64, 90, 91^, here we discovered that parts of the medulla, pons, and cerebellum are all selectively responsive with similar timing to those areas. Our data are not able to determine whether these different systems make unique contributions to decision formation and execution. However, they rule out a model in which only a limited set of systems subserve a given behaviour according to specific task demands.

The visual stimulus (Fig. 5) was represented (prior to movement) in a more restricted manner. Its processing followed a temporal sequence through traditional visual areas from the visual thalamus to cortex and to midbrain and hindbrain regions whose activity also correlated with choices. Importantly, the temporal structure of activity in these two groups of regions differed, with visual representations in classical visual regions showing a transient representation of the stimuli, and activity in midbrain and hindbrain showing later, ramping activity, consistent with a role of this activity in decision-making. The fact that visual information was found in hindbrain regions such as GRN and PRNr, even after accounting for correlates of choice, suggests a role for these regions in all phases of the cognitive decision-making process, rather than simply low-level motor control.

Although more than half of the recorded neurons in most brain regions were significantly modulated by at least some aspect of the task (Fig. S24), our ability to explain the total variance of single neurons was limited (Fig. S20). This indicates that the vast bulk of activity in the brain is not modulated by the task. It may instead be related to uninstructed movements^18, 19^ or other processes that are not timed to the task events. Even for the activity that is modulated by the task, it is notable that external cue-driven responses were consistently smaller than internally generated signals such as those arising in relation to integration of the stimulus and movement planning. As ever, absence of evidence for a neural representation of a task variable in a given region cannot be taken to indicate evidence of absence – particularly given that, for robustness, we used simple variants of analysis methods rather than, for instance, extensively parameterised deep neural networks.

Importantly, while our coverage is extensive, we do not have uniform coverage across all brain regions, particularly when considering the neurons that pass our stringent quality control metrics. This could bias our results against discovering task correlates in regions with specific anatomical arrangements (e.g., where cell bodies are densely packed making spike sorting more challenging). Furthermore, our behavioural training protocol is aimed at reducing individual differences in performance, which impedes a complete analysis of the relationship between neural activity in particular regions and factors such as the reward rate.

In summary, we have provided a strategy for examining brain-wide computations systematically, openly sharing a database of recordings spanning one entire mouse brain hemisphere as subjects performed a complex behavioural task. Our analyses have begun to elucidate the widespread representations of key task variables. Our freely available dataset provides a rich resource for deeper investigation of brain-wide neural computations, such as detailed analyses at the level of sub-regions (e.g. cortical layers or functional zones of the striatum) and cell types (as identifiable from extracellular waveforms, such as broad versus narrow spike shapes in cortex).

## Resources

### Data access

Please follow these instructions to download the data used in this article. You can also browse the data at https://viz.internationalbrainlab.org.

### Data quality

Please read our white paper on the released data for additional details about quality control and metrics.

### Code repository

Please visit our Github repository to access the code used to produce the results and figures presented in this article.

### Protocols and pipelines

Please visit our two Figshare repositories to access the protocols and pipelines used to train mice and perform the electrophysiology recording and histology validation.

## Methods

All procedures and experiments were carried out in accordance with the local laws and following approval by the relevant institutions: the Animal Welfare Ethical Review Body of University College London; the Institutional Animal Care and Use Committees of Cold Spring Harbor Laboratory, Princeton University, University of Washington, University of California at Berkeley and University of California at Los Angeles; the University Animal Welfare Committee of New York University; and the Portuguese Veterinary General Board.

### Animals

Mice were housed under a 12/12 h light/dark cycle (normal or inverted depending on the laboratory) with food and water available ad libitum, except during behavioural training days. Electrophysiological recordings and behavioural training were performed during either the dark or light phase of the cycle depending on the laboratory. The data from N=139 adult mice (C57BL/6; 94 male and 45 female, obtained from either Jackson Laboratory or Charles River) are used in this study. Mice were aged 13-178 weeks (mean 44.96 weeks, median 27.0 weeks) and weighed 16.1-35.7 g (mean 23.9 g, median 23.84 g) on the day of electrophysiological recording.

### Headbar implant surgery

A detailed account of the surgical methods for the headbar implant is in Appendix 1 of Ref.^22^. Briefly, mice were anesthetised with isoflurane and head-fixed in a stereotaxic frame. The hair was then removed from their scalp, which was subsequently removed along with the underlying periosteum. was Once the skull was exposed, Bregma and Lambda were marked. The head was positioned along the anterior-posterior and left-right axes using stereotaxic coordinates. The head bar was then placed in one of three stereotactically defined locations and cemented (Super-Bond C&B) in place. Future craniotomy positions were marked on the skull relative to Bregma. The exposed skull was then covered with cement and clear UV curing glue (Norland Optical Adhesives).

### Materials and apparatus

For detailed parts lists and installation instructions for the training rigs, see Appendix 3 of Ref.^22^; for the electrophysiology rigs, see Appendix 1 of Ref.^24^.

Each lab installed a standardised electrophysiological rig, which differed slightly from the apparatus used during behavioural training ^22^. The structure of the rig was constructed from Thorlabs parts and was placed on an air table (Newport, M-VIS3036-SG2-325A) surrounded by a custom acoustical cabinet. A static head bar fixation clamp and a 3D-printed mouse holder were used to hold a mouse such that its forepaws rested on the steering wheel (86652 and 32019, LEGO)^22^. Silicone tubing controlled by a pinch valve (225P011-21, NResearch) was used to deliver water rewards to the mouse. Visual stimuli were displayed on an LCD screen (LP097Q × 1, LG). To measure the timing of changes in the visual stimulus, a patch of pixels on the LCD screen flipped between white and black at every stimulus change, and this flip was captured with a photodiode (Bpod Frame2TTL, Sanworks). Ambient temperature, humidity, and barometric air pressure were measured with the Bpod Ambient module (Sanworks) and wheel position was monitored with a rotary encoder (05.2400.1122.1024, Kubler). Videos of the mouse were recorded from 3 angles (left, right and body) with USB cameras (CM3-U3-13Y3M-CS, Point Grey) sampling at 60, 150, 30 Hz respectively (for details see Appendix 1 of Ref.^24^). A custom speaker (Hardware Team of the Champalimaud Foundation for the Unknown, V1.1) was used to play task-related sounds, and an ultrasonic microphone (Ultramic UM200K, Dodotronic) was used to record ambient noise from the rig. All task-related data was coordinated by a Bpod State Machine (Sanworks). The task logic was programmed in Python and the visual stimulus presentation and video capture was handled by Bonsai^92^ and the BonVision package^93^.

Neural recordings were made using Neuropixels probes, either version 1.0 (3A or 3B2, N=109 and N=586 insertions) or 2.4 (N=4 insertions) (Imec^12^), advanced in the brain using a micromanipulator (Sen-sapex, uMp-4). Typically, the probes were tilted at a 15 degree angle from the vertical line. Data were acquired via an FPGA (for 3A probes) or PXI (for 3B and 1.0 probes, National Instruments) system using SpikeGLX, and stored on a PC.

### Habituation, training, and experimental protocol

For a detailed protocol on animal training, see Methods in Refs.^22, 24^. Briefly, at the beginning of each trial, the mouse was required to not move the wheel for a quiescence period of 400–700 ms. After the quiescence period, a visual stimulus (Gabor patch) appeared on either the left or right (±35*^◦^* azimuth) of the screen, with a contrast randomly sampled from a predefined set (100, 25, 12.5, 6, 0%). A 100 ms tone (5 kHz sine wave) was played at stimulus onset. Mice had 60 s to move the wheel and make a response. Stimuli were yoked to the rotation of the response wheel, such that a 1 millimetre movement of the wheel moved the stimulus by 4 visual degrees. A response was registered if the centre of the stimulus crossed the ±35*^◦^* azimuth line from its original position. If the mouse correctly moved the stimulus 35*^◦^* to the centre of the screen, it immediately received a 3 *µ*L reward; if it incorrectly moved the stimulus 35*^◦^* away from the centre, it received a timeout. If the mouse responded incorrectly or failed to reach either threshold within the 60 s window, a white noise burst was played for 500 ms and the inter-trial interval was set to 2 s. In trials where the visual stimulus contrast was set to 0%, the mouse had to respond as for any other trial by turning the wheel in the correct direction (assigned according to the statistics of the prevailing block) to receive a reward, but the mouse was not able to perceive whether the stimulus was presented on the left or right side of the screen. The mouse also received feedback (noise burst or reward) on 0% contrast trials.

Each session started with 90 trials in which the probability of a visual stimulus appearing on the left or right side was equal. Specifically, the 100%, 25%, 12.5%, and 6% contrast trials were each presented 10 times on each side, and the 0% contrast was presented 10 times in total (i.e. the ratio of the [100 : 25 : 12.5 : 6 : 0] % contrasts is set at [2 : 2 : 2 : 2 : 1]). The side (and thus correct movement) for the 0% contrast trials was chosen randomly between the right and left with equal probability. This initial block of 90 trials is referred to as the unbiased block (50/50).

After the unbiased block, trials were presented in biased blocks: in right bias blocks, stimuli appeared on the right on 80% of the trials, while in left bias blocks, stimuli appeared on the right on 20% of the trials. The ratio of the contrasts remained as above ([2 : 2 : 2 : 2 : 1]). Whether the first biased block in a session was left or right was chosen randomly, and blocks then alternated. The length of a block was drawn from a exponential distribution with scale parameter 60 trials, but truncated to lie between 20 and 100 trials.

The automated shaping protocol for training ^22^ involved two collections of sessions. In the first, the animals started performing a version of the task without biased blocks, being progressively introduced to harder stimuli with weaker contrasts as they became progressively more competent. They also experienced a debiasing protocol intended to dissuade them from persisting with just one of the choices. Once they were performing sufficiently well on all non-zero contrasts, they were faced with the biased blocks. When, in turn, performance on those was adequate (including on 0% contrast trials which are informed by the block), they graduated to recording. Fig. S25A shows a joint histogram of the number of sessions the mice took in the first and second collections – these were not correlated. Fig. S25B shows a joint histogram of the number of sessions the mice took in the second collection and the performance during the recording sessions. These were also not correlated.

### Electrophysiological recording using Neuropixels probes

For details on the craniotomy surgery, see Appendix 3 of Ref.^24^.

Briefly, upon the first day of electrophysiological recording, the animal was anaesthetised using isoflurane and surgically prepared. The mouse was administered with analgesics (typically Carprofen) subcutaneously. The glue was removed (typically using a biopsy punch (Kai Disposable Biopsy Punches (1mm)) or a drill), exposing the skull over the planned craniotomy site(s). A test was made to check whether the implant could hold liquid; the bath was then grounded either via a lose or implanted pin. One or two craniotomies (approximately 1 × 1 mm) were made over the marked locations. The dura was left intact, and the brain was lubricated with ACSF. A moisturising sealant was applied over the dura (typically DuraGel (Cambridge NeuroTech) covered with a layer of Kwikcast (World precision instruments). The mouse was left to recover in a heating chamber until locomotor and grooming activity were fully recovered.

Subjects were head-fixed for recording after a minimum recovery period of 2 hours. Once a craniotomy was made, up to 4 subsequent recording sessions were made in that same craniotomy. Once the first set of craniotomy was fully recorded from, a subject could undergo another craniotomy surgery in accordance with the institutional licence. Up to two probes were implanted in the brain on a given session. CM-Dil (V22888 Thermofisher) was used to label probes for subsequent histology.

### Serial section two-photon imaging

Mice were given a terminal dose of pentobarbital and perfuse-fixed with PBS followed by 4% formaldehyde solution (Thermofisher 28908) in 0.1M PB pH 7.4. The whole mouse brain was dissected, and post-fixed in the same fixative for a minimum of 24 hours at room temperature. Tissues were washed and stored for up to 2-3 weeks in PBS at 4C, prior to shipment to the Sainsbury Wellcome Centre for image acquisition. For full details, see Appendix 5 of Ref.^24^.

For imaging, brains were equilibrated with 50mM PB solution and embedded into 5% agarose gel blocks. The brains were imaged using serial section two-photon microscopy^94, 95^. The microscope was controlled with ScanImage Basic (Vidrio Technologies, USA), and BakingTray, a custom software wrapper for setting up the imaging parameters^96^. Image tiles were assembled into 2D planes using StitchIt^97^. Whole brain coronal image stacks were acquired at a resolution of 4.4 x 4.4 x 25.0 *µ*m in XYZ, with a two-photon laser wavelength of 920 nm, and approximately 150 mW at the sample. The microscope cut 50 µm sections but imaged two optical planes within each slice at depths of about 30 µm and 55 µm from the tissue surface. Two channels of image data were acquired simultaneously using multialkali PMTs (‘Green’ at 525 nm ±25 nm; ‘Red’ at 570 nm low pass).

Whole brain images were downsampled to 25 *µ*m isotropic voxels and registered to the adult mouse Allen common coordinate framework^5^ using BrainRegister^98^, an elastix-based^99^ registration pipeline with optimised parameters for mouse brain registration. For full details, see Appendix 7 of Ref.^24^.

### Probe track tracing and alignment

Neuropixels probe tracks were manually traced to yield a probe trajectory using Lasagna^100^, a Python-based image viewer equipped with a plugin tailored for this task. Traced probe track data was uploaded to an Alyx server^101^; a database designed for experimental neuroscience laboratories. Neuropixels channels were then manually aligned to anatomical features along the trajectory using electrophysiological landmarks with a custom electrophysiology alignment tool^102, 103^. For full details, see Appendix 6 of Ref.^24^.

### Spike sorting

The spike sorting pipeline used at IBL is described in detail in Ref.^104^. Briefly, spike sorting was performed using a modified version of the Kilosort 2.5 algorithm^105^. We found it necessary to improve the original code in several aspects (scalability, reproducibility, and stability, as discussed in Ref.^24^), and developed an open-source Python port; the code repository is in Ref.^106^.

### Inclusion criteria

We applied a set of inclusion criteria to sessions, probes and neurons to ensure data quality. Table 1 lists the consequences of these criteria for the number of sessions and probes that survived. They are described below.

### Sessions and insertions

Each Neuropixels insertion was repeated in at least two laboratories, with reproducibility of outcomes across laboratories verified with extensive analyses that we have previously reported^24^.

Sessions were included in the data release if the mice performed at least 250 trials, with a performance of at least 90% correct on 100% contrast trials for both left and right blocks, and, in order to be able to analyze the feedback variable, if there were at least three trials with incorrect choices (after applying the trial exclusions below). Furthermore, sessions were included in the release only if they reached threshold on a collection of hardware tests (definitions can be found at https://int-brain-lab.github.io/iblenv/_autosummary/ibllib.qc.task_metrics.html)

Insertions were excluded if the neural data failed the whole recording visually assessed criteria of the ‘Recording Inclusion metrics and Guidelines for Optimal Reproducibility’ (RIGOR) from Ref.^24^, by presenting major artefacts (see examples in Ref.^104^), or if the probe tract could not be recovered during the histology procedure. Furthermore, only insertions whose alignments had been resolved (see Appendix 6 of Ref.^24^ for definition) were used in this study.

After applying these criteria, a total of 459 sessions, 699 insertions and 621733 neurons remained, constituting the publicly released data set.

### Trials

For the analyses presented here, trials were excluded if one of the following trial events could not be detected: `choice`, `probabilityLeft`, `feedbackType`, `feedback times`, `stimOn times`, `firstMovement times`. Trials were further excluded if the time between stimulus onset and first movement of the wheel (the first wheel movement time) was outside the range of [0.08, 2.00] seconds.

### Neurons and brain regions

Neurons yielded by the spike sorting pipeline were excluded from the analyses presented here if they failed one of the three criteria described in Ref.^104^ (the single unit computed metrics of RIGOR^24^): amplitude *>*50 *µ*V ; noise cut-off *<* 20 *µ*V; refractory period violation. Neurons that passed these criteria are termed ‘good’ neurons (or often just ‘neurons’) in this study. Out of the 621733 neurons collected, 75708 were considered good neurons. Final analyses were additionally restricted to regions that a) are designated gray matter in the adult mouse Allen common coordinate framework^5^, b) contained at least 5 good neurons per session, and c) were recorded from in at least 2 such sessions.

### Video analysis

We briefly describe the video analysis pipeline; full details can be found in Ref.^107^. The recording rigs contain three cameras, one called ‘left’ at full resolution (1280×1024) and 60 Hz filming the mouse from one side; one called ‘right’ filming the mouse symmetrically from the other side at half resolution (640×512) and 150 Hz; and one called ‘body’ at half resolution and 30 Hz filming the body of the mouse from above. We developed several quality control metrics to detect raw video issues such as poor illumination (as infrared light bulbs broke) or accidental misplacement of the cameras^107^.

We computed the motion energy (the mean across pixels of the absolute value of the difference between adjacent frames) of the whisker pad areas in the ‘left’ and ‘right’ videos (Fig. 1d). The whisker pad area was defined empirically using a rectangular bounding box anchored between the nose tip and the eye, both found using DeepLabCut^108^ (DLC; see more below). This metric quantifies motion in the whisker pad area and has a temporal resolution of the respective camera.

We also performed markerless pose estimation of body parts using DLC^29^, which is used within a fully automated pipeline in IBL (version 2.1) to track various body parts such as the paws, nose, tongue, and pupil (Fig. 1d). In all analyses using DLC estimates, we drop predictions with likelihood *<* 0.9. Furthermore, we developed several quality control metrics for the DLC traces^107^.

### Receptive field mapping

At the end of the behavioral task session, we performed 5 minutes of receptive field mapping experiment for most of the recordings (504/699 insertions). During the receptive field mapping phase, visual stimuli were random square pixels in a 15 × 15 grid occupying 120 degrees of visual angle both horizontally and vertically. There were three possible colors for each pixel: white, grey, and dark. The color of pixels randomly switched at the frame rate of 60 Hz, with an average duration of around 100 ms (Fig. S13a).

To compute the receptive field, we identified the moments when color switching occurred for each pixel. We defined the moments when color brightness increased as the “On” stimulus onset time, which includes the transition from dark to grey and grey to white. Similarly, we defined the moments when color brightness decreased as the “Off” stimulus onset time, which includes the transition from grey to dark and white to grey. We then computed the average spike rate aligned with On and Off stimulus onset for each pixel, from 0 to 100 ms. We defined two types of receptive fields, “On” and “Off”, as the average“On” and “Off” spike rates across pixels.

To estimate the significance of the receptive field, we fitted the receptive field to a 2d-gaussian function, then compared the variance explained to the fitting of a randomly shuffled receptive field (200 shuffles), and computed the p-value of significance. We defined a neuron as having a significant receptive field if either “On” or “Off” receptive field had *p <* 0.01.

### Assessing statistical significance

In this work, we studied the neural correlates of task and behavioural variables. In order to assess the statistical significance of these analyses, we need to account properly for spurious correlations. Spurious correlations can be induced in particular by slow continuous drift in the neurophysiological recordings, due to various factors including movement of the Neuropixels probes in the brain. Such slow drifts can create temporal correlations across trials. Because standard correlation analyses assume that all samples are independent, they can yield apparently significant nonsense-correlations even for signals that are completely unrelated^109, 110^.

Null distributions were generated against which we tested the statistical significance of our results. More specifically, we used distinct null distributions for each of the three types of variables we considered: a discrete behaviour-independent variable (the stimulus side), discrete behaviour-dependent variables (e.g., reward and choice), and continuous behaviour-dependent variables (e.g., wheel speed and wheel velocity). For the rest of the section, we will denote the aggregated neural activity across *L* trials and *N* neurons by *S* ∈ R*^L×N^*, and denote the vector of scalar targets across all trials by *C* ∈ R*^L^*.

For a discrete behaviour-independent variable, we generated the null distribution from so-called “pseudo-sessions”, i.e., sessions generated from the same generative process as the one used for the mice. This ensured that the time series of trials in each pseudo-session shares the same summary statistics as ones used in the experiment. We generated *M* (typically M = 200) pseudo-targets 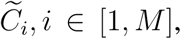 and performed the given analysis on the pair 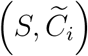 and obtained a fit score 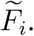 In pseudo-sessions, the neural activity *S* should be independent of 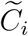 as the mouse did not see 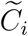 but rather *C*. Any predictive power from 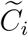 to *S* (or from *S* to 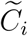) would arise, for instance, from slow drift in *S* unrelated to the task itself. These pseudo-scores 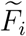 can be compared to the actual score *F* obtained from the neural analysis on (*S, C*) to assess statistical significance.

For discrete behaviour-dependent variables (such as choice or reward), we could not use the pseudo-session procedure above as we do not know the underlying generative process in the mouse. We therefore used “synthetic” sessions to create a null distribution. These depend on a generative model of the process governing the animals’ choices. In turn, this requires a model of how the animals estimated the prior probability that the stimulus appears on the right or left side of the screen, along with a model of its response to different contrasts given this estimated prior. In our companion paper on the subjective prior^23^, we found that the best model of the prior across all animals uses a running average of the past actions as a subjective prior of the side of the next stimulus, which we refer to as the ‘action-kernel’ model. The subjective prior *π_t_*follows the update rule:

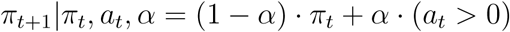

with *a_t_* ∈ {−1, 1} {(left, right)} the action performed by the mouse on trial *t* and *α* the learning rate, which we fitted on a session-by-session basis. This effectively models how mice use information from previous trials to build a subjective prior of where the stimulus is going to appear at the next trial. The details of how this prior is integrated with the stimulus to produce a decision policy is described in the companion paper^23^.

We fit the parameters of this model of the mouse’s decision-making behaviour separately for each session, and then created “synthetic” targets 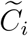 for that session by applying the model (with those fitted parameter values) to stimuli generated from pseudo-sessions to obtain time series of choice and reward. Then, as for the pseudo-sessions above, we obtained pseudo-scores 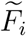 based on 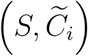 and assessed statistical significance by comparing the distribution of pseudo-scores to the actual score *F* obtained from the neural analysis on (*S, C*).

For the third type of variable – continuous behaviour-dependent variables such as wheel speed – generating synthetic sessions is harder, as we do not have access to a reasonable generative model of these quantities. We instead used what we call “imposter” sessions, generated from the continuous behaviour-dependent variable from another mouse on another session. More precisely, an imposter session for an original session of *L* trials is generated by performing the following steps:

1. concatenate trials across all sessions analysed in this paper (leaving out the session under consideration)
2. randomly select a chunk of *L* consecutive trials from these concatenated sessions
3. return the selected chunk, the imposter session

The continuous behaviour-dependent variable can then be extracted from the imposter session. As with the pseudo-sessions and the synthetic sessions, we obtained pseudo-scores 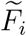 from a collection of imposter sessions and assess statistical significance by comparing the distribution of pseudo-scores to the actual score *F* obtained from the neural analysis on (*S, C*).

To apply the linear shift method ^111, 112^ to compare spiking with movement variables, we first truncated both the movement and spiking time series by removing N= 20 samples from both ends of both timeseries. We computed the Pearson correlation coefficient of the central segments, and compared the square of this coefficient to a null distribution obtained by repeatedly shifting the spiking timeseries linearly from the beginning to the end of the full behavioral timeseries. Significance was assessed using the approximate criterion, rejecting the null with significance *α* = 0.05 if the unshifted correlation was in the top *α*(2*N* + 1) of the shifted values.

Additional information about assessing statistical significance for individual analyses are detailed in the analysis-specific sessions below. For decoding, single-cell and population trajectory analyses, the results come in the form of per-region p-values. We used the false discovery rate to correct for comparisons across all the regions involved in each analysis (201 for the main figures), at a level of *q* = 0.01. We employed the Benjamini-Hochberg procedure^113^ as we expected substantial independence among the tests. As noted, we were not able to assess significance for the encoding analysis because of a lack of a convenient null distribution.

### Decoding: overview

We performed a decoding analysis to measure how much information the activity of populations of neurons contained about task variables such as stimulus side and choice. To do this we used cross-validated, maximum likelihood regression with L1 regularization (to zero out the contribution from noisy neurons). The neural regressors were defined by binning the spike counts from each neuron in each session in a given region within a specific time window on each trial. The duration of the time window, the number of bins in that time window (i.e. bin size), and the trial event to which it was aligned depended on the variable that is the target of our regression (Table 2). These are discussed further below and include a variety of behavioural and task variables: stimulus side, choice, feedback, and wheel speed/velocity. Although a session may have included multiple probe insertions, we did not perform decoding on these probes separately because they are not independent. Instead, neurons in the same session and region were combined across probes for our decoding analysis. Decoding was cross validated and compared to a null distribution to test for significance. A given region may have been recorded on multiple sessions, and thus in the main figures (Figs. 5, 6, 7, 8) the region *p*-value was defined by combining session *p*-values using Fisher’s combined probability test, and the region effect size was defined by subtracting the median of the null distribution from the decoding score and reporting the median of the resulting values across sessions. The *p*-values for all regions were then subjected to false discovery rate correction for multiple comparisons at level *q* = 0.01.

**Table 2.** Details of decoding analysis. Decoded variables are the targets for regression. Spike sorted activity is summed within trial-relative bins and used as the regressors (Fig. S4b). Regression is performed with the specified regression type using L1 regularization and the cross-validation scheme described in the text. Performance is reported on held-out trials using the specified score, and this score is compared to a null distribution of the form listed to evaluate statistical significance.

| Decoded variable | Trial-relative time | Window start (ms) | Window end (ms) | Bin size (ms) | Regression type | Score | Null distribution |
| --- | --- | --- | --- | --- | --- | --- | --- |
| Stim side | Stim onset | 0 | 100 | 100 | Logistic | bal. acc. | pseudo-session |
| Wheel speed | first wheel movement time | -200 | 1000 | 20 | Linear | $R^2$ | imposter |
| Wheel velocity | first wheel movement time | -200 | 1000 | 20 | Linear | $R^2$ | imposter |
| Choice | first wheel movement time | -100 | 0 | 100 | Logistic | bal. acc. | synthetic |
| Feedback | Feedback time | 0 | 200 | 200 | Logistic | bal. acc. | synthetic |

**Table 3.**
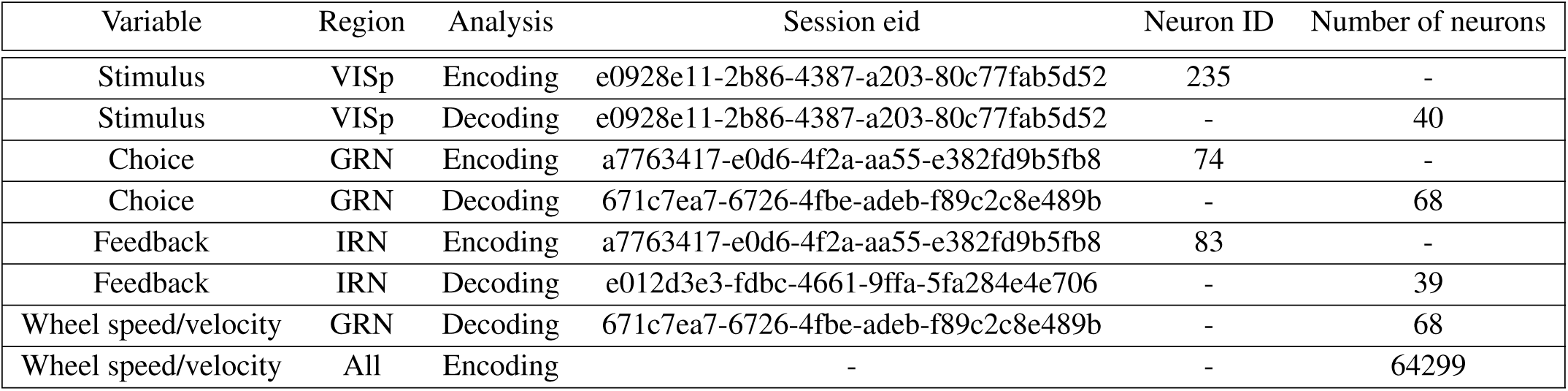
Session and neurons used for example encoding and decoding analysis. For the stimulus (Fig. 5), choice (Fig. 6), feedback (Fig. 7), and wheel speed/velocity (Fig. 8) we included figure panels showing example encoding and decoding analyses. Here, we list the variable name, region, and analysis type along with the session eid, encoding neuron ID number, and decoding number of neurons.

### Decoding: target variables

Stimulus side, choice, and feedback were treated as binary target variables for logistic regression. For stimulus side, trials which had zero contrast were excluded. We used the `LogisticRegression` module from scikit-learn^114^ (version 1.1.2) with 0.001 tolerance, 20000 maximum iterations, “l1” penalty, “liblinear” solver, and “fit intercept” set to True. We balanced decoder classes by weighting samples by the inverse of the class frequency, 1*/*(2*p_i,class_*). Decoding performance was evaluated using the balanced accuracy of classification, which is the average of the recall probabilities for the two classes. Fig. S26 shows histograms of the regression coefficients for all the variables.

Wheel values (speed and velocity) change over the course of a trial, unlike the previous decoding targets, and we had therefore to treat these target variables differently. We averaged wheel values in non-overlapping 20 ms bins, starting 200 ms before first wheel movement time and ending at 1000 ms after first wheel movement time. Spike counts were binned similarly. The target value for a given bin (ending at time *t*) was decoded from spikes in a preceding (causal) window spanning *W* bins (ending at times *t*, … , *t*-*W*). Therefore, if decoding from *N* neurons, there were (*W* + 1)*N* predictors of the target variable in a given bin. In practice we used *W* = 10. To decode these continuous-valued targets we performed linear regression using the `Lasso` module from scikit-learn^114^ (version 1.1.2) with 0.001 tolerance, 1000 maximum iterations, and “fit_intercept” set to True. Decoding performance was evaluated using the *R*^2^ metric.

### Decoding: cross validation

We performed all decoding using nested cross-validation. Each of five “outer” folds was based on a training/validation set comprising 80% of the trials and a test set of the remaining 20% of trials. We selected trials at random in an “interleaved” manner. The training/validation set of an outer fold was itself split into five “inner” folds, again using an interleaved 80%/20% partition. When logistic regression was performed, the folds had to be selected such that the trials used to train the decoder included at least one example of each class. Because both outer and inner folds were selected at random, it was possible that this requirement was not met. In those circumstances, we re-sampled the outer or inner folds. Likewise, we disallowed pseudo/synthetic sessions which had too few class examples. We fit regression models on the 80% training set of the inner fold using regularization coefficients {10*^−^*^5^, 10*^−^*^4^, 10*^−^*^3^, 10*^−^*^2^, 10*^−^*^1^, 10^0^, 10^1^} for logistic regression (input parameter *C* in sklearn) and {10*^−^*^5^, 10*^−^*^4^, 10*^−^*^3^, 10*^−^*^2^, 10*^−^*^1^} for linear regression (input parameter *α* in sklearn). We then used each model to predict targets on the remaining 20% of the trials of the inner fold, i.e. the validation set. We repeated this procedure such that each trial in the original training/validation set of the outer fold is used once for the validation set and four times for the train set. We then took the regularization coefficient that performed best across all validation folds and retrained a regression model using all trials in the training/validation set of the outer fold. This final model was used to predict the target variable on the 20% of trials in the test set of the outer fold. We repeated the above train/validate/test procedure five times, each time holding out a different 20% of test trials such that, after the five repetitions, each trial had been included in the test set exactly once, and included in the training/validation set exactly four times. The concatenation of all test set predictions, covering 100% of the trials, was used to evaluate the decoding score.

We found that for some regions and sessions,the resulting decoding score was sensitive to the precise assignment of trials to different folds. Therefore, to provide additional robustness to this procedure, we repeated the full five-fold cross-validation over multiple separate runs, each of which used a different random seed for selecting the interleaved training/validation/test splits. We then took the average decoding score across all runs as the final reported decoding score. When decoding stimulus side, choice, and feedback we performed ten runs, and while decoding wheel speed and wheel velocity, we used two runs due to the added computational burden of decoding the wheel values which include multiple bins per trial.

In order to further reduce the sensitivity of decoding scores due to fold allocation, the companion prior paper^23^ requires a minimum of 250 trials in order to perform decoding of a given session. We waive that requirement for the decoding analyses in this paper in order to match the same neurons used in the other analyses. We find that relaxing this requirement only affects the significance of a small number of regions for each target variable (Fig. S27).

### Decoding: significance testing with null distributions

We assessed the significance of the decoding score resulting from the multi-run cross-validation procedure by comparing it to those of a bespoke null distribution of decoding scores. To construct appropriate null distributions, we fixed the regressor matrices of neural activity and generated new vectors of target values that follow similar statistics (Table 2), as described above. Once the new target values were generated, we carried out the full multi-run cross validation procedure described above to get a new decoding score. This was repeated multiple times to produce a null distribution of decoding scores: stimulus side, choice, and feedback were repeated 200 times, while wheel speed and velocity were repeated 100 times to reduce the computational burden.

The null distribution was used to define a *p*-value for each region-session pair, where the *p*-value was defined as 1 − *ρ* where *ρ* was the percentile relative to the null distribution. Each brain region was recorded in ≥ 2 sessions, and we employed two different methods for summarizing the decoding scores across sessions: a) the median corrected decoding score amongst sessions which was used as the effect size in the main figures; the values were corrected by subtracting the median of the decoding score of the null distribution; b) the fraction of sessions in which decoding was significant, that is if the *p*-value was less than *α* = 0.05, which is shown in the supplement. We combined session-wide *p*-values using Fisher’s combined probability test (also known as Fisher’s method^30, 31^) when computing a single statistic for a region. Finally, the combined *p*-value for a region was subjected to a false discovery rate correction for multiple comparisons at level *q* = 0.01. We note the combined *p*-value may be significant, but the computed effect size may be negative. This is because many sessions used for decoding in that region may have been insignificant, driving the effect size down, while a small number of sessions may have been significant causing Fisher’s combined probability test to produce a significant combined *p*-value.

### Single-cell correlates of sensory, cognitive, and motor variables

We quantified the sensitivity of single neurons to three task variables: the visual stimulus (left versus right location of the visual stimulus), choice (left versus right direction of wheel turning), and feedback (reward versus non-reward). We computed the sensitivity metric for each task variable using the condition combined Mann-Whitney U statistic^15, 115, 116^ (Fig. S4a,c). Specifically, we compared the firing rates from those trials with one task-variable value *V*_1_ (e.g., trials with stimulus on the left side) to those with the other value *V*_2_ (e.g., with stimulus on the right side), while holding the values of all other task variables fixed. In this way, we could isolate the influence of individual task variables on neural activity. To compute the U statistic, we first assigned numeric ranks to the firing rate observations in each trial. We then computed the sum of ranks *R*_1_ and *R*_2_ for the observations coming from *n*_1_ and *n*_2_ trials associated with the task-variable values *V*_1_ and *V*_2_, respectively. The U statistic is defined as:

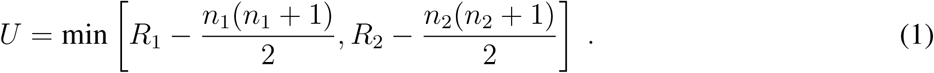

The probability that the firing rate on *V*_1_ trials is different (greater or smaller) from the firing rate on *V*_2_ trials is computed as 1 − *P*, where *P* is given by

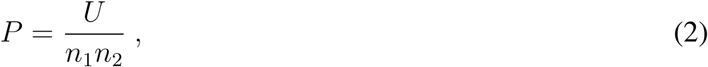

which is equivalent to the area under the receiver operating characteristic curve^117, 118^. The null hypothesis is that the distributions of firing rates on *V*_1_ and *V*_2_ trials are identical.

To obtain a single probability across conditions, we combined observations across different trial conditions *j* by a sum of U statistic in these conditions^15^:

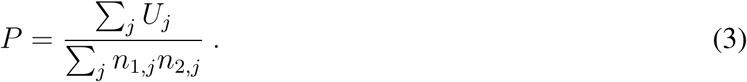

Here *n*_1*,j*_ and *n*_2*,j*_ are the numbers of *V*_1_ and *V*_2_ trials, respectively, within the condition *j*.

For the visual stimulus, we compared firing rate on trials with the stimulus on the left versus stimulus on the right during time-window [0, 100] ms aligned to the stimulus onset time. For choice, we compared firing rate on trials with the left versus right choice during time-window [−100, 0] ms aligned to the first wheel movement time. For the feedback, we compared firing rate on trials with reward versus non-reward during time-window [0, 200] ms aligned to the feedback onset time.

To estimate statistical significance, we used a permutation test in which trial labels for one task variable were randomly permuted 3,000 times within each subset of trials with fixed values of all other task variables, and the Mann-Whitney U statistic was computed for each permutation. We computed the *p*-value for each task variable as the fraction of permutations with the statistic *P* greater than in the data. This approach controls for correlations among task variables and allows us to isolate the neuron’s sensitivity to stimulus that is not due to sensitivity to block and choice, and vice versa. Random permutations, however, do not control for spurious correlations that can arise due to autocorrelations in the time series of the firing rate and task variable^109^. To control for spurious correlations, we used a within-block permutation test to control simultaneously for both temporal correlations and correlations among task variables. Specifically, we generated the null distribution by randomly permuting trial labels with fixed values of all other task variables within each individual block, which effectively reduces the serial dependencies of task variables at the timescale of block duration.

The combined condition Mann-Whitney U statistic is known to have a relatively high false positive rate due to the limited number of trials in each condition. To obtain a sufficient number of trials, we also computed a simple Mann-Whitney U statistic without separating different conditions. We defined *p <* 0.001 (*α_MW_* = 0.001) as the criterion of significance for the simple Mann-Whitney U statistic, and *p <* 0.05 (*α_CCMW_* = 0.05) for the combined condition Mann-Whitney U statistic. We defined neurons that were statistically significant in both tests to be sensitive neurons for a specific task variable.

To quantify the overall responsiveness of single neurons to the behavioural task, we used the Wilcoxon rank-sum test to compare firing rates between the baseline ([−200, 0] ms window aligned to the stimulus onset) and different task periods: [50, 150] ms and [0, 400] ms aligned to stimulus onset, [−100, 50] ms and [−50, 200] ms aligned to the first wheel movement time, and [0, 150] ms aligned to the reward delivery. These time windows are selected based on the test of responsiveness in previous work on large-scale neural coding with a very similar task structure^15^.

To measure the behavioural movement correlates of single neurons in the entire recording sessions, we computed zero time-lag Pearson correlation coefficient between time-series of spike counts in 50 ms bins and time-series of four behavioural variables (nose, pupil, paw, and tongue) each extracted from videos of the subject using DeepLabCut software^29^. To assess the significance of these correlations, we applied a time-shift test^78^ and computed 2*K* = 40 time-shifted correlations varying the offset between time-series of spiking activity and behavioural variables from 50 to 1000 ms (both positive and negative offsets). We then counted the number of times *m* where the absolute value of time-shifted correlation exceeds that of zero time-lag correlation and assigned the *p*-value as the fraction of the absolute value of permuted correlations greater than in the data *p* = *m/*(*K* + 1). We then assigned each neuron as being significantly responsive relative to a particular threshold on this *p*-value.

We then computed the fraction of neurons in each brain region that were significantly responsive to the behavioural task, movement, visual stimulus, choice, and feedback, and identified brain regions that were most responsive to these conditions. Specifically, for each region, we computed the *p*-value of the fraction of neurons (*f_i_*) in i-th session by comparing the fraction to a binomial distribution of fractions due to false positive events: Binomial(*N_i_, α*), where *N_i_* is the number of neurons in i-th session, and *α* is the false positive rate:

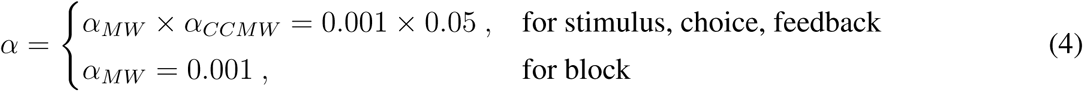

We defined the *p*-value *p_i_* as the probability of the fraction *f_i_* that is larger than the distribution Binomial(*N_i_, α*). Next, we used Fisher’s combined probability test to compute a combined *p*-value of each brain region by combining the *p*-values of all sessions (*i* = 1, 2*, …m*). After computing combined *p*-values of each brain region, these *p*-values were then entered into the false discovery rate (FDR) procedure (Benjamini–Hochberg procedure) at level *q* = 0.01 to correct for multiple comparisons. We defined a list of regions to be significant based on this FDR procedure.

### Population trajectory analysis methods

We examined how responsive different brain regions were to a task variable *v* of interest. To do so, we constructed a pair of variable-specific “supersessions” 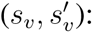 We partitioned all the IBL data into two, corresponding to the opposing pair of conditions for the variable (e.g. for stimulus discrimination, we split the trials into the L and R stimulus conditions) and replaced the trial-by-trial responses of each cell within the condition and within each session with one trial-averaged response (Fig. 4e). These trials were aligned to a variable-specific reference time (e.g., stimulus onset time for stimulus discrimination). We used the canonical time windows shown in Fig. 4a around the alignment time for the main figures unless stated otherwise (e.g. for feedback we used a longer time window in the temporal evolution plot to illustrate licking), time bins of length 12.5 ms, and stride 2 ms. The supersessions *S_v_,* 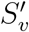 had a number of rows equalling the number of IBL sessions passing quality control for that variable condition times the number of cells per session; columns correspond to time-bins.

We then subdivided the supersessions by brain region 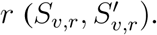 These defined a pair of across-IBL response trajectories (temporal evolution of the response) to the pair of variable *v* conditions for each brain region.

We next computed the time-resolved difference in response of brain region *r* to the opposing conditions of task variable *v*. We restricted our analyses to regions with ≥ 20 rows in 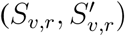 for all analyses. Our primary distance metric, which we call *d_v,r_*(*t*), was computed as a simple Euclidean distance in neural space, normalized by the square root of the number of cells in the given region.

Given a time-resolved distance curve, we computed the maximum and minimum distances along the curve to define a variable- and region-specific modulation amplitude:

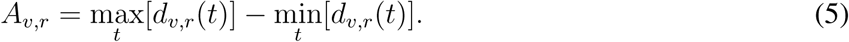

We obtained a variable- and region-specific response latency by defining it as the first time *t* at which *d_v,r_*(*t*) = min*_t_*[*d_v,r_*(*t*)] + 0.7(max*_t_*[*d_v,r_*(*t*)] − min*_t_*[*d_v,r_*(*t*)]). Using modulation amplitude as a measure of effect size, we then quantified the combined modulation amplitude and latency of regions as a function of task variable.

To generate a significance measure for the variable- and region-specific distance measures, we used a pseudo-trial method for generating null distance distributions, as described below. Distances were significant if they were greater in size than the corresponding null distance distribution with *p <* 0.01. Although the statistical significance of regions was therefore controlled for the effects of other task variables, note that the distance amplitudes and latencies were not.

Below we list the three task variables examined and the associated null distributions:

- “Stimulus” supersession: *S_v_,* 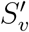 correspond to trials with the stimulus on the left or right, respectively, aligned by stimulus onset time and including 0 ms before to 150 ms after onset. To generate pseudo-trials, we permuted the stimulus side labels among trials that share the same block and choice side.
- “Choice” supersession: *S_v_,* 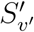 correspond to trials with the animal’s response (wheel movement) to the left or right, respectively, aligning by first wheel movement time and including 0 ms before to 150 ms after onset. To generate pseudo-trials, we permuted the choice labels among trials with the same block and stimulus side.
- “Feedback” supersession: *S_v_,* 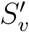 correspond to trials where the animal’s response was correct (recall that the feedback was water delivery) or incorrect (recall that the feedback was tone and timeout delivery), respectively, aligning by feedback onset including 0 ms before to 150 ms after onset. To generate pseudo-trials, we permuted the choice labels among trials with the same block and stimulus side and then compared these pseudo-choices with the true stimulus sides to obtain pseudo-feedback types.

For each *S_v,r_,* 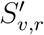 pair we repeated the pseudo-trial process M (=1000) times, then followed the same distance computation procedures described above to obtain a null distribution of M modulation amplitude scores. We obtained a *p*-value by counting *n* (as the number of pseudo-scores that were greater than the true score for this region) as: 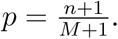

For regions with significant and large effect sizes to a given variable, we generated visualizations of the population dynamics by projecting the trajectories in *S_v,r_,* 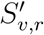 into a low-dimensional subspace defined by the first three principal components of the pair *S_v,r_,* 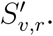 In addition to the main figure results, population trajectory results on the maximal dataset are shown in Fig. S16.

### Multiple linear regression model of single neuron activity

We fit linear regression models to single neuron activity, measured as spikes binned into 20 ms intervals. These models aim to express {*s_lt_*}, the neural activity in time bin *t* ∈ [1*, T*] on trial *l* ∈ [1*, L*] based on *D* time-varying task-related regressors *X* ∈ R*^L,T,D^*. We first represented the regressors across time using a basis of raised cosine “bump” functions in log space^119^. Each basis function was associated with a weight in the regression model, with the value of the basis function at time *t* described by 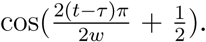 The basis functions were computed in log space and then mapped into linear time to more efficiently capture both fast neuronal responses in the *<*100 ms range and slow changes beyond that time (Fig. 4c). The width *w* and centre *τ* of each basis were chosen to ensure even coverage of the total duration of the kernel. In an example kernel with 3 bases, 3 separate weights would be fit to the event in question with weights describing early, middle, and late activity predicted by the event. These bases were convolved with a vector describing the effects of each regressor. In the case of timing events, the bases were convolved with a Kronecker delta function, resulting in a copy of the kernel at each time when the event occurred. We describe the simple case that each regressor enjoys the same number *B* of basis functions. This produced a new regression tensor 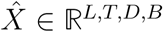

We then sought regression weights ***β*** ∈ R*^D,B^* such that, as closely as possible, 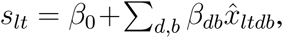 where {*β_db_*} are linear regression weights. Each single-neuron model used regressors for stimulus onset (left and right separately), first wheel movement time (L/R), correct feedback, incorrect feedback, value of the block probability, movement initiation, and wheel speed. Fitting was performed using an L2-penalised objective function (as implemented in the scikit-learn python ecosystem as 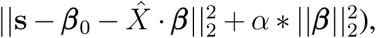 with the weight of the regularization *α* determined via cross-validation. Note that the intercept of the model is not included in the regularization, in order to capture fully the mean of the distribution of **s**.

We used a kernel composed of 5 basis functions to parameterise left and right stimulus onset, and correct and incorrect feedback. These bases spanned 400 ms, and corresponded to 5 weights per regressor for each of these 4 regressors in the model.

Previous work has shown that difficulty in perceptual decision-making tasks^120^, along with neural responses, do not change linearly with contrast. To account for this we modulated the height of the stimulus onset kernels as a function of contrast *c* with height 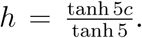 The resulting kernels would produce a response that was lower at low contrasts for the same set of weights {*β_db_*}.

To capture statistical dependencies between wheel movements and spiking, we used anti-causal kernels (in which the convolution of signal and kernel produces a kernel peak before peaks in the signal) describing the effect of first wheel movement time for leftward and rightward movements. These kernels described 200 ms of activity preceding first movement using 3 basis functions. We also used an additional anti-causal kernel of 3 bases covering 300 ms describing the effect of wheel speed, and was convolved with the trace of wheel speed for each trial. With these regressors we aimed to capture preparatory signals preceding movements related to the wheel.

Models were fit on a per-neuron basis with the L2 objective function using 5-fold cross-validation. Trials for cross-validation were chosen from a uniform distribution, and not in contiguous blocks. Models were then fit again using a leave-one-out paradigm, with each set of regressor weights *β_d_*_1_ *… β_dB_*being removed as a group and the resulting model fit and scored again on the same folds. The change between the base model score 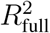 and the omission model 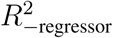 was computed as 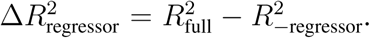 Additionally, the sensitivity for several pairs of associated regressors, such as left/right stimulus onset and correct/incorrect feedback, were defined as 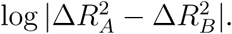 This computation was applied to: right/left stimulus, right/left first wheel movement time, and correct/incorrect feedback.

### Granger analysis across simultaneously recorded regions

Granger causality has been suggested as a statistically principled technique to estimate directed information flow from a pair of time series^121^. We used nonparametric spectral Granger causality^122^, implemented in python^123^, to compute a Granger score for all simultaneously recorded region pairs in the IBL’s brain-wide dataset.

For a given session, binned spikes (12.5 ms bin size) from both probes were averaged across regions to obtain a firing rate time series for the complete recording (excluding regions with fewer than 10 neurons per recording). Those series (typically 1.5 h long) were then divided in non-overlapping (10 s segments (irrespective of task contingencies or alignment), resulting in data input of shape #regions × #segments × #observations from which a Granger score as a function of frequency was computed for each directed region pair with the Spectral Connectivity python package^123^. We obtained a single Granger score per directed region pair by averaging across frequencies ^124^.

Statistical significance for a Granger score and region pair for a given session was established by a permutation test. That is, a null distribution of pseudo Granger scores was obtained by swapping the two region labels randomly across segments. 1000 of these pseudo scores were computed and a *p*-value obtained by counting the number of pseudo scores that were greater than the true Granger score and dividing this count by the number of pseudo scores plus 1. *p*-values across all Granger scores were corrected for multiple comparison via the Benjamini-Yekutieli method. Measurements were combined across sessions by taking the mean Granger score and using Fisher’s combined probability test to combine the *p*-values.

### Visualization and comparison of results across neural analyses

To facilitate the comparison of neural analyses across brain regions, for each task variable we visualised effect sizes in a table (e.g. Fig. 5f) specifying the effect size for each analysis and brain region. Cells of the table were coloured according to effect size using the same colour map as in the corresponding flatmap. Before summing, the effect sizes for each analysis were normalised to lie in the interval [0, 1]. This method highlights regions with large effects across all analyses, and indicates the extent to which the analyses agree. For a direct comparison of analyses scores, see flatmaps in Fig. S6 and scatter plots of scores for analysis pairs in Fig. S7.

## Example sessions and cells in main figures

## Acknowledgements

This work was supported by grants from the Wellcome Trust (216324), the Simons Foundation, The National Institutes of Health (NIH U19NS12371601), the National Science Foundation (NSF 1707398), the Gatsby Charitable Foundation (GAT3708), and by the Max Planck Society and the Humboldt Foundation. Part of the data analysis for this project was performed on Stanford University’s Sherlock cluster; we would like to thank Stanford University and the Stanford Research Computing Center for providing computational resources and support that contributed to these research results. Another part was performed at University of Geneva on “Baobab” and “Yggdrasil” high performance computing clusters. We also acknowledge computing resources from Columbia University’s Shared Research Computing Facility project, which is supported by NIH Research Facility Improvement Grant 1G20RR030893-01, and associated funds from the New York State Empire State Development, Division of Science Technology and Innovation (NYSTAR) Contract C090171. We thank Peter Latham, Tom Mrsic-Flogel and IBL colleagues for helpful comments on the manuscript. The production of all IBL Platform Papers is led by a Task Force, which defines the scope and composition of the paper, assigns and/or performs the required work for the paper, and ensures that the paper is completed in a timely fashion. The Task Force members for this platform paper include authors AP, BG, BB, CF, CL, DB, FH, GAC, IRF, JMH, KDH, KZS, MRW, MC, MS, NJM, NAS, OW, PD, TAE, YS.

## Supplementary information

**Figure S1.**
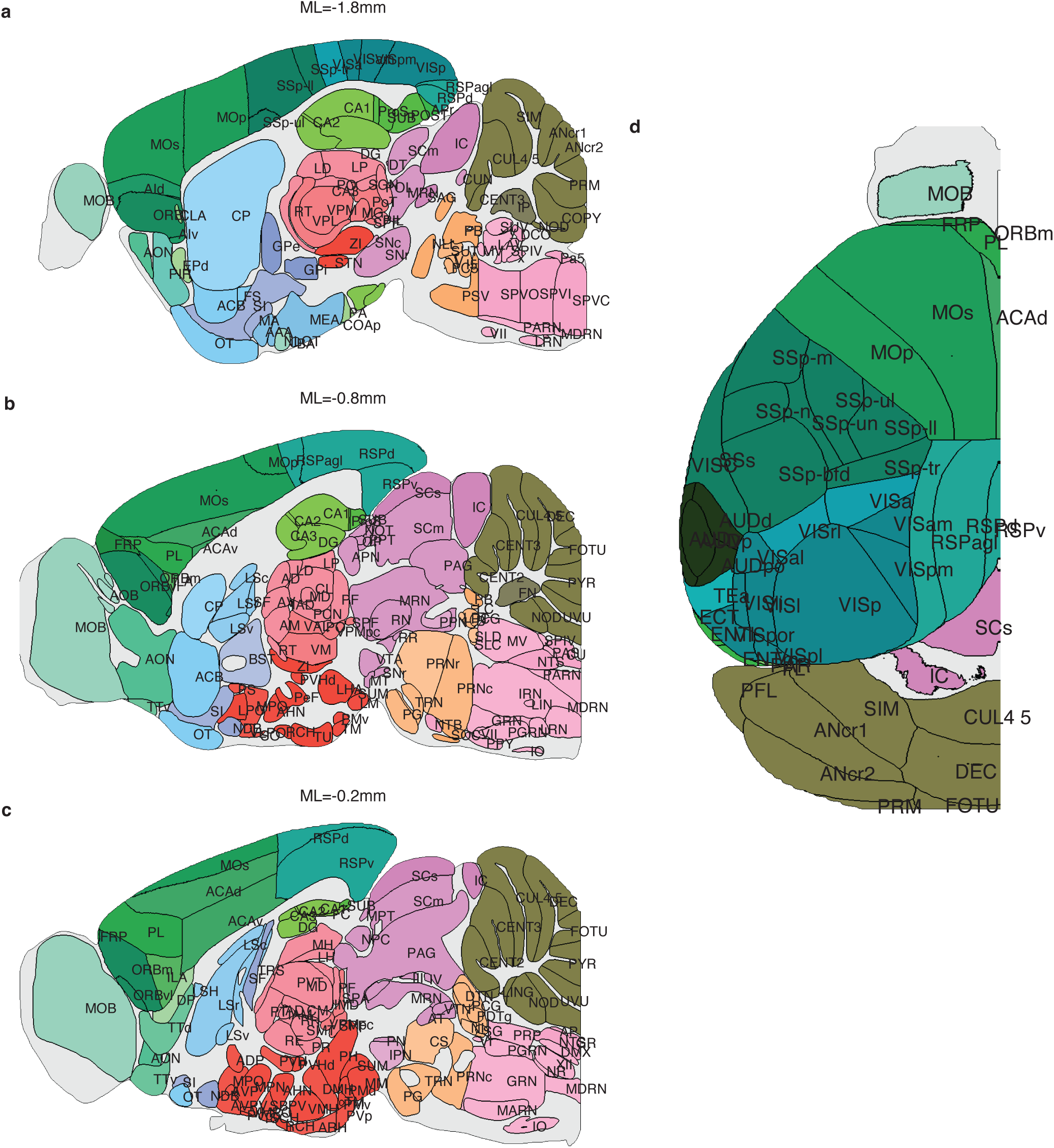
2d-brain slices maps annotated with region acronyms. **a)** Region acronyms for sagittal slices with coordinates: ML=-1.8mm, **b)** ML=-0.8mm, **c)** ML=-0.2mm. **d)** Region acronyms for the top view of the dorsal cortex.

**Figure S2.**
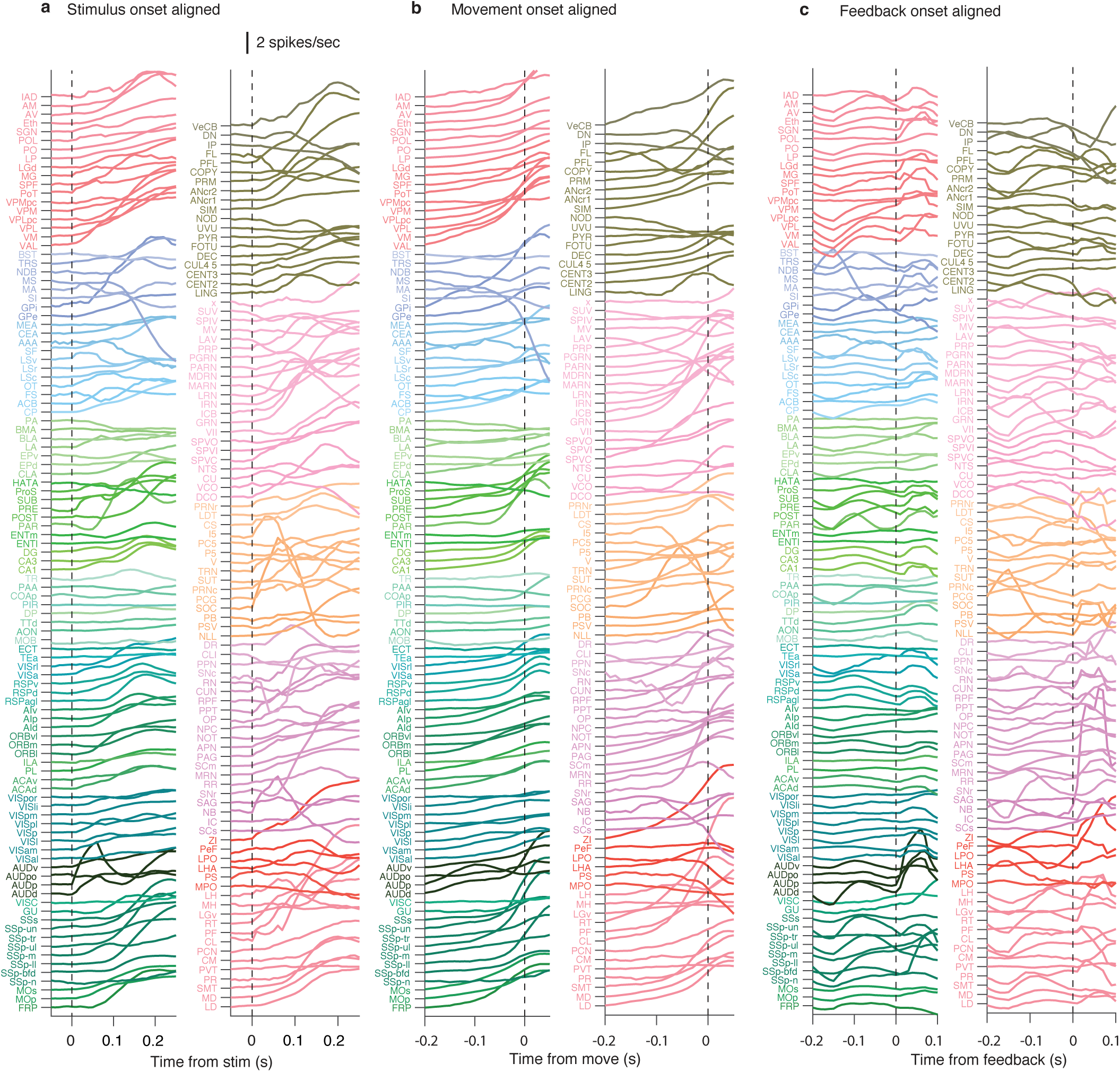
PSTH of average neural activity across the brain. **a)** PSTH of average neural activity aligned to stimulus onset, **b)** first wheel movement time, and **c)** Feedback onset.

**Figure S3.**
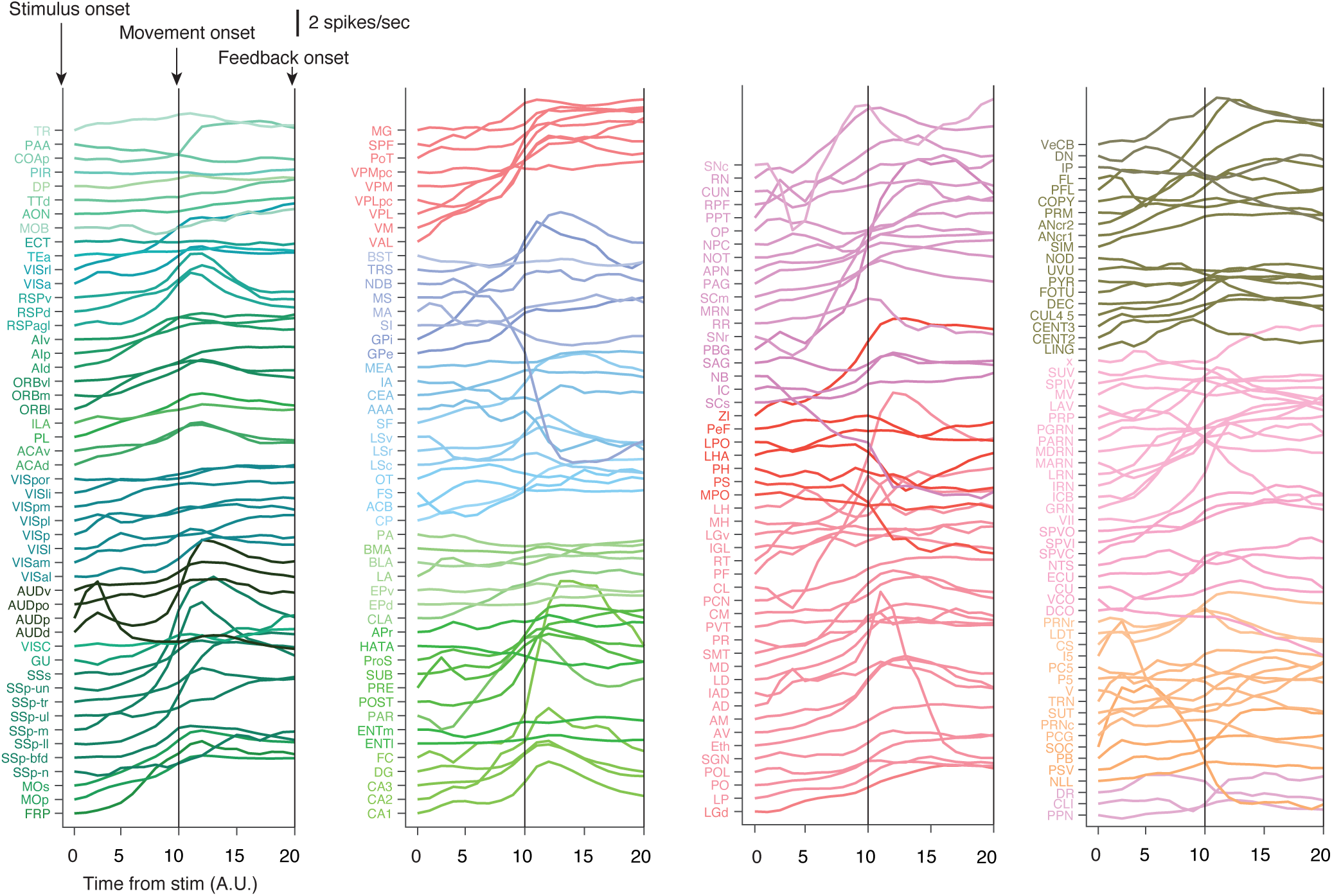
Time-warped PSTH of average neural activity across the brain. The duration between stimulus onset and first wheel movement time is divided into 10 equal-size time bins, as is the duration between onset to first wheel movement time and feedback onset for each trial (thus the length of time bin varies trail by trial). The spike rate of each time bin is computed and averaged across trials and sessions. This approach ensures stimulus, movement, and feedback onsets are perfectly aligned across trials.

**Figure S4.**
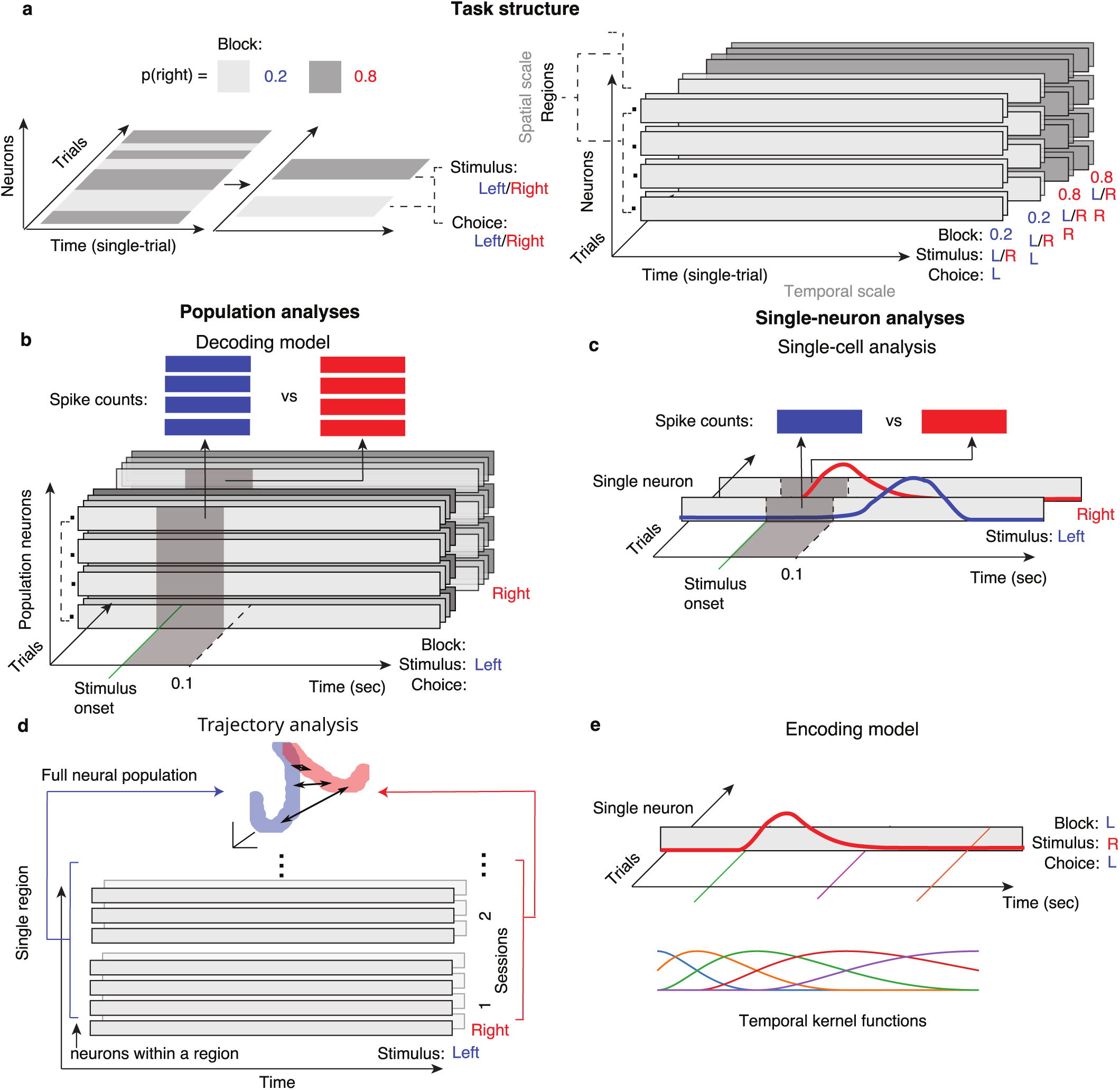
Detail of spatiotemporal structure of neural analyses. **a)** Task structures. In each session, consecutive trials form a block structure with the probability of a right-side stimulus being 0.2 and 0.8. In each block, there are trials with stimulus and choice side that are left or right. By regrouping trials, we can obtain 8 categories of trials with different combinations of stimulus side, block identify, and choice side. **b)** The decoding model studies the what can be extracted from the population neural activity about individual task variables within short time windows (without marginalizing other variables). **c)** Single-cell analysis studies the modulation of single-neuron activity by individual task variables within a short time window. **d)** Population trajectory analysis combines neural responses across multiple sessions (each neuron averaged across trials within a session) and analyses the trajectory of population neural activity. **e)** The encoding model uses temporal kernel functions to describe single-neuron activity during the entire trial at high temporal resolution.

**Figure S5.**
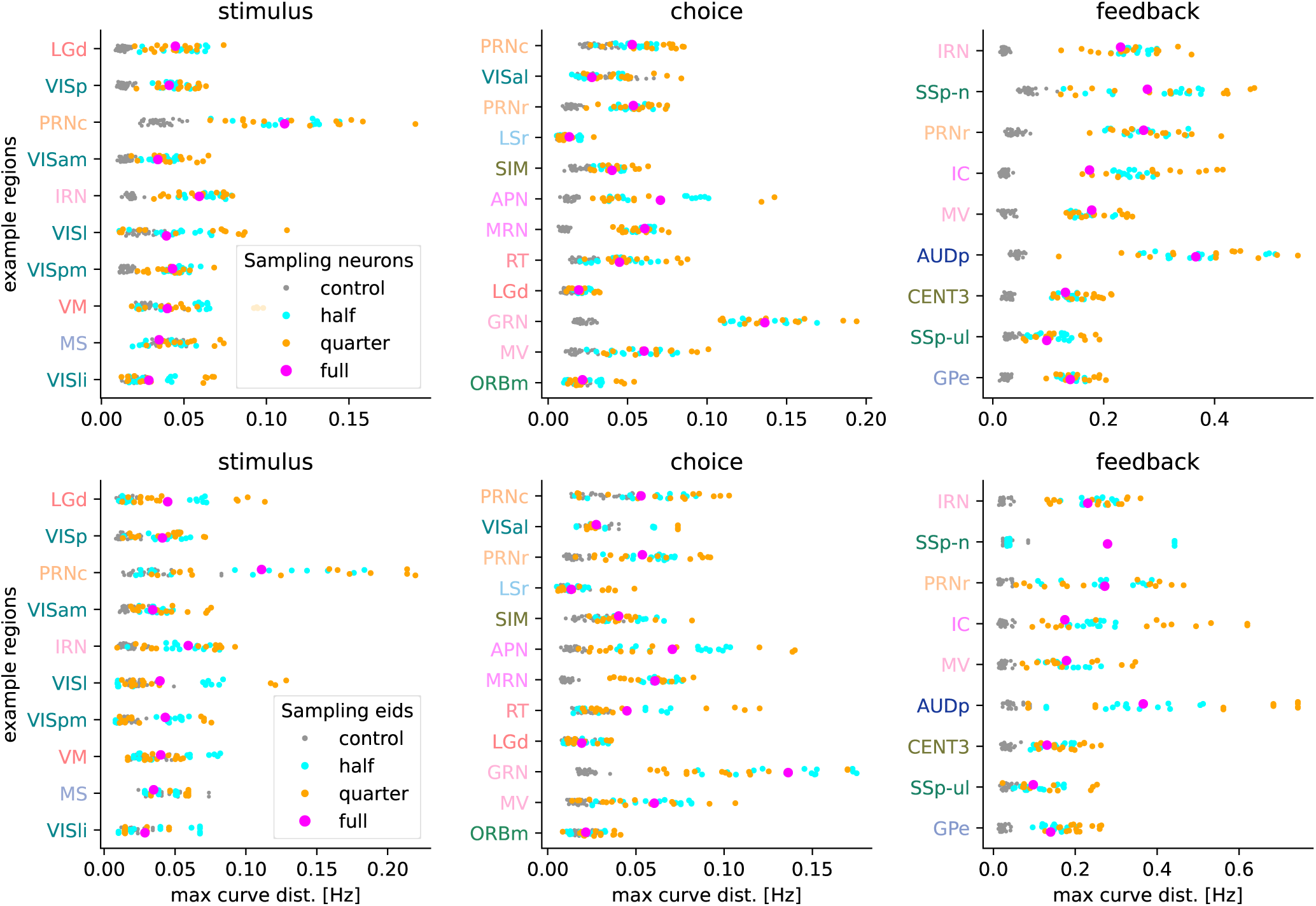
Maximal Euclidean distances for example regions with random data subsets. Each plot shows maximal population trajectory distances for the same highlighted regions as in the main figures (5l, 6l, 7l). The distances are computed after 15 subsamplings from half or a quarter of all neurons (top row of panels) or sessions (bottom row of panels). Grey dots indicate control scores for the sampled data, from trial randomization as in the main population trajectory analysis. For both, sampling neurons or sampling sessions, and most regions, the mean of the sampled scores visually matches the scores of the full dataset, showing that the strongest regional differences are also present for subsets of the data, however with more variance.

**Figure S6.**
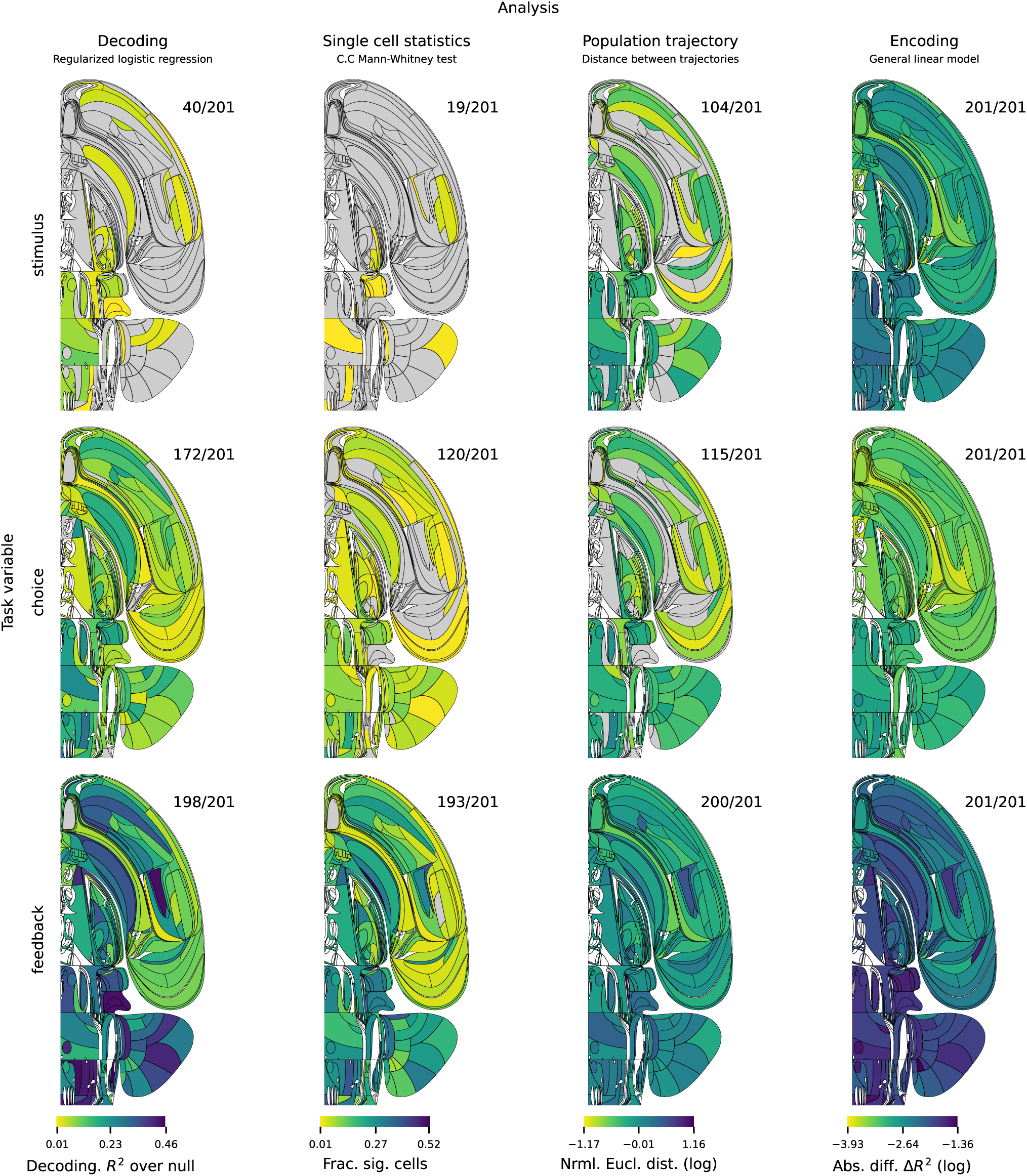
Comparison of effect sizes across task variables. Each column corresponds to a particular neural analysis and each row a task variable. For each analysis, the colour scale is fixed across all variables to enable comparison of effects between variables. For most analyses, the feedback variable has the largest effect amongst all task variables. The numbers at the top right indicate the fraction of significant regions across all analysed regions.

**Figure S7.**
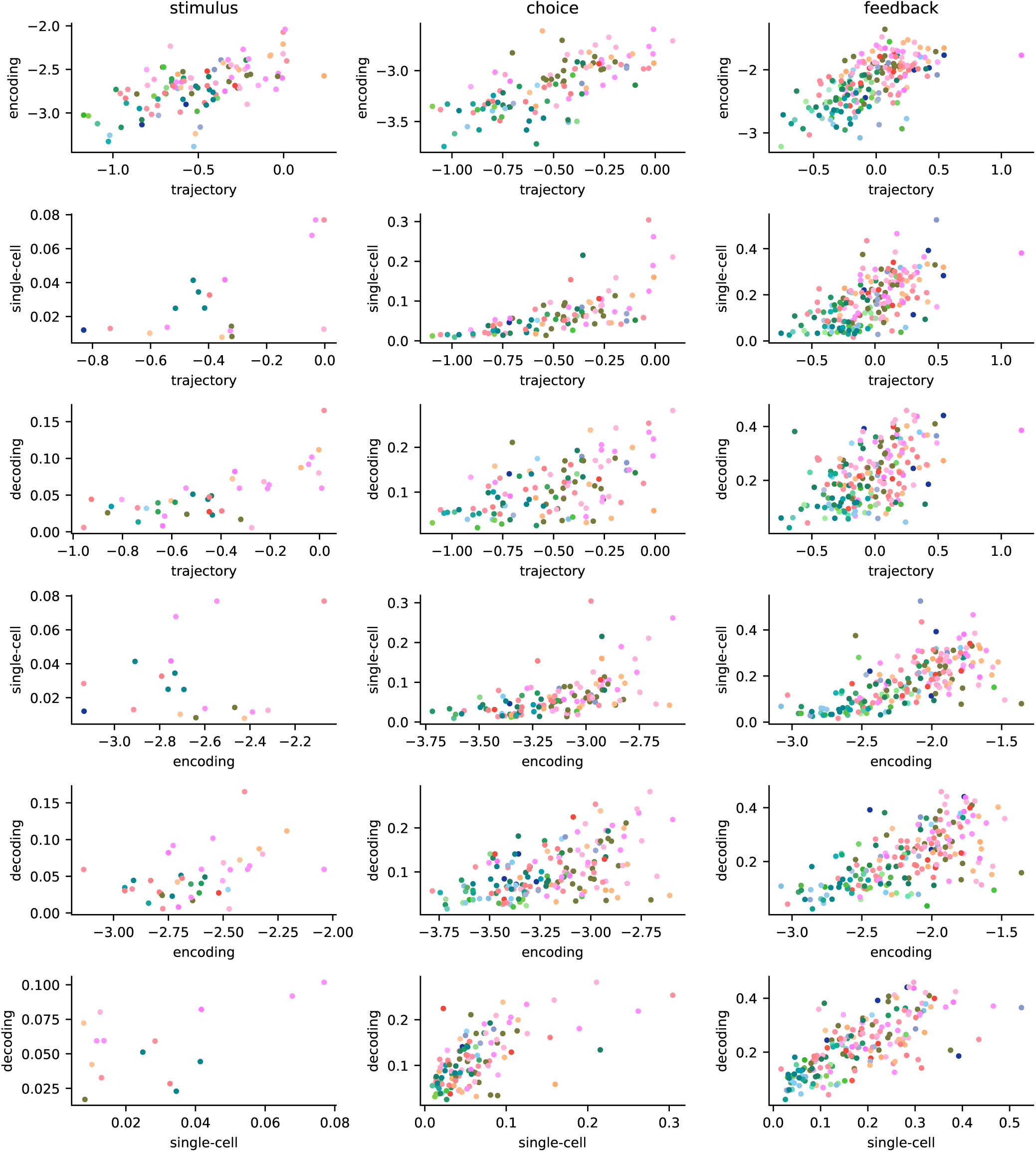
Amplitudes of analysis pairs for the three main variables. For a given analysis pair, say encoding and population trajectory, and a variable, say stimulus, all regions for which both analyses were significant are shown as dots in a scatter plot with the amplitudes as coordinates, colored using our canonical region coloring. There are 6 possible analysis pair combinations (rows) and 3 main variables (columns).

**Figure S8.**
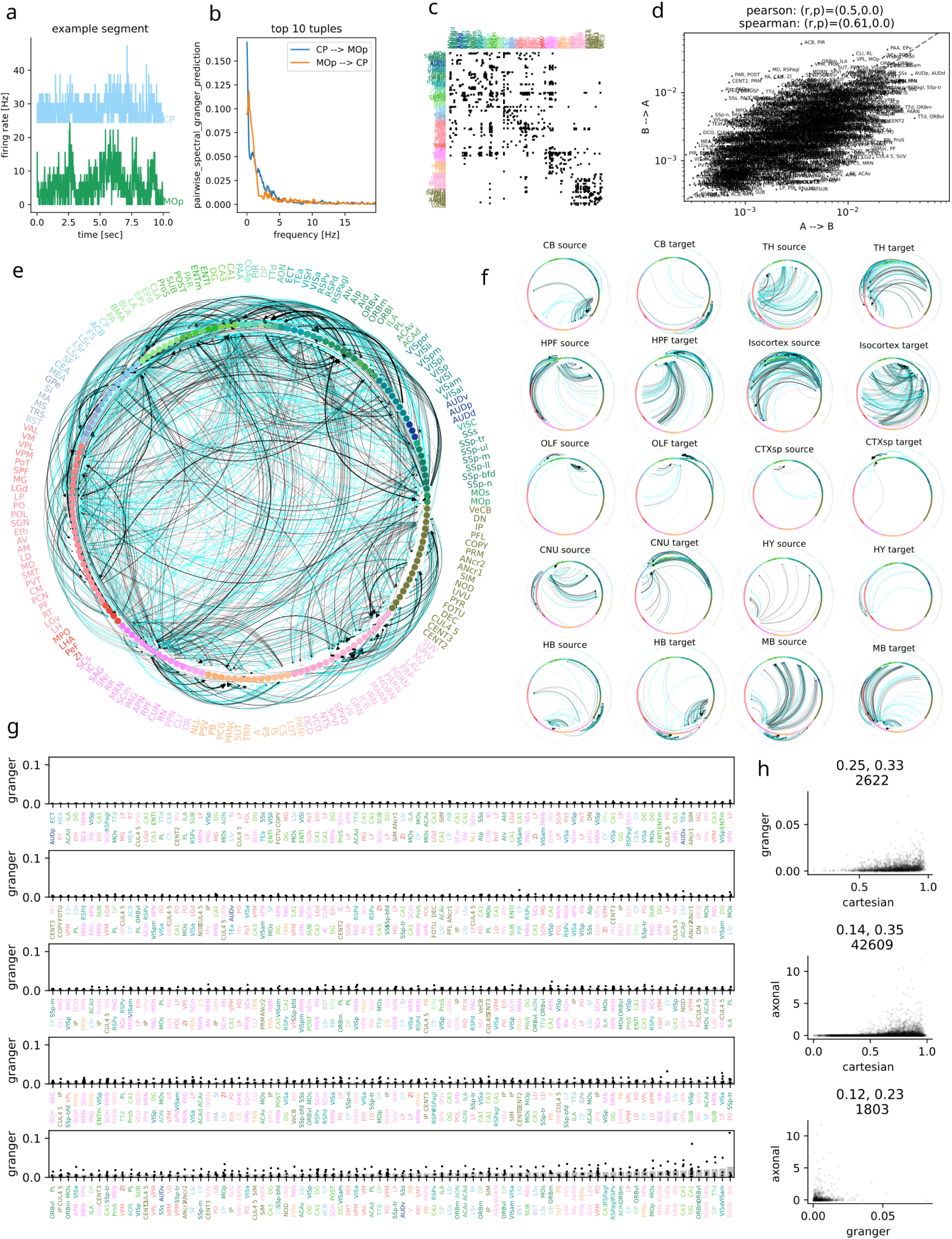
Granger scores for simultaneously recorded region pairs. **a)** Firing rates in two regions (CP and MOp) for an example session (eid = af55d16f-0e31-4073-bdb5-26da54914aa2); first 10 sec of recording. **b)** Directed spectral Granger prediction for an example region pair from this example session as a function of frequency. This is the average across consecutive 10 sec windows of the whole recording, irrespective of trial-structure. The mean Granger prediction across frequencies is the Granger score, used in all other panels. **c)** Binarised significant Granger score adjacency matrix, canonical region ordering (as in circular graph plot). Note the near-symmetry. **d)** Symmetry of Granger scores for all significant region pairs, log scale. Correlation scores in panel title. **e)** Granger scores for region pairs as averages across recordings, edge width proportional to Granger score, black if significant. Only region pairs with at least 2 recordings are shown. **f)** Graph of **e)** restricted to incoming/outgoing Granger scores for subsets of regions (Cosmos hierarchical level). **g)** Significant Granger scores for all region pairs, black dots are individual recordings, gray bars are mean across recordings, ordered by mean. Only region pairs with at least 3 recordings are shown. **h)** Granger scores in relation to two other connectivity metrics: axonal (axonal projection tracing, Figure 3 in ^125^) and cartesian (inverse Euclidean distance between centroids of region pairs). Weak but significant correlations (Pearson, Spearman, on top of panels, together with number of directed region pairs for the plot) are found for cartesian/Granger (.25, .33), cartesian/axonal (.14, .35) and Granger/axonal (.12, .23). All results are further listed in this online table.

**Figure S9.**
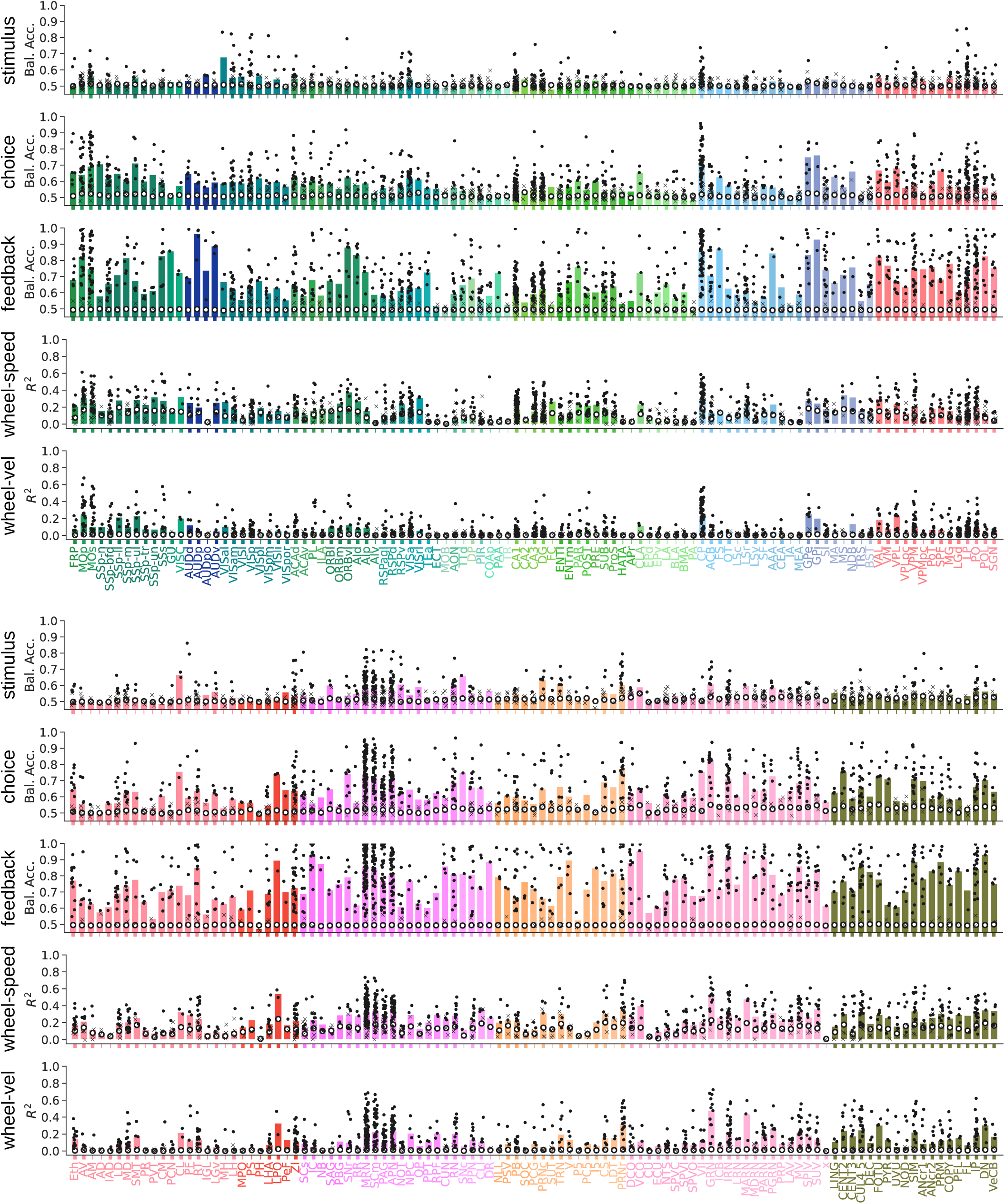
Decoding performance per region with per session results. Decoding analysis as done for stimulus in Fig. 5, choice in Fig. 6, feedback in Fig. 7, and wheel-speed and wheel-velocity in Fig. 8. No FDR correction has been applied in the bar plots, but the bold ticks indicate those regions that survive FDR_0.01_ (and are shown in the main figures). Black dots and x’s indicate decoding performance on individual sessions; dots are significant at *α* = 0.05 and x’s are insignificant. The bar height is the median of all sessions within that region, and the white dot is the across-session median of the null distribution medians.

**Figure S10.**
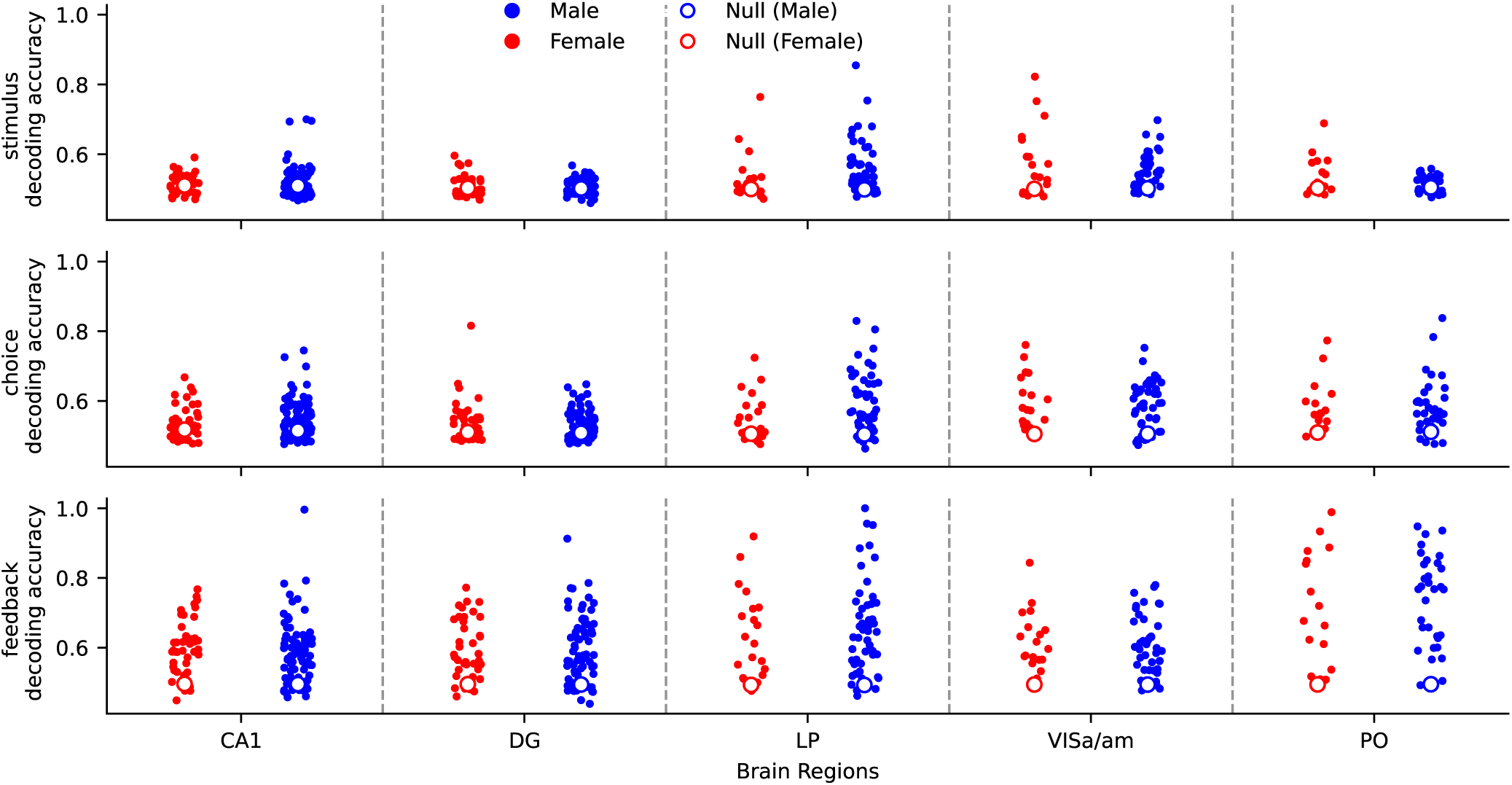
Decoding scores split by sex of mice. For the 5 brain regions of the repeated site and our three main variables, decoding accuracies are shown, split by the sex of the mice. There is no obvious difference between the distributions. The total number of unique animals shown is 104, of which 69 are male.

**Figure S11.**
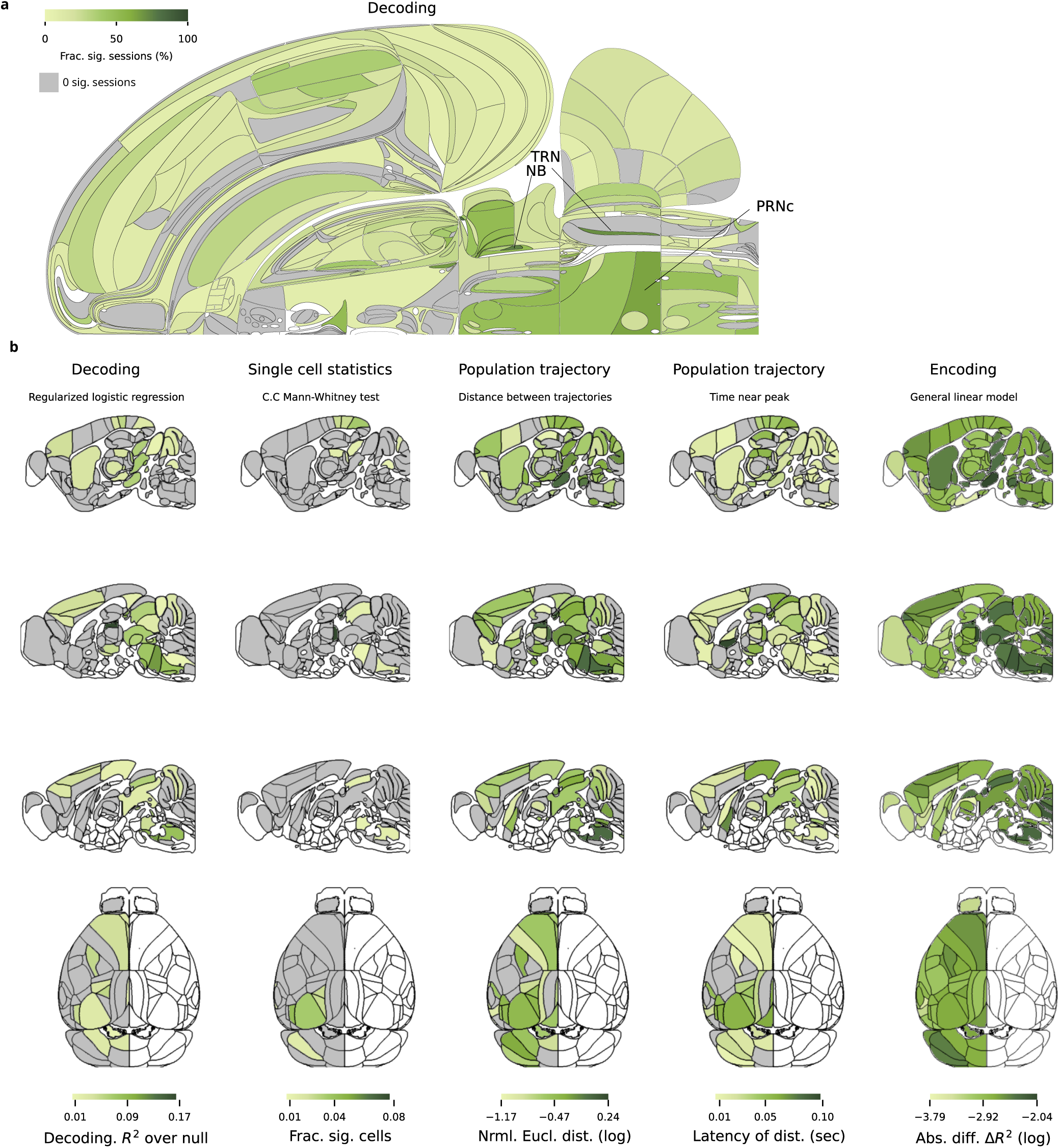
The stimulus variable. **a)** Fraction of sessions with significant decoding performance for the stimulus variable relative to the null. **b)** 2d-brain slices of analysis results for the stimulus variable in Fig. 5a-e. Instead of Swanson flat map, here we use 3 sagittal slices with coordinates ML=-1.8mm, -0.8mm, -0.2mm, and the top view of the dorsal cortex to visualize the representation of task variables across the brain. The locations of sagittal brain slices are optimised to display 252 brain regions. The region acronyms for these slices are listed in Fig. S1.

**Figure S12.**
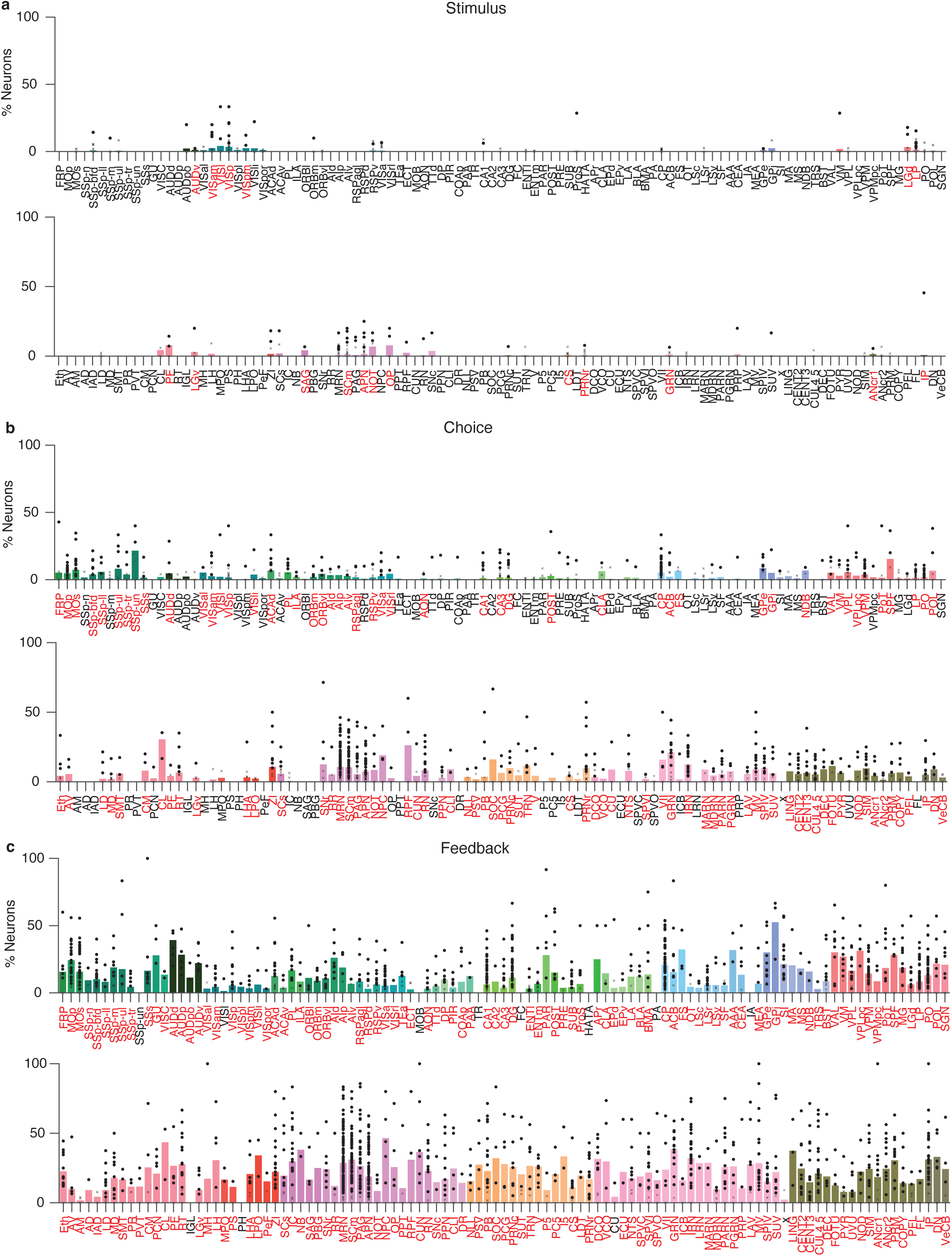
Fraction of significant cells per region in single-cell analysis. Summary of single-cell analysis for stimulus in Fig. 5, **b)** choice in Fig. 6, **c)** feedback in Fig. 7. No FDR correction has been applied in the bar plots; but the red colour labels indicate those regions that survive FDR_0.01_ (and are shown in the figures in the main paper). Black dots and x’s indicate single-cell analysis is done on individual sessions where dots are significant at *α* = 0.05 and x’s are insignificant. The bar height is the mean of all sessions within that region.

**Figure S13.**
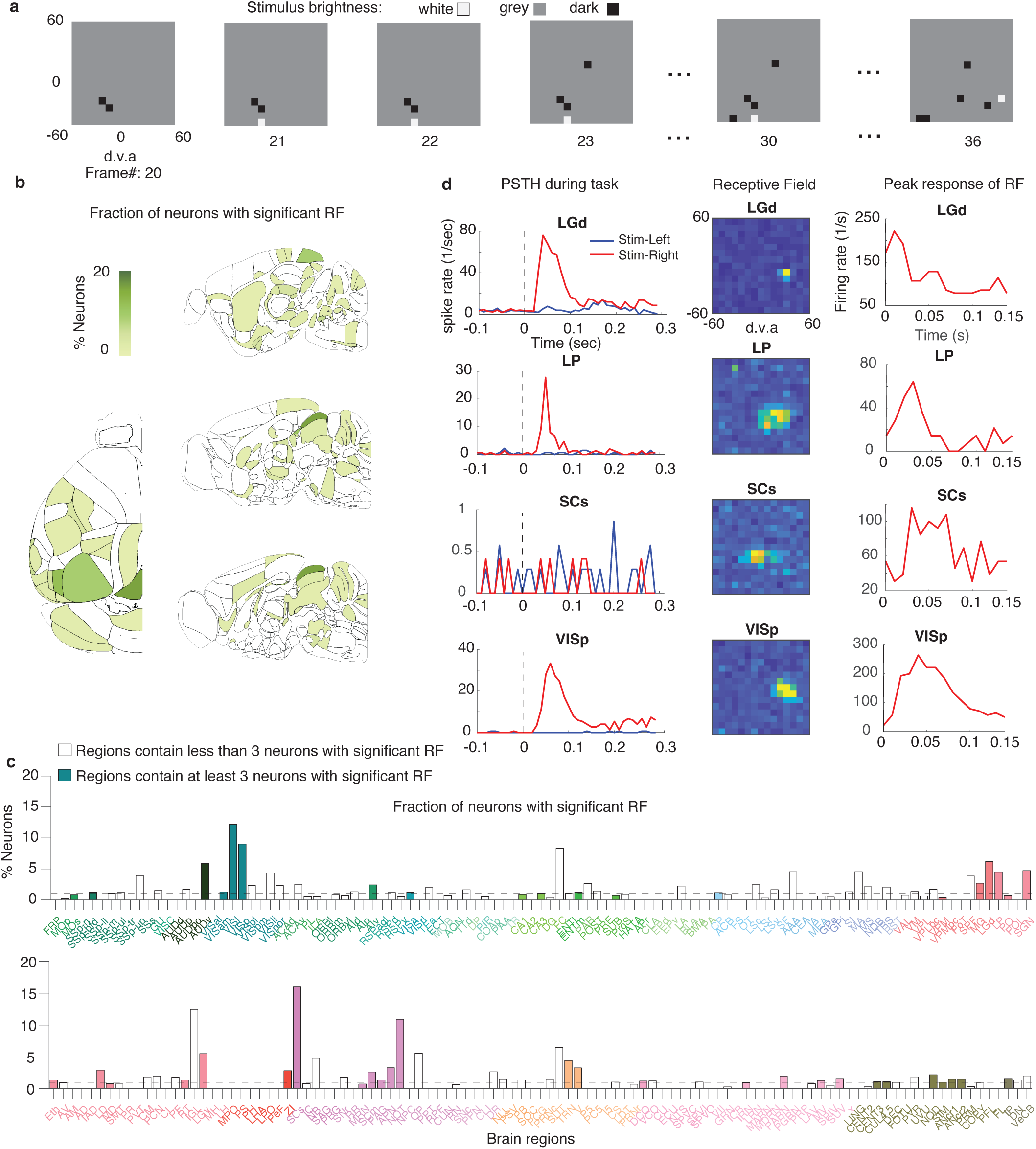
Receptive field mapping of single-cell across the brain. **a)** Examples of the sequence of visual stimulus for receptive field mapping. Frame rate is 60 Hz. White/grey/dark pixels indicate white/grey/dark stimulus, respectively. (d.v.a. stands for degrees of visual angle) **b,c)** Fraction of neurons with significant receptive field in each brain region. In panel **c)**, the hollow bars indicate regions containing fewer than 3 neurons with significant RFs, while the filled bars indicate regions containing at least 3 neurons with significant RFs. **d)** PSTH during the task, the shape of the receptive fields, and the peak response of the receptive fields aligned to stimulus onset for example single cells with a significant receptive field. The peak response of the receptive fields is defined as the PSTH of the pixel in the receptive field with a maximal average spike rate.

**Figure S14.**
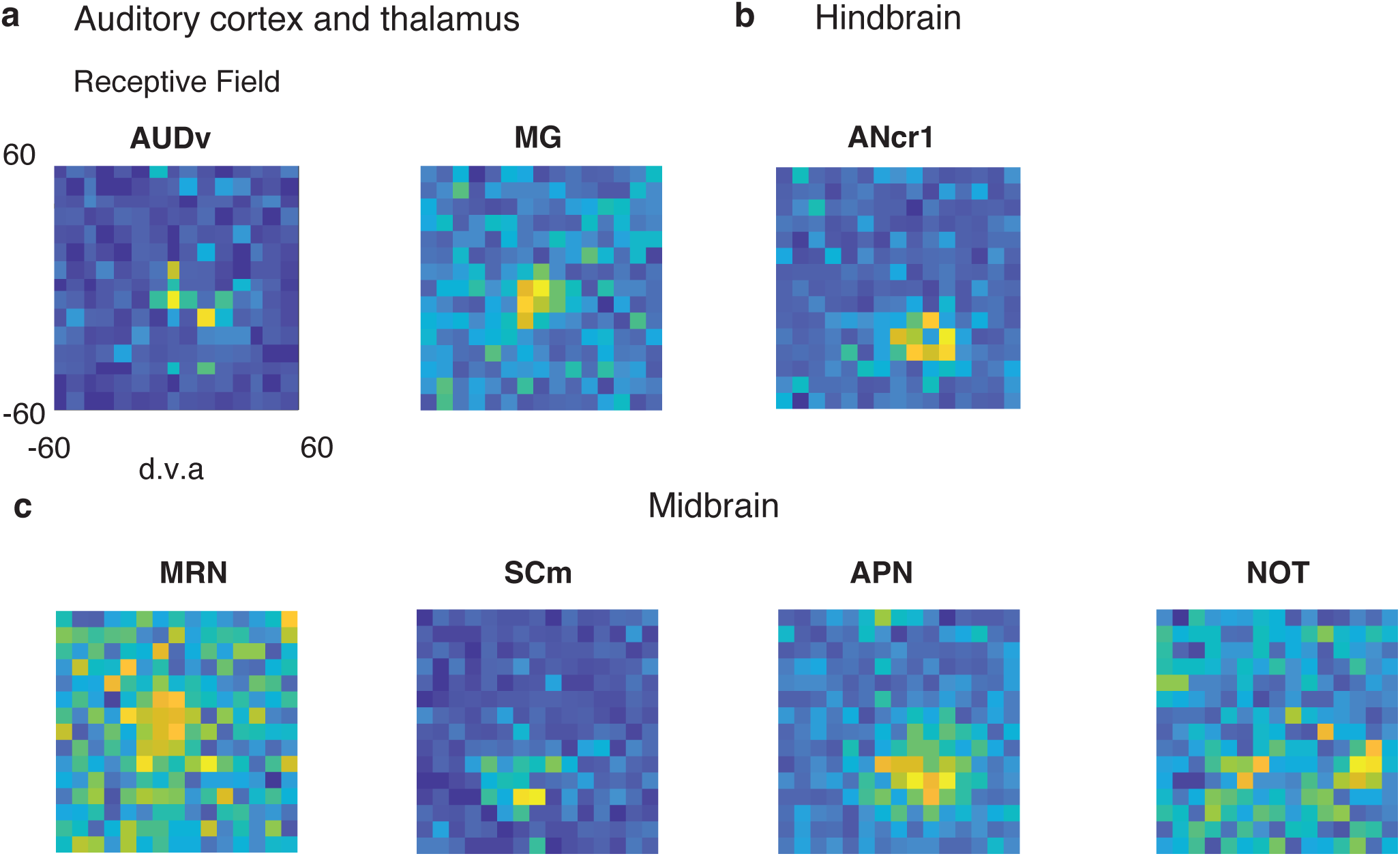
Example of significant receptive fields of single neurons in auditory areas, hindbrain, and midbrain. **a)** Example of receptive fields in auditory cortex (AUDv) and auditory thalamus (MG) (d.v.a. stands for degrees of visual angle). Each pixel in the receptive field denotes 8 *×* 8 d.v.a.. The receptive field is computed by averaging spike rate aligned with On and Off stimulus onset for each pixel, from 0 to 100 ms (Methods). **b)** Example of receptive fields of single neurons in hindbrain. **c)** Example of receptive fields of single neurons in midbrain.

**Figure S15.**
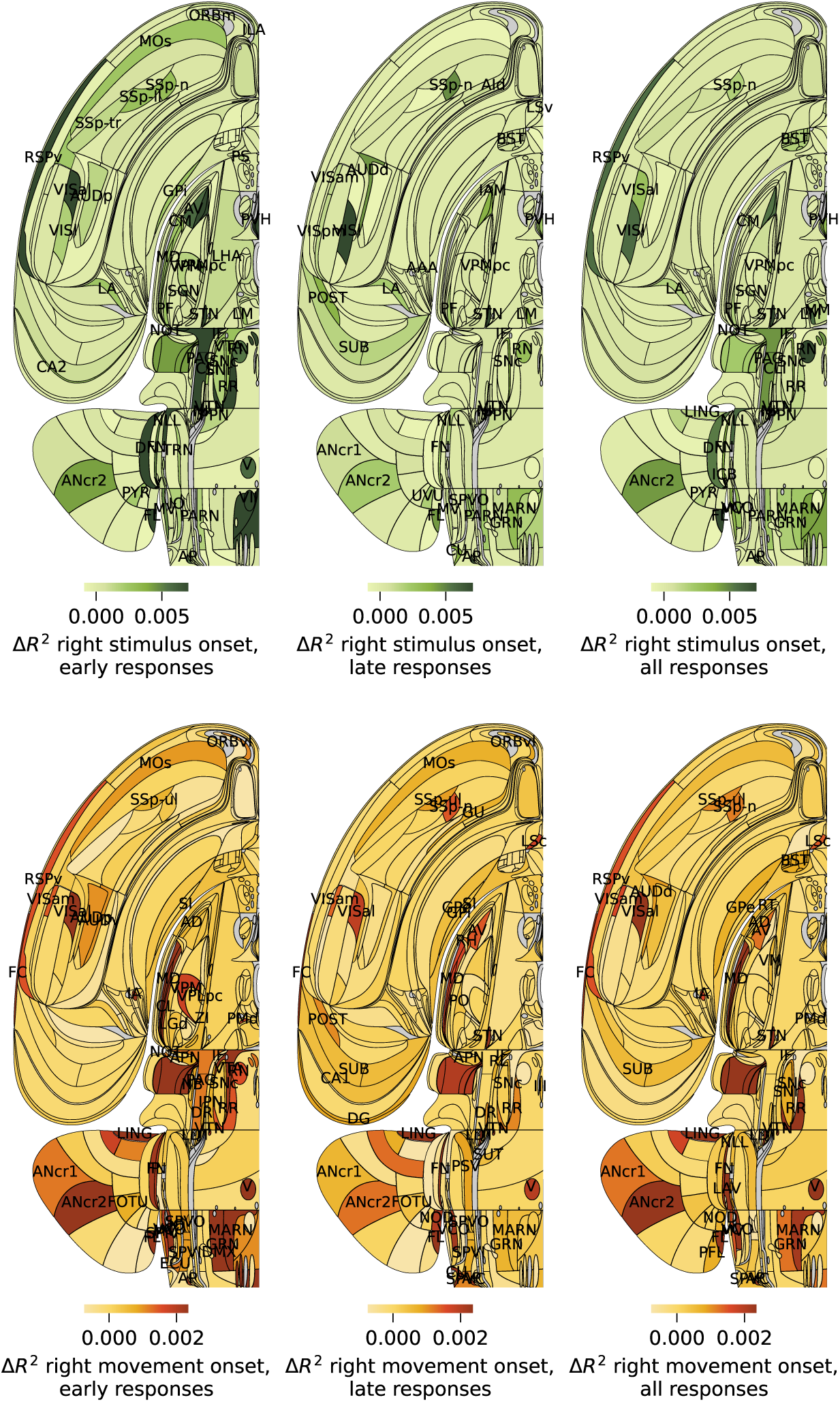
Variance explained by stimulus and choice kernels in GLMs fit to early (below median), late (above median), and all RT trials. **a)** Mean Δ*R*^2^ from right stimulus onset kernel per region in trials with response time below median (left), above median (middle), and all trials (right). **b)** Mean Δ*R*^2^ from right first wheel movement time kernel per region in trials with response time below median (left), above median (middle), and all trials (right).

**Figure S16.**
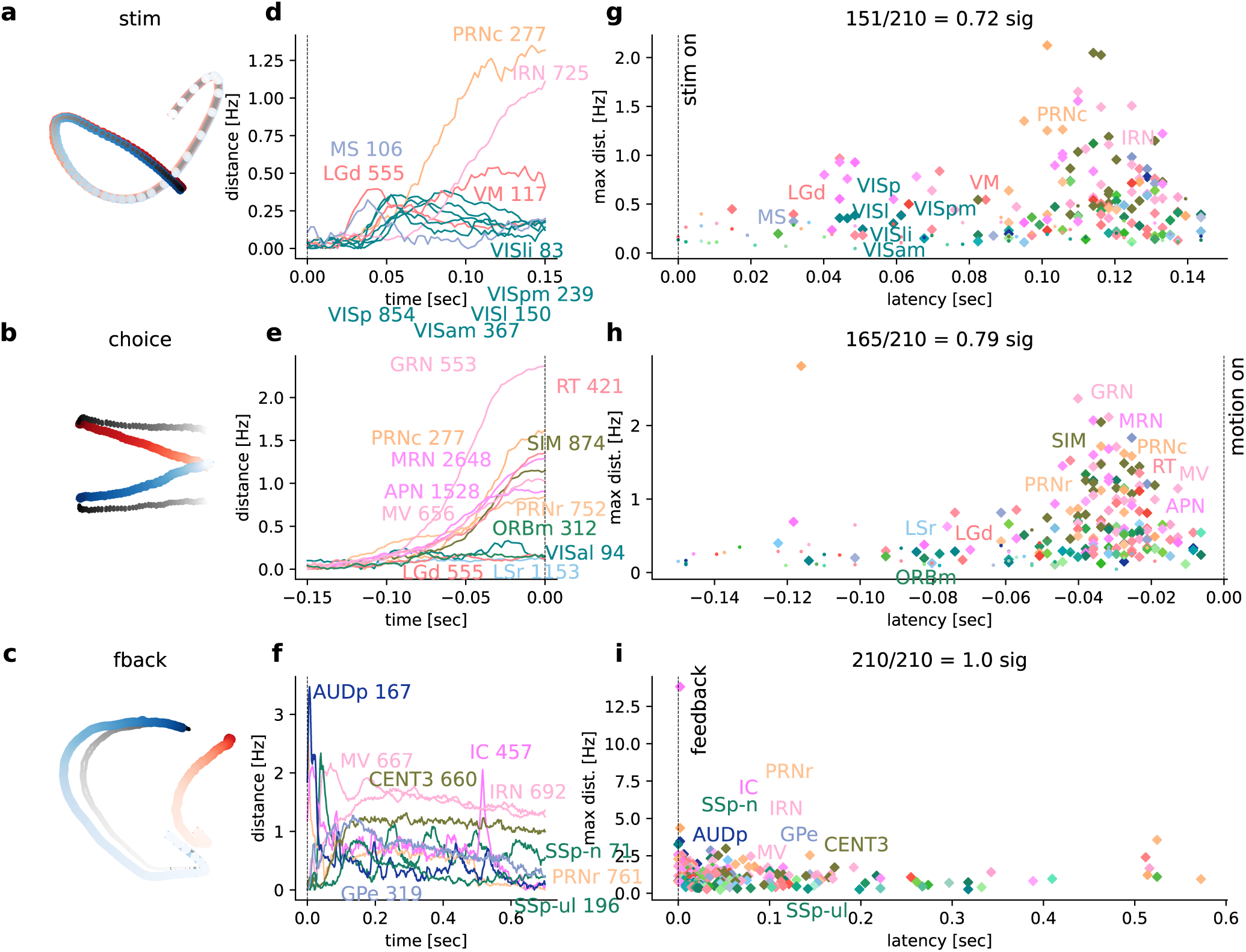
Population dynamics across the brain on the full dataset. Using all good units and considering regions with at least 20 neurons after pooling across sessions, results in about 446 more neurons (in 9 more regions) than in the canonical set of cells that are used across analyses and shown in the main figures. **a-c)** Visualizations (through low-dimensional PCA-embedding) of whole-brain population dynamics (combined across all cells, all sessions, all regions) for three task variables (left versus right stimulus, left versus right choice, correct versus wrong feedback. Blue/red dots represent one time-bin of the population response for left/right (or correct/wrong) trials; colour gradient indicates temporal evolution (darker is later). Grey dots: pseudo-trials. **d-f)** Quantification of the time-resolved distance between opposite trajectories for each variable, based on Euclidean distance (in spikes/second) in the full-dimensional space (dimension = number of cells) for example brain regions, selected based on response magnitude and to illustrate different response profiles. Curves are annotated by region name and number of cells. Scalebars in all panels represent spikes/s/cell. **g-i)** Summary of variable discriminability for stimulus side, choice side, and feedback type, respectively, by magnitude and latency of response across all recorded brain regions. Diamonds indicate all regions that have statistically significant discrimination (*p <* 0.01 relative to pseudo-trial controls), and line plot examples are labelled by region name. Dots indicate responses of non-significant regions.

**Figure S17.**
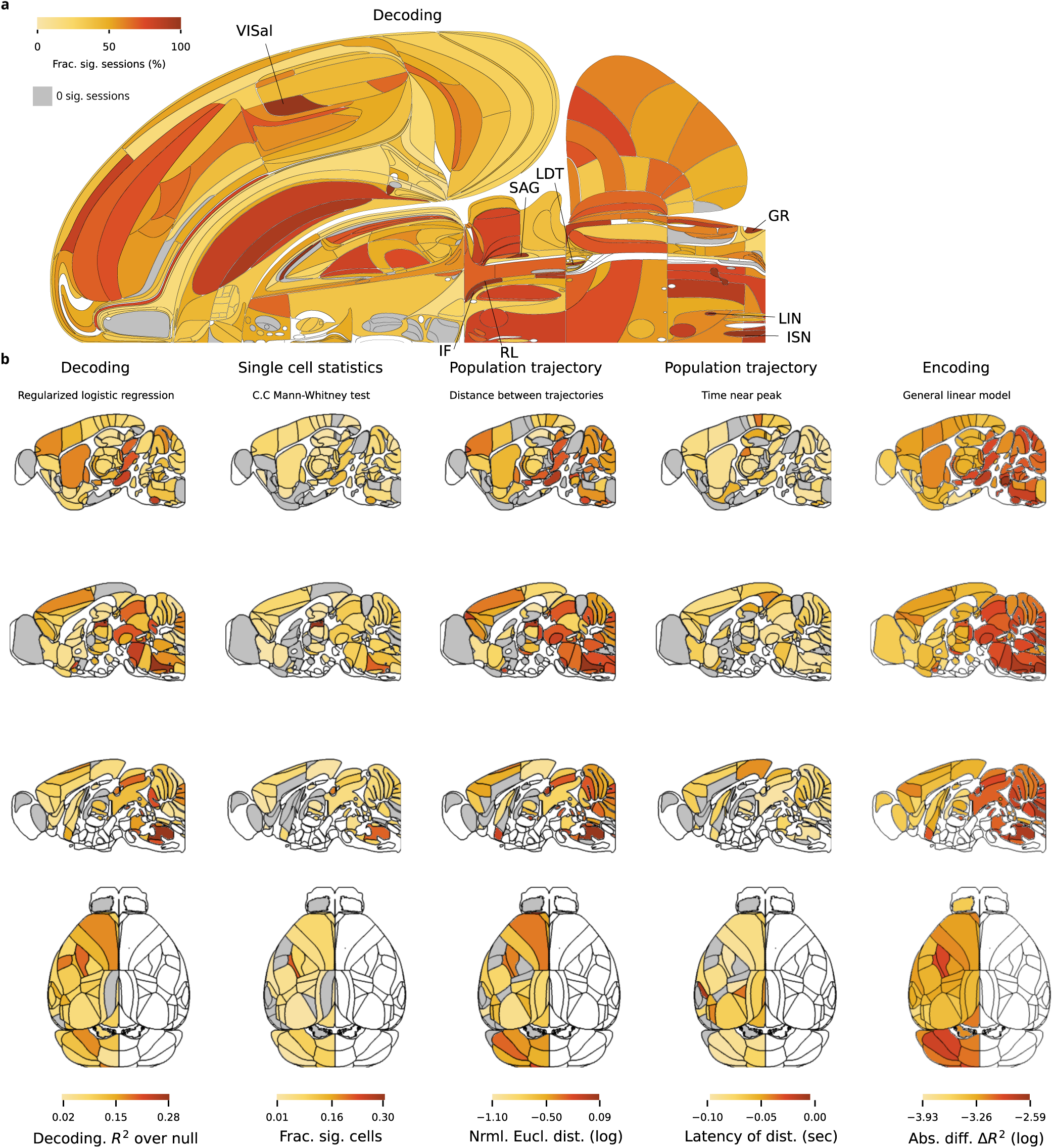
The choice variable. Analysis of the choice variable, with conventions as in Fig. S11.

**Figure S18.**
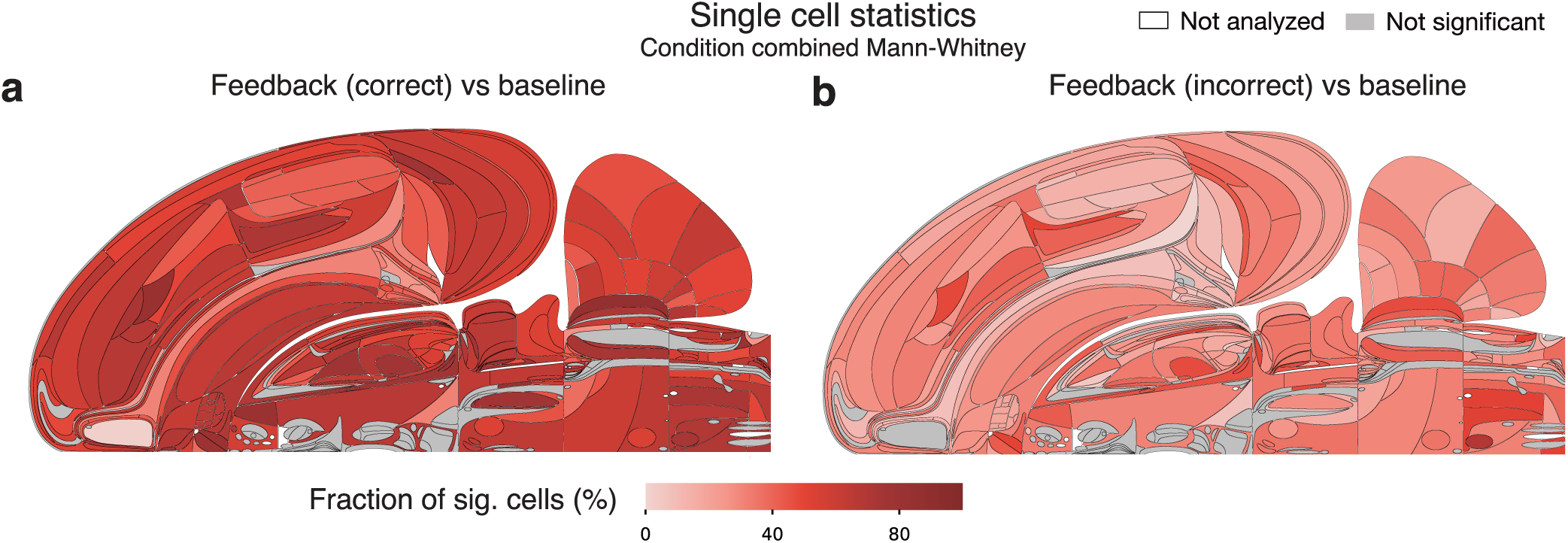
The modulation of neural activity by the feedback signal across the brain relative to baseline. **a)** Fraction of significant neurons per region identified by the condition combined Mann-Whitney test. We compared neural activity after correct feedback ([0, 200] ms) with baseline inter-trial neural activity ([-200, 0] ms aligned to stimulus onset). We deemed a region significant if the number of significant neurons there exceeded the (1 *− α*)th percentile of a binomial (N, *α*) distribution (*α*=0.001), using FDR_0.01_ to correct for multiple comparisons. (Methods). **b)** Comparison between neural activity after incorrect feedback ([0, 200] ms) with baseline inter-trial neural activity ([-200, 0] ms aligned to stimulus onset).

**Figure S19.**
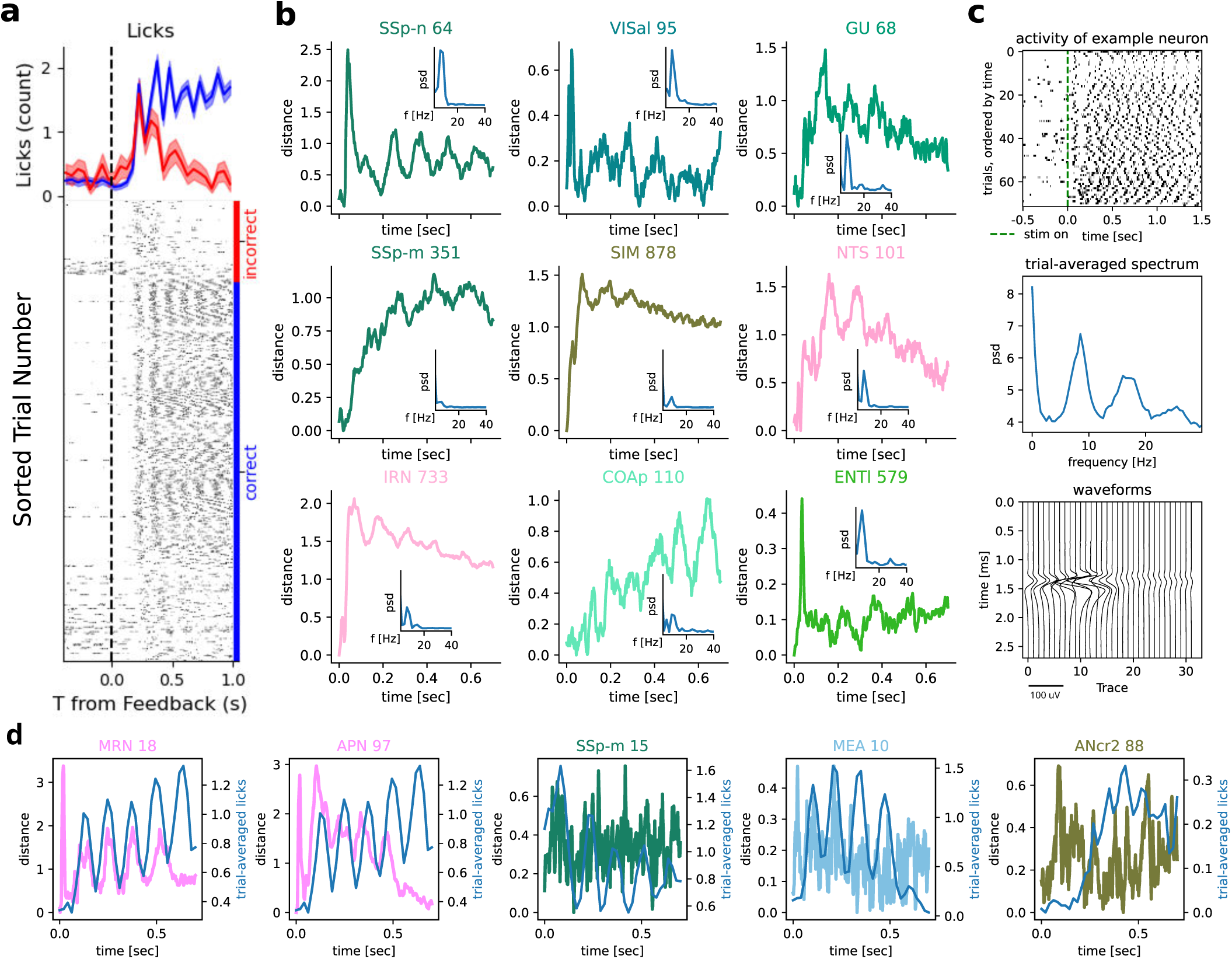
Neural correlates of licking. **a)** Example lick activity for a single session, top trial-averaged, bottom per trial. Animals lick more for correct trials (blue) with a clear rhythm around 10 Hz. Licks were detected using tongue tracking via DLC from side videos. **b)** Population trajectory distance between correct and incorrect trials for example regions selected manually for visible oscillations, with the number of cells (pooled across sessions) next to the region acronym in the title, aligned to feedback. The inset of each panel shows the power spectral density of the distance curve, all having a peak around 10 Hz, correlating with licking. **c)** One example neuron’s activity (pid = ‘3b729602-20d5-4be8-a10e-24bde8fc3092’, region VPL) to show activity is physiological and not an artefact. Top panel, raster per trial with rhythmic 10 Hz activity, also shown in the middle panel by the power spectral density of the raster, averaged across trials. Bottom panel, waveforms of this neuron across adjacent traces, illustrating that the spikes we counted are physiological rather than being caused by an electrical artefact. Artefacts could arise, for example, from current flowing through the drinking spout into the Neuropixels probe, which would result in all traces having a strong waveform. We exclude saturated segments prior to analysis and after this found no evidence for such artefacts when sampling various neurons and inspecting the waveforms. **d)** Single-session population trajectory distance for select regions with trial-averaged lick activity in blue on top. E.g. in MRN a clear correlation with licking was found when restricting the analysis to a single session, while much less so when considering the session-averaged results (not shown).

**Figure S20.**
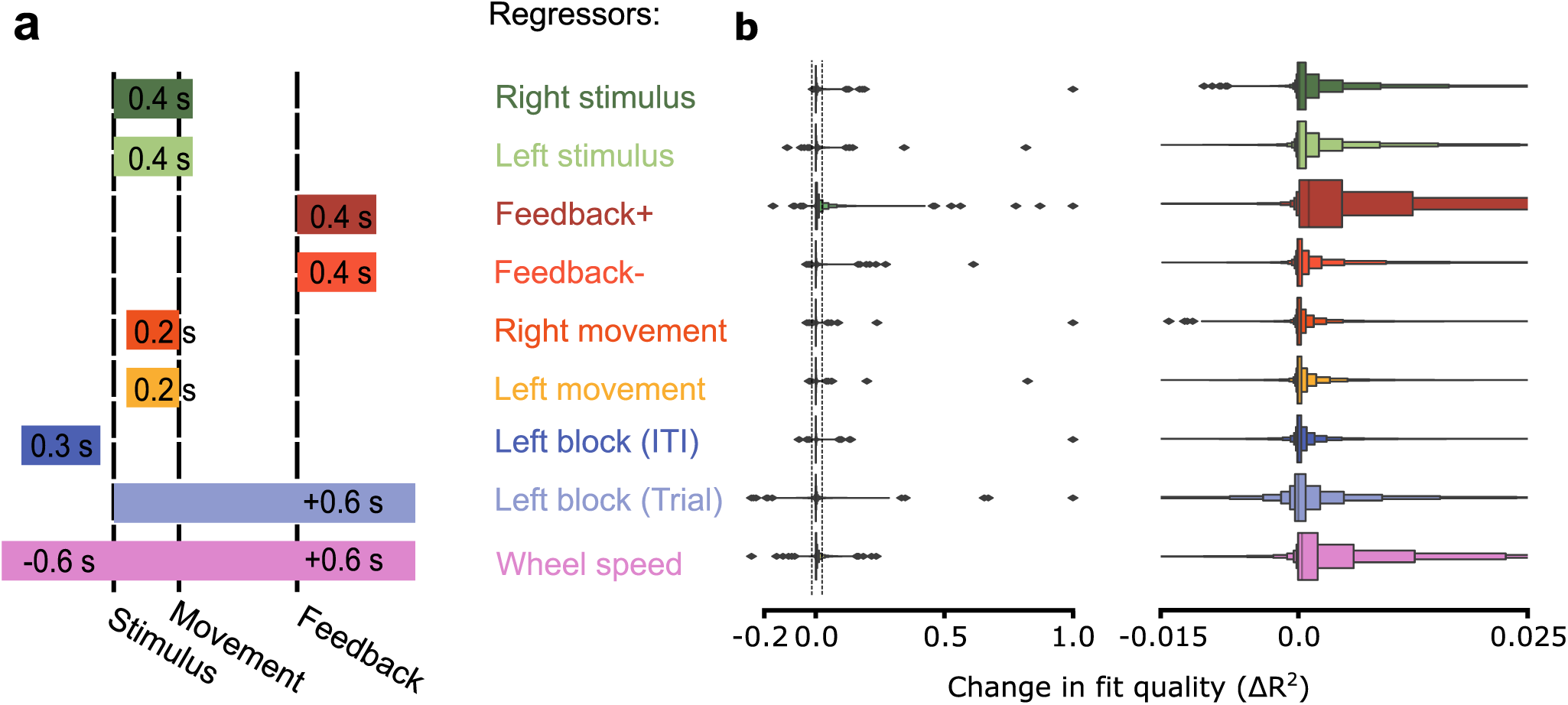
Regressor windows and variance explained in linear encoding model. **a)** Schematic of within-trial windows in which different regressors in the encoding model apply to firing predictions. **b)** Additional variance explained in a leave-one-out paradigm by each regressor for the full distribution (left) and zoomed-in to the medians of the distributions (right). Note that the range on the right panel is depicted on the left via dotted lines.

**Figure S21.**
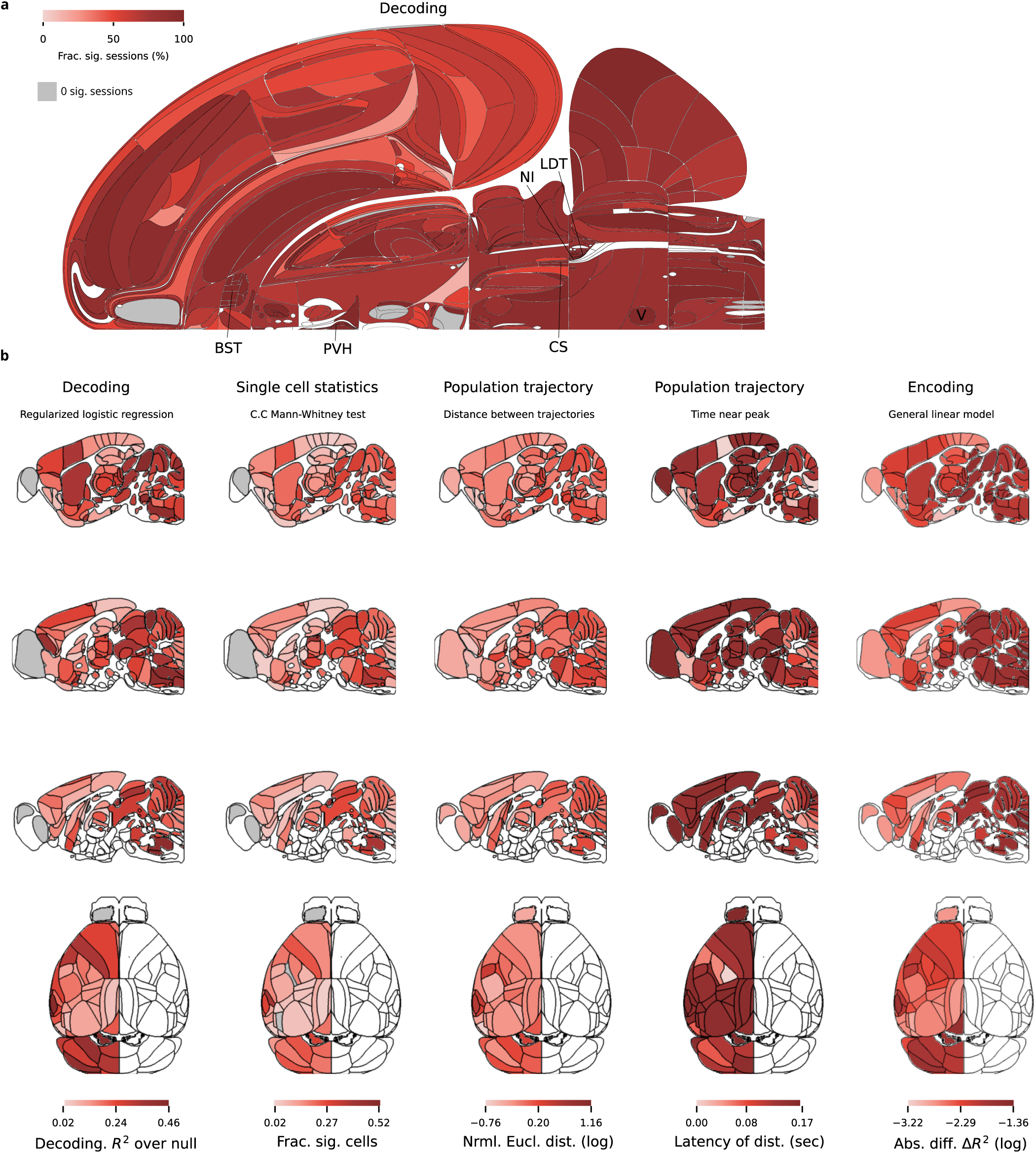
The feedback variable. Analysis of the feedback variable, with conventions as in Fig. S11.

**Figure S22.**
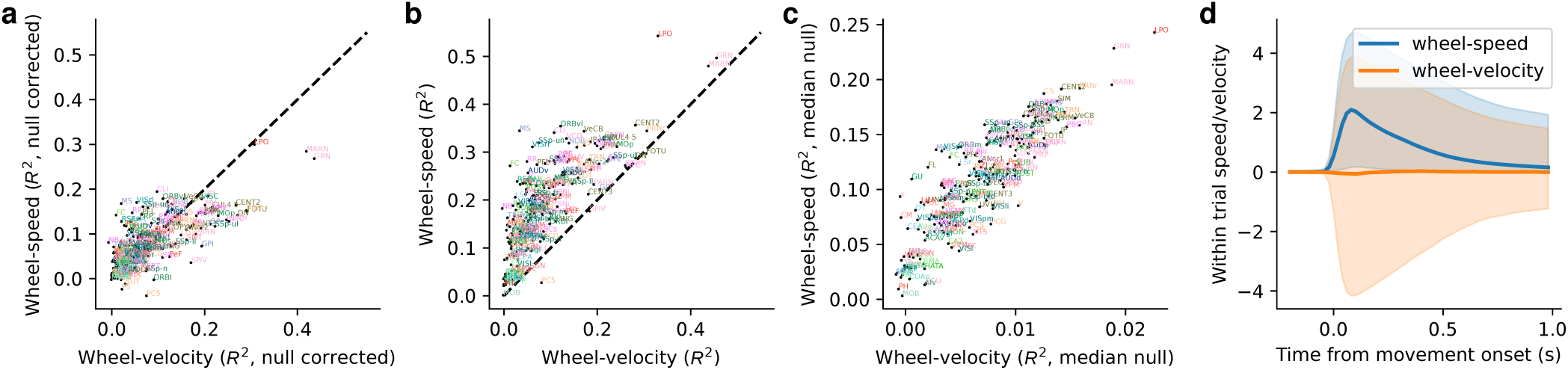
Decoding wheel-velocity versus wheel-speed. Scatter plots comparing per-region wheel-velocity decoding results against wheel-speed for all canonical regions. Decoding is performed for all session-region pairs in the canonical set and the median metric of all such pairs in a given region is plotted. Three such metrics are shown: **(a)** *R*^2^ scores corrected by the median of the null distribution, **(b)** *R*^2^ scores, and **(c)** median of the null distribution. Note the difference in scales for the axes in c. **(d)** The median wheel-speed and wheel-velocity trajectories across all trials are shown and the 5th to 95th percentiles are lightly shaded. The stereotyped shape of wheel-speed produces higher *R*^2^ scores for sessions and null sessions. For example, computing *R*^2^ between 400 randomly chosen trials and 400 repetitions of the median wheel-speed trajectory gives an *R*^2^ of 0.244 (averaged across 1000 repeats). The same computation for wheel-velocity yields *R*^2^ = 0.000.

**Figure S23.**
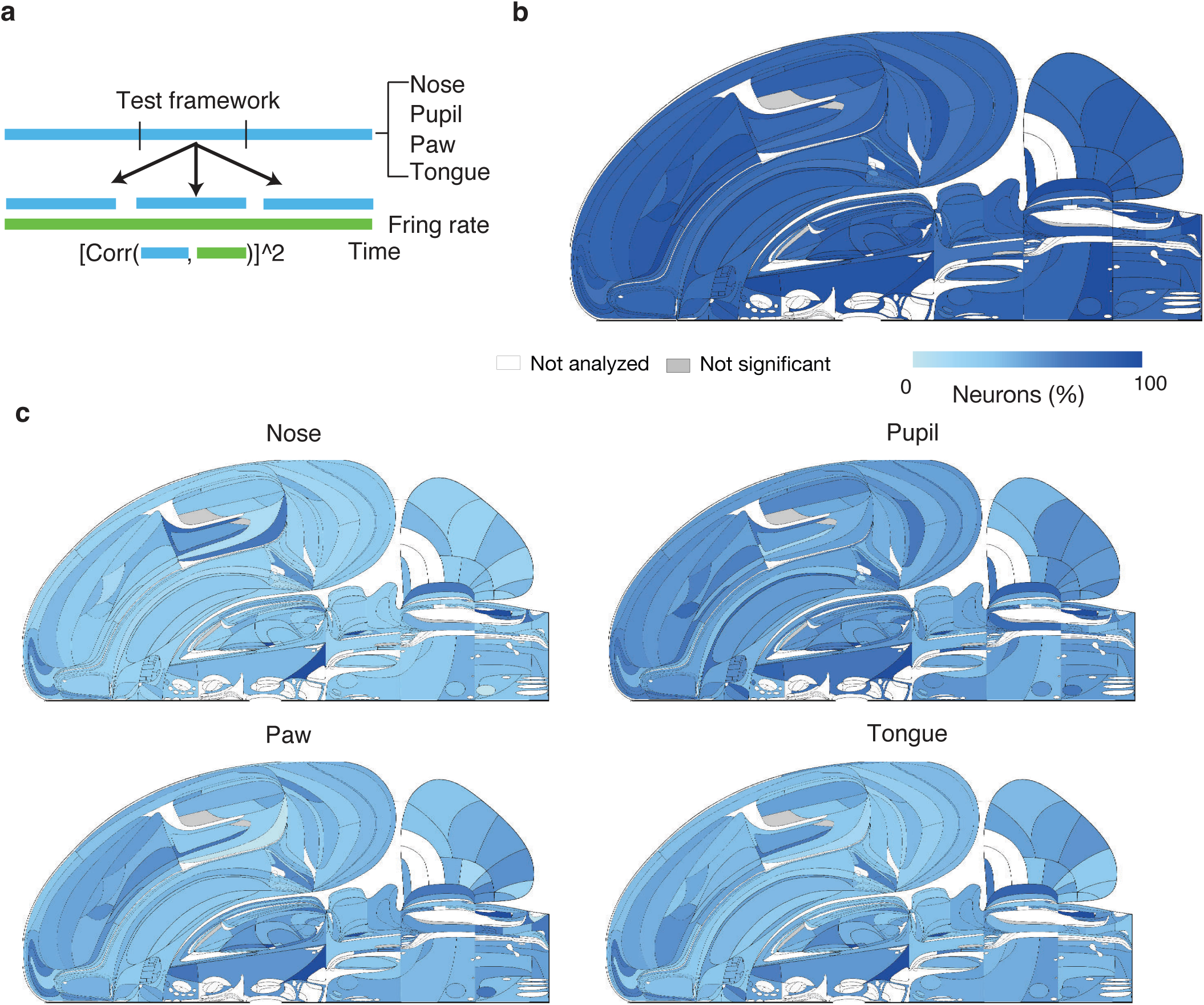
The behavioural correlates of single-neuron activity across the brain. **a)** Statistical tests to measure the behavioural correlates of single neurons across all sessions. We compute the Pearson correlation coefficient between the time series of neural activity and five behavioural variables (nose position, pupil diameter, paw position, and licks, extracted from behaviour video by using DLC; see Methods). The significance of correlation is estimated by a time-shift test^78^ (Methods), using FDR_0.01_ to correct for multiple comparisons. **b)** The flat brain map of the fraction of neurons significantly correlates with at least one of the movement variables. **c)** The flat brain map of the fraction of neurons that significantly correlate with one of the movement variables: noise, pupil, paw, tongue.

**Figure S24.**
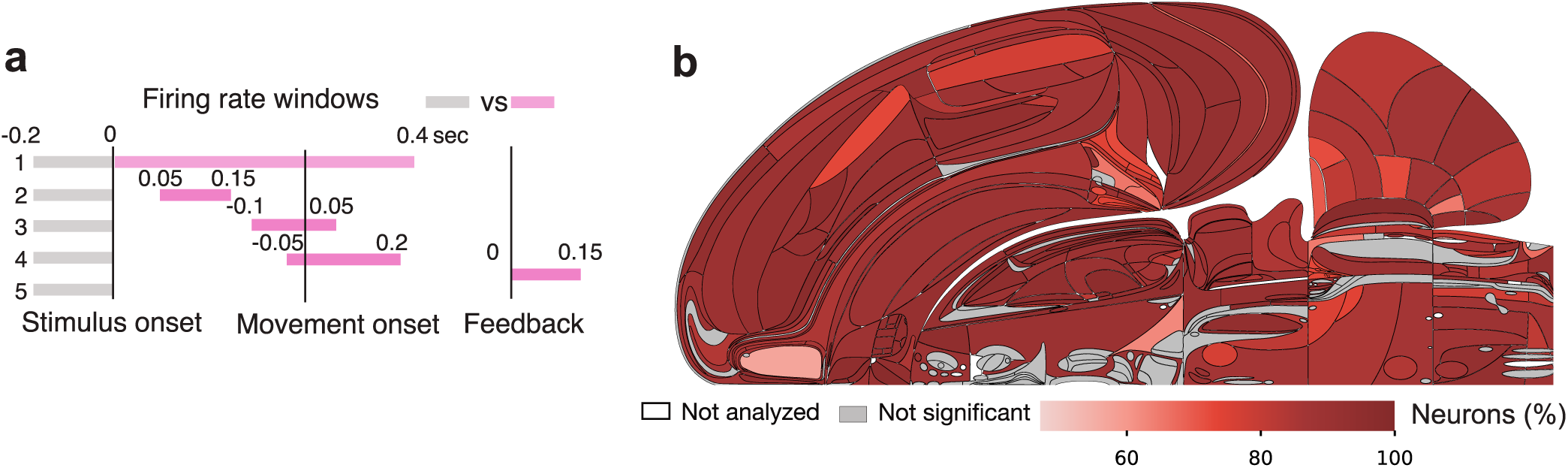
The neural correlates of the task across the brain. **a)** Statistical tests to measure responsiveness in different task windows. The schematics show the summary of all tests, superimposed on the task timeline. Each row represents a separate Wilcoxon rank-sum test comparing firing rates in two different periods over which firing rates were estimated. **b)** The flat brain map of the fraction of neurons that show significant task response during at least one of the task epochs (test of responsiveness: **a**), using FDR_0.01_ to correct for multiple comparisons.

**Figure S25.**
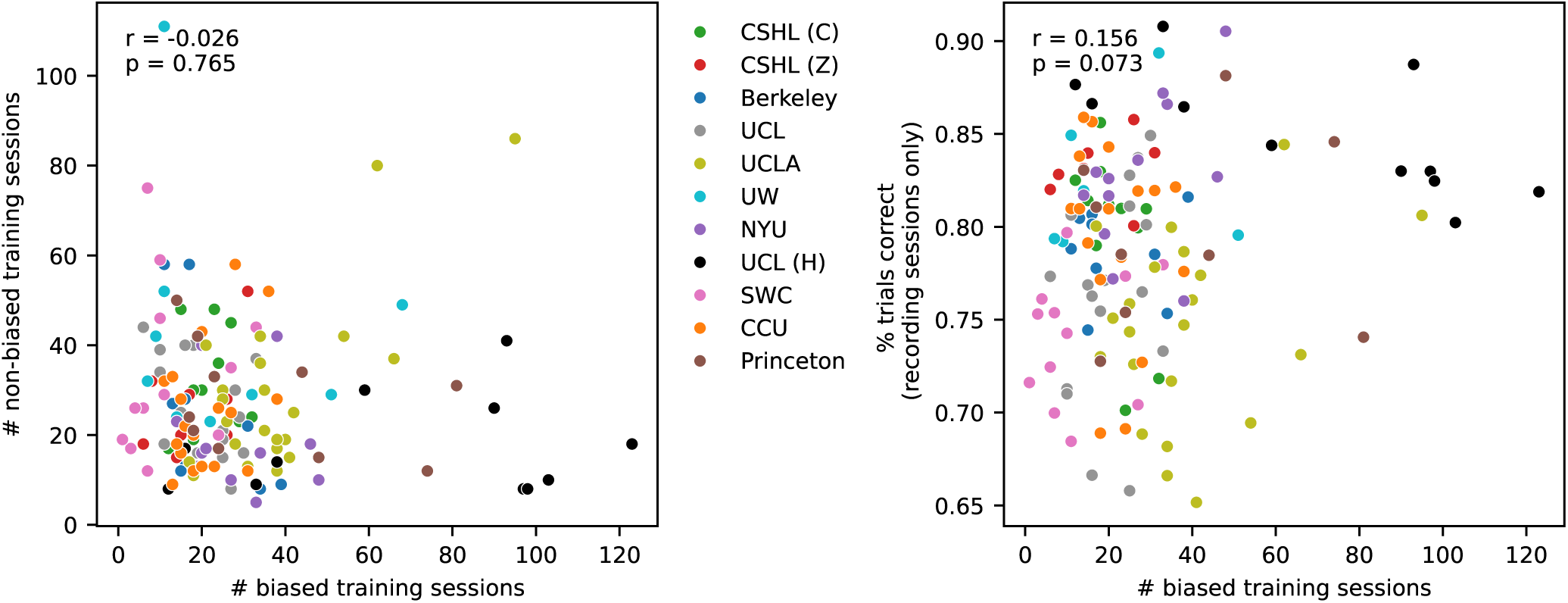
Training and performance statistics for each mouse, colored by lab, and their correlations across animals. **Left:** Scatter-plot of the number of biased versus unbiased training sessions of each animal. There is no significant correlation between these stages of training. **Right:** Scatter-plot of the number of biased training sessions of each mouse, against the overall percentage of correct trials across all recording sessions of that animal. There is again no significant correlation. The correlation between the number of pre-bias sessions and performance during recording is *r* = *−*0.142*, p* = 0.094 (not shown).

**Figure S26.**
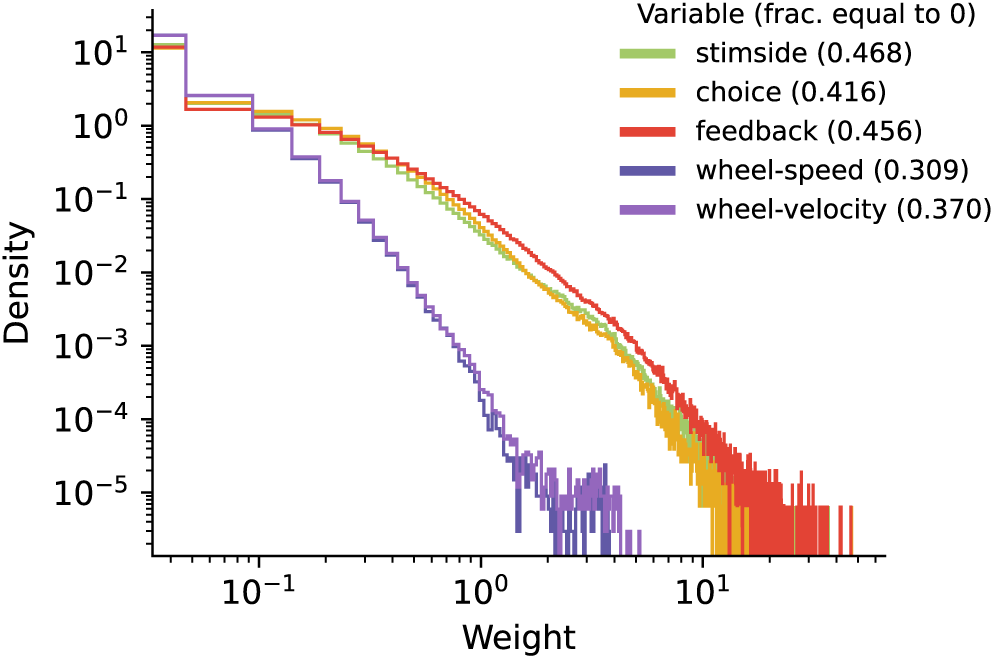
Histogram of regressor weights in decoding analysis. The distributions include decoding weights across all region-session pairs in the canonical set. The distribution combines all weights used on held-out test folds including all temporal bins (for wheel-speed and wheel-velocity) and all repeated decoding runs, but excludes regression intercepts. The legend indicates the decoded variable, and the fraction of weights equal to zero (due to L1 regularization) is shown in parentheses.

**Figure S27.**
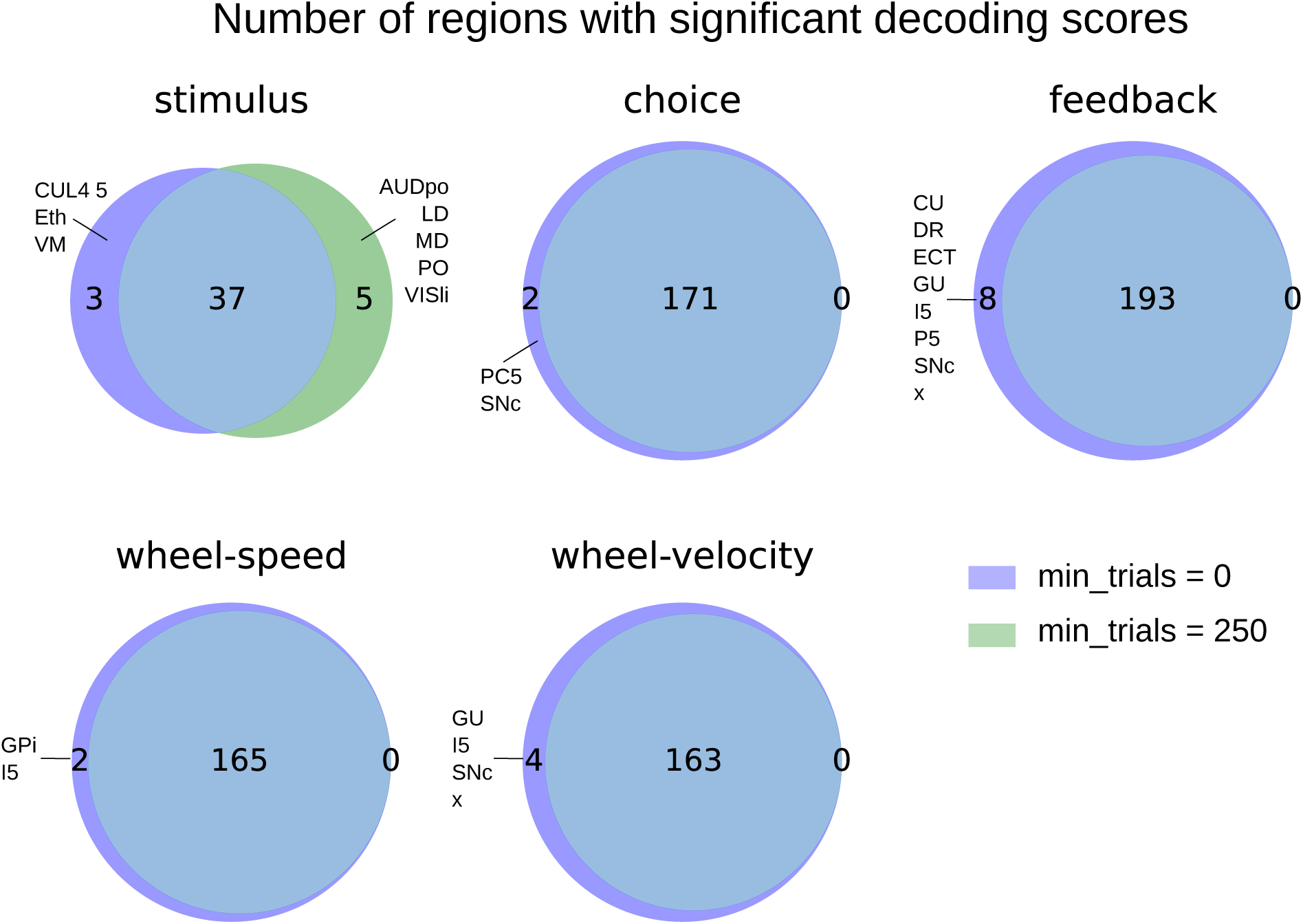
Number of regions with significant decoding scores when including a constraint on the minimum number of trials. The companion prior paper^23^ requires a minimum of 250 trials in order to perform decoding of a given session. We waive that requirement for the decoding analyses in this paper in order to match the same neurons used in the other analyses, which affects the significance of only a small proportion of regions for each target variable.

